# The p53–p21–Cyclin D2 regulatory axis drives metabolic reprogramming and a distinct senescent macrophage senotype during aging and MASLD

**DOI:** 10.64898/2026.09.22.753597

**Authors:** Grasiela Torres, Ivan A. Salladay-Perez, Christina Y. Deng, Itzetl Avila, Yennifer Delgado, Gregory Tong, Jesús Muñoz-Estrada, Lizeth Estrada, Anika Dhingra, Wesley R. Armstrong, Aishini Singh, Isabella Cooper, Eduardo Enriquez, Prashant Kaushal, Jocelyn Ni, Kushan Chowdhury, Calvin Pan, Linsey Stiles, Cristiane Beninca, Simon T. Hui, Abbigail S. Krall, Heather R. Christofk, Aldons J. Lusis, Timothy E. O’Sullivan, Jesse G. Meyer, Rajat Singh, Valerie A. Tornini, Orian S. Shirihai, Mehdi Bouhaddou, Anthony J. Covarrubias

**Author notes:** Contributed Equally (Ivan A. Salladay-Perez, Christina Y. Deng). **Correspondence** to Anthony J. Covarrubias.

## Abstract

Aging drives chronic disease in part through senescent cells, including macrophages, which fuel inflammation. Senescent macrophages are functionally heterogeneous: canonical p16-high macrophages promote tumorigenesis or, in other contexts, disease tolerance, whereas we previously identified a distinct p21-high, p16-low senotype that drives metabolic dysfunction-associated steatotic liver disease (MASLD). The molecular basis of this senotype has remained undefined. Using genetic and multi-omic approaches, we show that a p53-p21-dependent program actively represses p16 and is required for senescent macrophage viability. We identify Cyclin D2 as a non-canonical downstream effector that redistributes from the nucleus to mitochondria and lipid droplets, where it partners with MIC60 to drive metabolic reprogramming and modulates AKT1-mTORC1 signaling that sustains the SASP. Cyclin D2-p21 double-positive macrophages accumulate with aging and MASLD in mice and in human liver cirrhosis, and can be selectively depleted by senolytic treatment. Together, these findings define a druggable p53-p21-Cyclin D2 axis that specifies macrophage senotype.

**Highlights:**

● A p53–p21-dependent program drives macrophage senescence while actively repressing p16
● p16 repression promotes cell survival during acute genotoxic stress, but is dispensable for cell-cycle arrest and core senescence features in macrophages
● Cyclin D2 is a non-canonical p53–p21 effector that relocalizes from nucleus to the cytosol and mitochondria
● Cyclin D2 engages the AKT1-mTORC1 pathway and interacts with MIC60 to drive metabolic reprogramming
● Cyclin D2+p21+ macrophages are abundant in aging, MASLD, and human liver cirrhosis, and can be targeted by senolytics

## Introduction

Cells respond to DNA damage by activating distinct cell fate programs downstream of the DNA damage response (DDR), including programmed cell death (apoptosis) and cellular senescence ^1^. Cellular senescence is a state of irreversible cell-cycle arrest accompanied by extensive phenotypic remodeling, including metabolic reprogramming, resistance to apoptosis, and secretion of pro-inflammatory factors collectively termed the senescence-associated secretory phenotype (SASP) ^2^. Although initially recognized as a consequence of replicative exhaustion, senescence can also be induced by diverse physiological and pathological stimuli, including DNA damage, lipid and cholesterol accumulation, chronic infection, and mitochondrial dysfunction^3,4^. Senescence can serve beneficial functions in development, tissue remodeling, and tumor suppression, but the accumulation of senescent cells with age can promote chronic inflammation and contribute to disease.

Macrophages are increasingly recognized as an important component of the senescent cell burden in aging and chronic disease, where their persistence and inflammatory activity can influence tissue dysfunction^5^. We previously demonstrated that p21^+^TREM2^+^ senescent macrophages accumulate with age and contribute to the pathogenesis of metabolic dysfunction-associated steatotic liver disease (MASLD) in mice and liver cirrhosis in humans^6^. However, the molecular circuitry that establishes and maintains macrophage senescence remains unclear.

Molecular mechanisms engaged during senescent programming are highly cell type and stimulus-dependent, underscoring the heterogeneity of senescent cell populations. Despite this complexity, senescent cells are commonly identified by increased senescence-associated β-galactosidase (SA-β-gal) activity and by upregulation of canonical cell-cycle inhibitors such as p21 (*Cdkn1a*) and p16 (*Cdkn2a*). Emerging evidence indicates that these markers are not simply associated with a single, interchangeable senescent state but instead define their specific ‘senotypes’, which are distinct senescence programs shaped by cell type, stimulus and tissue context with sometimes opposing, functional consequences^7^. This is particularly evident in macrophages: p16-high macrophages accumulate in the aging and tumor-bearing lung and promote early-stage tumorigenesis^8,9^, whereas a separate p16-high macrophage population has recently been shown to enforce disease tolerance and limit inflammatory tissue damage during infection and sepsis^10^. This apparent paradox, the same canonical marker demarcating both a pro-tumorigenic and a protective macrophage state, illustrates that p16 expression alone cannot predict the functional consequences of macrophage senescence. We recently described a third, molecularly distinct senotype: p21-high, p16-low senescent macrophages that accumulate in aging tissues and drive MASLD^6^. Thus, the question of what pro-senescent molecular pathways help drive a distinct p16- or p21-dominant senotype, and whether these programs are mechanistically distinct drivers of divergent disease outcomes remains unclear. Resolving this question requires mechanistic insight into the pathways that establish and maintain each senotype.

Here, we address this question by delineating the molecular mechanisms that establish the p21-high, p16-low macrophage senotype in response to DNA damage. We identify a p53–p21-dependent program that promotes senescent macrophage survival while actively repressing *Cdkn2a* (p16), demonstrating that p16 is not only dispensable but actively antagonized in this context. We further identify Cyclin D2 as a critical downstream effector of the p53–p21 axis that undergoes cytoplasmic relocalization and promotes mitochondrial and metabolic remodeling in senescent macrophages, revealing a non-canonical function for a canonical cell-cycle regulator beyond proliferation control. Together, our findings define a p53–p21–Cyclin D2 regulatory axis that specifies a distinct macrophage senotype by coordinating senescent cell survival with mitochondrial, metabolic, and secretory remodeling, providing a mechanistic framework for understanding how divergent senescence programs arise within the same cell type and may inform the development of senotype-targeted therapeutic strategies.

## Results

### p53–p21 signaling governs senescent macrophage viability and suppresses p16 expression

Our identification of a p21-high, p16-low senotype in senescent macrophages raised the question of how this distinct pattern of cell-cycle inhibitor expression is established. Because p21 (*Cdkn1a*) is a canonical transcriptional target of p53 (*Trp53*), we hypothesized that p53–p21 signaling may play a central role in specifying this senescent macrophage state. To test this, bone marrow-derived macrophages (BMDMs) isolated from wild-type (WT) and *Trp53*^−/−^ (p53 KO) mice were subjected to our established *in vitro* model of macrophage senescence, in which cells are exposed to 10 Gy irradiation (IR) and allowed to undergo senescence (Sen(IR)) for 10 days (Fig. 1A). Western blot analysis revealed increased accumulation of γH2AX in p53 KO BMDMs beginning at 2 days post-IR, indicating greater DNA damage following irradiation (Fig. 1B). Notably, cleaved caspase-3, a marker of end-stage apoptosis, was also elevated in p53 KO BMDMs beginning at this early 2-day time point (Fig. 1B). To determine whether this early increase in apoptotic signaling was accompanied by increased cell death, we performed Annexin V/Propidium iodide (PI) staining at 2 days post-IR to quantify cell death via apoptosis and necrosis, respectively. Loss of p53 resulted in an approximately 2.2-fold increase in total cell death compared with WT BMDMs two days post-IR (Fig. 1C,D). Together, these findings indicate that p53 is required early in the macrophage response to genotoxic stress to promote the DNA damage response (DDR) and survival during the subsequent senescence program.

**Figure 1.**
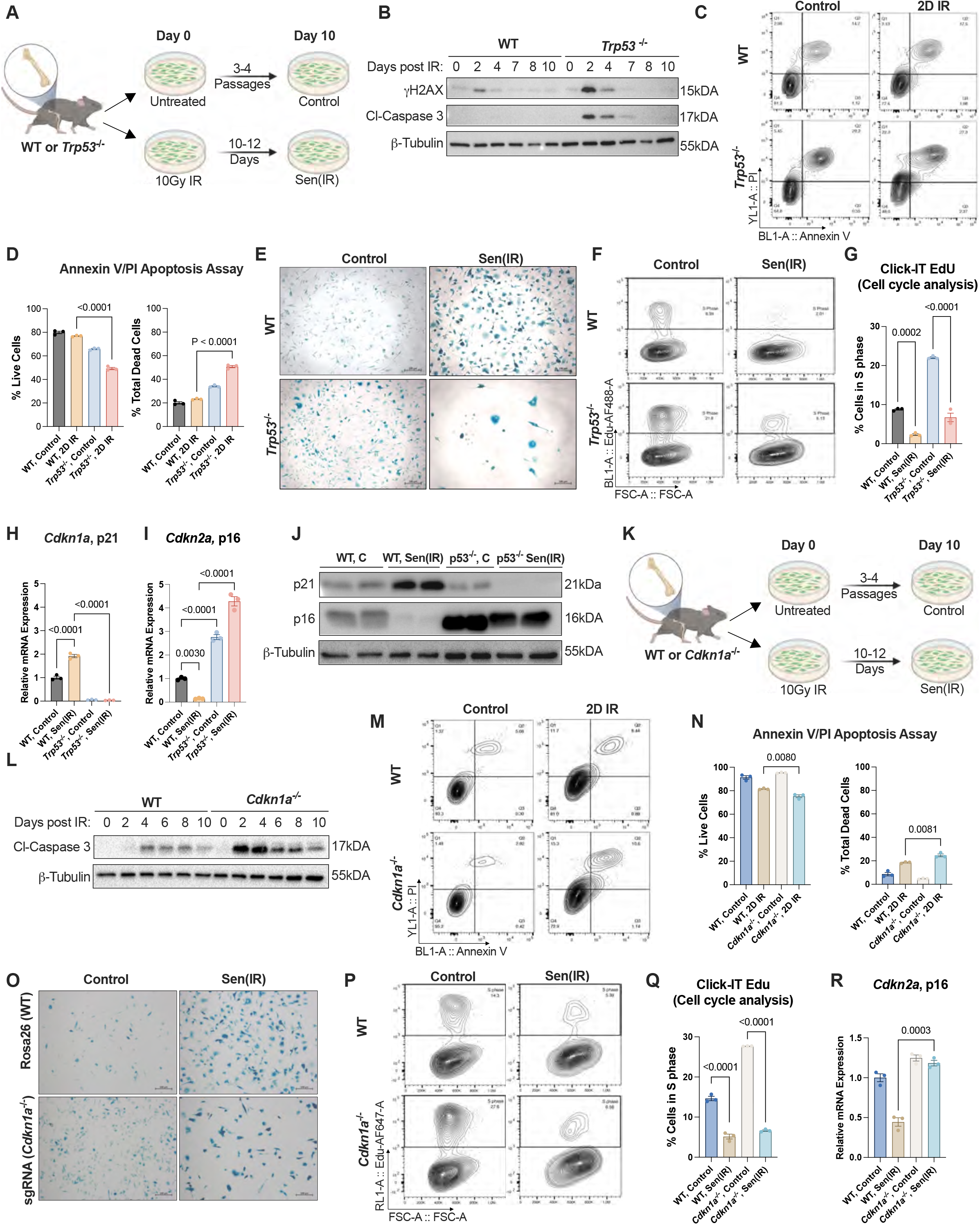
The p53–p21 axis promotes macrophage survival and negatively regulates p16 expression in response to genotoxic stress. (A) Experimental schematic for induction of irradiation-induced senescence in bone marrow-derived macrophages (BMDMs). BMDMs from WT or *Trp53*^⁻/⁻^ mice were either left untreated or exposed to 10 Gy ionizing radiation (IR). Cells analyzed 2 days following irradiation are designated 2D IR, whereas cells analyzed 10-12 days following irradiation are designated senescent (Sen(IR)) macrophages. Created with BioRender.com. (B) SDS-PAGE gels and immunostaining (western blot) for γH2AX and cleaved caspase-3 in WT and *Trp53*^⁻/⁻^ BMDMs at the indicated days following irradiation. Β-Tubulin was used as a loading control. (C) Representative flow cytometry plots of Annexin V and propidium iodide (PI) staining in control and 2D IR WT and *Trp53*^⁻/⁻^ BMDMs. (D) Mean ± s.e.m. percentage of live (Q4) and total dead cells (Q1-3) from Annexin V/PI staining. *P* value of unpaired two-tailed Student’s *t*-test of *n* = 3 biological replicates. I Representative images of senescence-associated β-galactosidase (SA-β-gal) staining in control and Sen(IR) WT and *Trp53*^⁻/⁻^ BMDMs. (F) Representative flow cytometry plots of EdU incorporation in control and Sen(IR) WT and *Trp53*^⁻/⁻^ BMDMs. (G) Mean ± s.e.m. percentage of cells in S phase based on EdU incorporation. *P* value of one-way ANOVA with Tukey’s multiple-comparisons test of *n* = 3 biological replicates. (H) Mean ± s.e.m. *Cdkn1a* (p21) mRNA transcript levels, relative to control, in WT and *Trp53*^⁻/⁻^ BMDMs following senescence induction. *P* value of one-way ANOVA with Tukey’s multiple-comparisons test of *n* = 3 biological replicates. (I) Mean ± s.e.m. *Cdkn2a* (p16) mRNA transcript levels, relative to control, in WT and *Trp53*^⁻/⁻^ BMDMs following senescence induction. *P* value of one-way ANOVA with Tukey’s multiple-comparisons test of *n* = 3 biological replicates. (J) SDS-PAGE gels and immunostaining (western blot) for p21 and p16 in control and Sen(IR) WT and *Trp53*^⁻/⁻^ BMDMs. Β-Tubulin was used as a loading control. (K) Experimental schematic for induction of irradiation-induced senescence in BMDMs from WT or *Cdkn1a*^⁻/⁻^ mice. Created with BioRender.com. (L) SDS-PAGE gels and immunostaining (western blot) for cleaved caspase-3 in WT and *Cdkn1a*^⁻/⁻^ BMDMs at the indicated days following irradiation. Β-Tubulin was used as a loading control. (M) Representative flow cytometry plots of Annexin V and PI staining in control and 2D IR WT and *Cdkn1a*^⁻/⁻^ BMDMs. (N) Mean ± s.e.m. percentage of live (Q4) and total dead cells (Q1-3) from Annexin V/PI staining. *P* value of unpaired two-tailed Student’s *t*-test of *n* = 3 biological replicates. (O) Representative images of SA-β-gal staining in control and Sen(IR) Rosa26(WT) and sgRNA(*Cdkn1a*^⁻/⁻^) macrophages. (P) Representative flow cytometry plots of EdU incorporation in control and Sen(IR) WT and *Cdkn1a*^⁻/⁻^ BMDMs. (Q) Mean ± s.e.m. percentage of cells in S phase based on EdU incorporation. *P* value of one-way ANOVA with Tukey’s multiple-comparisons test of *n* = 3 biological replicates. I Mean ± s.e.m. *Cdkn2a* (p16) mRNA transcript levels, relative to control, in WT and *Cdkn1a*^⁻/⁻^ BMDMs following senescence induction. *P* value of unpaired two-tailed Student’s *t*-test of *n* = 3 biological replicates.

Having established that loss of p53 compromises macrophage survival following genotoxic stress, we next asked whether p53 deficiency also altered other features of the senescence phenotype. Despite an approximately 82% reduction in live p53 KO Sen(IR) BMDMs compared with WT Sen(IR) BMDMs 10 days post-IR, the surviving p53 KO Sen(IR) BMDMs remained SA-β-gal positive (Supp. Fig. 1A, Fig. 1E). Consistent with these findings, SPiDER-gal analysis confirmed detectable β-gal activity in p53 KO Sen(IR) BMDMs; however, SA-β-gal-associated fluorescence was significantly reduced compared with WT Sen(IR) cells (Supp. Fig. 1B). Thus, although p53-deficient macrophages exhibit a marked loss of viability following IR, the surviving cells retain features of the senescence-associated lysosomal phenotype.

Notably, the surviving p53 KO Sen(IR) cells retained the cell cycle arrest characteristic of senescent macrophages, exhibiting an approximately 74% reduction in S-phase entry following IR compared with p53 KO non-senescent control cells, as measured by Click-iT EdU incorporation (Fig. 1F,G). Consistent with loss of p53, *Cdkn1a* (p21) expression was completely ablated in p53 KO Sen(IR) BMDMs (Fig. 1H). Strikingly, *Cdkn2a* (p16) expression was restored and markedly elevated in p53 KO Sen(IR) BMDMs relative to WT Sen(IR) cells, revealing an unexpected negative regulatory role for p53 in controlling p16 expression during macrophage senescence (Fig. 1I). This finding was further supported by western blot analysis, which demonstrated increased p16 protein levels in both untreated p53 KO control BMDMs and p53 KO Sen(IR) BMDMs compared with their WT counterparts (Fig. 1J).

To determine whether p21 contributes to the p53-dependent regulation of macrophage survival post IR, we next examined *Cdkn1a*^−/−^ (p21 KO) BMDMs (Fig. 1K). Similar to p53 KO cells, p21 deficiency resulted in increased cleaved caspase-3 expression two days post-IR compared with WT BMDMs (Fig. 1L). Consistent with this increase in apoptotic signaling, Annexin V/PI staining revealed an approximately 1.3-fold increase in total cell death in p21 KO BMDMs compared with WT BMDMs two days post-IR (Fig. 1M,N). Although this increase was less pronounced than the approximately 2.2-fold increase observed following p53 loss, these findings indicate that p21 contributes to p53-dependent regulation of macrophage survival following genotoxic stress.

Despite an approximately 64% reduction in live p21 KO Sen(IR) BMDMs compared with WT Sen(IR) BMDMs 10 days post-IR, the surviving p21 KO Sen(IR) BMDMs retained SA-β-gal activity (Supp. Fig. 1C, Fig. 1O). Consistent with these findings, SPiDER-gal analysis confirmed detectable β-gal activity in p21 KO Sen(IR) BMDMs, with no significant difference in SA-β-gal-associated fluorescence compared with WT Sen(IR) cells (Supp. Fig. 1D). Notably, despite loss of the canonical senescent cell cycle regulator, p21 KO Sen(IR) BMDMs retained the cell cycle arrest characteristic of senescent macrophages, exhibiting an approximately 76% reduction in S-phase entry following IR compared with p21 KO control cells, as measured by Click-iT EdU incorporation (Fig. 1P,Q). Because chronic p21 KO macrophages may develop compensatory pathways, we instead generated acute p21-knockout macrophages using CRISPR-Cas9 ribonucleoprotein (RNP)-mediated approach (Supp. Fig. 1F). We targeted differentiating monocytes with RNP complexes containing guide RNAs targeting either *Cdkn1a* or the *Rosa26* allele, with Rosa26-targeted macrophages serving as WT controls. Following macrophage differentiation, the edited macrophages were subjected to our *in vitro* model of macrophage senescence. Loss of p21 was confirmed by western blot across the senescence time course (Supp. Fig. 1G), and CRISPR-mediated *Cdkn1a* deficiency similarly retained robust cell-cycle arrest following IR, exhibiting a significant reduction in S-phase entry comparable to that observed in the whole-body p21 KO model (Supp. Fig. 1H,I). Loss of p21 did not affect the establishment of the senescence-associated cell cycle arrest in this context. In parallel, loss of p21 resulted in increased *Cdkn2a* (p16) expression at both the transcript and protein levels (Fig. 1R; Supp. Fig. 1E), further supporting negative regulation of p16 by the p53–p21 axis. Collectively, these findings establish the p53–p21 axis as a central regulator of senescent macrophage viability, uncoupling cell-cycle arrest from cell survival and revealing antagonistic suppression of p16 during both acute genotoxic stress and cellular senescence.

### p16 is dispensable for senescence macrophage programming

In contrast to its canonical association with cellular senescence, *Cdkn2a* (p16) expression was consistently downregulated at both the mRNA and protein levels following irradiation (Fig. 2A,C), suggesting that p16 downregulation is a characteristic feature of macrophage senescence in response to genotoxic stress. Given our observation that p16 is negatively regulated by the p53–p21 axis, this finding further supports a model in which p53–p21 signaling, rather than p16 induction, characterizes the macrophage response to genotoxic stress and cellular senescence. We therefore asked whether p16 downregulation is required for the macrophage senescence program or instead reflects the dispensability of p16 in this context.

**Figure 2.**
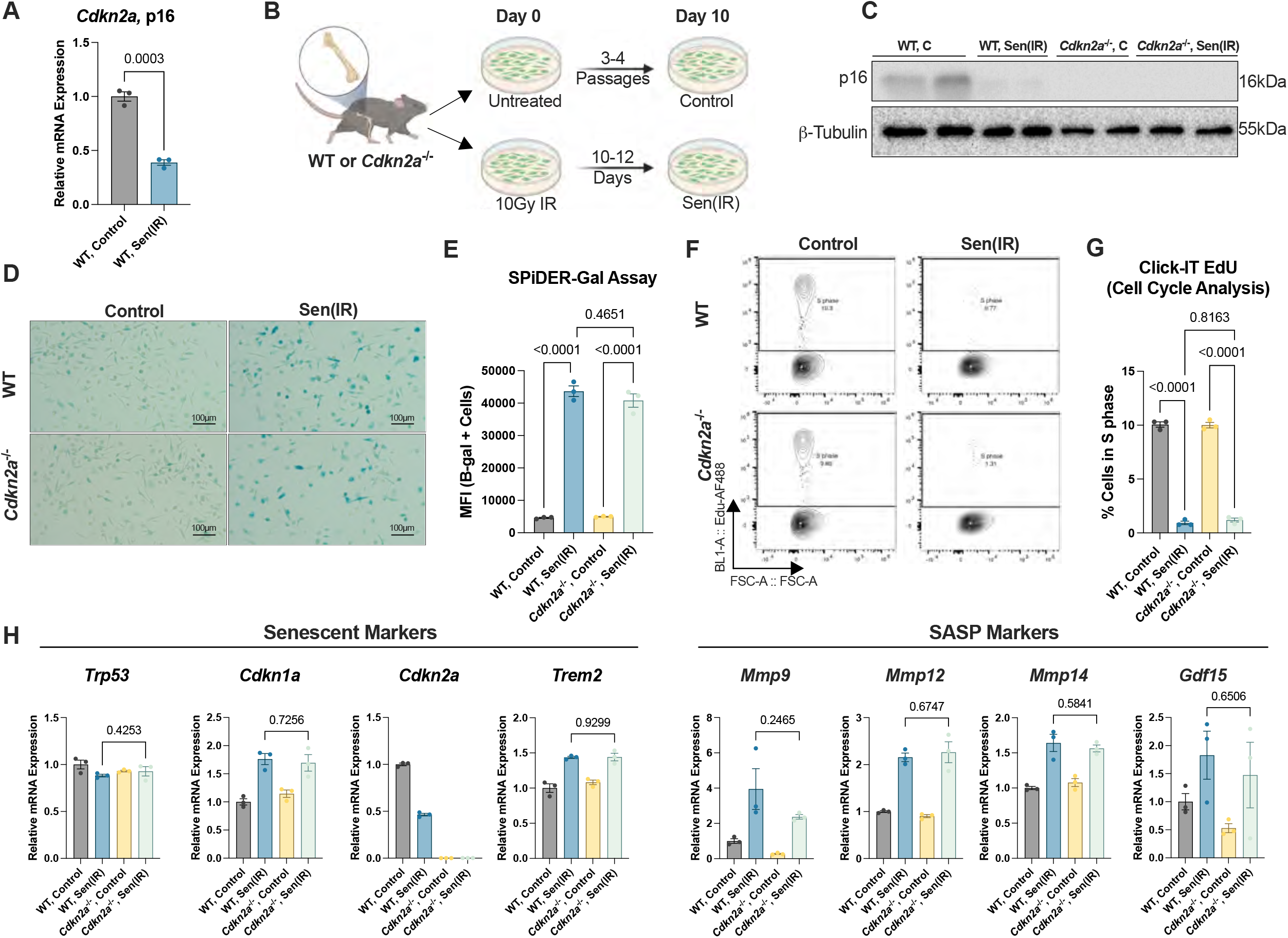
p16 is downregulated in response to genotoxic stress in macrophage senescent programming. (A) Mean ± s.e.m. *Cdkn2a* (p16) mRNA transcript levels, relative to control, in WT control and Sen(IR) macrophages. *P* value of unpaired two-tailed Student’s *t*-test of *n* = 3 biological replicates. (B) Experimental schematic for induction of irradiation-induced senescence in BMDMs from WT or *Cdkn2a*^⁻/⁻^ mice. BMDMs were either left untreated or exposed to 10 Gy IR and cultured for 10 days to generate control and Sen(IR) macrophages, respectively. Created with BioRender.com. (C) SDS-PAGE gels and immunostaining (western blot) for p16 in control and Sen(IR) WT and *Cdkn2a*^⁻/⁻^ macrophages. Β-Tubulin was used as a loading control. (D) Representative images of senescence-associated β-galactosidase (SA-β-gal) staining in control and Sen(IR) WT and *Cdkn2a*^⁻/⁻^ macrophages. Images were acquired at 20X magnification. I Mean ± s.e.m. β-galactosidase activity, measured by SPiDER-β-gal staining, in control and Sen(IR) WT and *Cdkn2a*^⁻/⁻^ macrophages. *P* value of one-way ANOVA with Tukey’s multiple-comparisons test of *n* = 3 biological replicates. (F) Representative flow cytometry plots of EdU incorporation in control and Sen(IR) WT and *Cdkn2a*^⁻/⁻^ macrophages. (G) Mean ± s.e.m. percentage of cells in S phase based on EdU incorporation in control and Sen(IR) WT and *Cdkn2a*^⁻/⁻^ macrophages. *P* value of one-way ANOVA with Tukey’s multiple-comparisons test of *n* = 3 biological replicates. (H) Mean ± s.e.m. mRNA transcript levels, relative to control, for senescence-associated markers (*Trp53*, *Cdkn1a*, *Cdkn2a*, and *Trem2*) and SASP-associated markers (*Mmp9*, *Mmp12*, *Mmp14*, and *Gdf15*) in control and Sen(IR) WT and *Cdkn2a*^⁻/⁻^ macrophages. *P* value of unpaired two-tailed Student’s *t*-test of *n* = 3 biological replicates.

To directly test this, we isolated monocytes from *Cdkn2a*^fl/fl^, LysM-Cre^−/−^ (WT) and *Cdkn2a*^fl/fl^, LysM-Cre^+/−^ (p16 KO) mice and subjected them to our *in vitro* model of macrophage senescence (Fig. 2B). We independently validated the genetic knockout, and the specificity of our p16 antibody, by confirming loss of p16 protein in the KO BMDMs (Fig. 2C). Despite p16 deficiency, p16 KO Sen(IR) macrophages exhibited comparable SA-β-gal activity and morphology to WT Sen(IR) macrophages (Fig. 2D,E). Consistent with these findings, loss of p16 did not impair establishment of the cell-cycle arrest characteristic of senescent macrophages, as p16 KO Sen(IR) macrophages exhibited an approximately 88% reduction in S-phase entry following IR compared with p16 KO control cells, with no significant difference in S-phase entry between p16 KO and WT Sen(IR) macrophages (Fig. 2F,G).

Moreover, p16 deficiency had no effect on the expression of key macrophage senescence-associated genes, including *Trp53*, *Cdkn1a*, and *Trem2*, nor on canonical SASP factors such as *Mmp9*, *Mmp12*, *Mmp14*, and *Gdf15* (Fig. 2H). Together, these findings challenge the canonical role of p16 as a primary regulator of macrophage senescence and demonstrate that macrophage senescence is possible independently of p16.

Given that loss of p53 or p21 resulted in increased early cell death accompanied by restoration of p16 expression, we next asked whether p16 downregulation contributes to the survival phenotype associated with the p53–p21 axis. Consistent with this possibility, p16-deficient BMDMs exhibited a modest but significant increase in the proportion of live cells and a corresponding reduction in total cell death two days post-IR compared with WT BMDMs (Supp. Fig. 2A,B). Notably, expression of *Bcl11a*, a transcriptional regulator that has been shown to promote expression of pro-survival genes including *Bcl2*^11^, was significantly elevated in p16-deficient BMDMs at this early time point, whereas expression of *Bcl2*, a well-established pro-survival gene^12^, was not yet increased (Supp. Fig. 2C,D). These findings suggest that the early survival advantage associated with p16 loss precedes *Bcl2* induction and may involve alternative pro-survival mechanisms. By 12 days post-IR, the survival advantage was no longer observed, while *Bcl11a* expression was no longer elevated and *Bcl2* expression was increased in p16 KO Sen(IR) macrophages (Supp. Fig. 2E-H). Together, these findings suggest that p16 downregulation may contribute to macrophage survival during the acute response to DNA damage via distinct pro-survival programs, with early *Bcl11a* induction preceding later *Bcl2* upregulation. Thus, p16 downregulation may modulate macrophage survival during the early response to genotoxic stress, but is not required for establishment or maintenance of the broader senescent macrophage program.

### Cyclin D2 is a downstream effector of the p53–p21 axis

To identify downstream effectors of the p53–p21 axis that may contribute to the senescent macrophage phenotype, we performed bulk RNA sequencing of WT, p53 KO, and p21 KO Sen(IR) BMDMs alongside their non-irradiated controls. Principal component analysis (PCA) showed that PC1 explained 41.5% of the transcriptional variance and separated non-irradiated controls from irradiated samples across genotypes. PC3 explained an additional 12.2% of the variance and showed genotype-associated separation, with p53- and p21-deficient Sen(IR) BMDMs positioned closer to their respective non-irradiated controls than to WT Sen(IR) cells (Fig. 3A). This separation suggests that loss of p53 or p21 alters the transcriptional response to genotoxic stress and results in a transcriptional state distinct from that of WT Sen(IR) BMDMs. Consistent with this, differential expression analysis revealed widespread transcriptional differences in p53- and p21-deficient Sen(IR) BMDMs compared with WT Sen(IR) cells (Fig. 3B,C).

**Figure 3.**
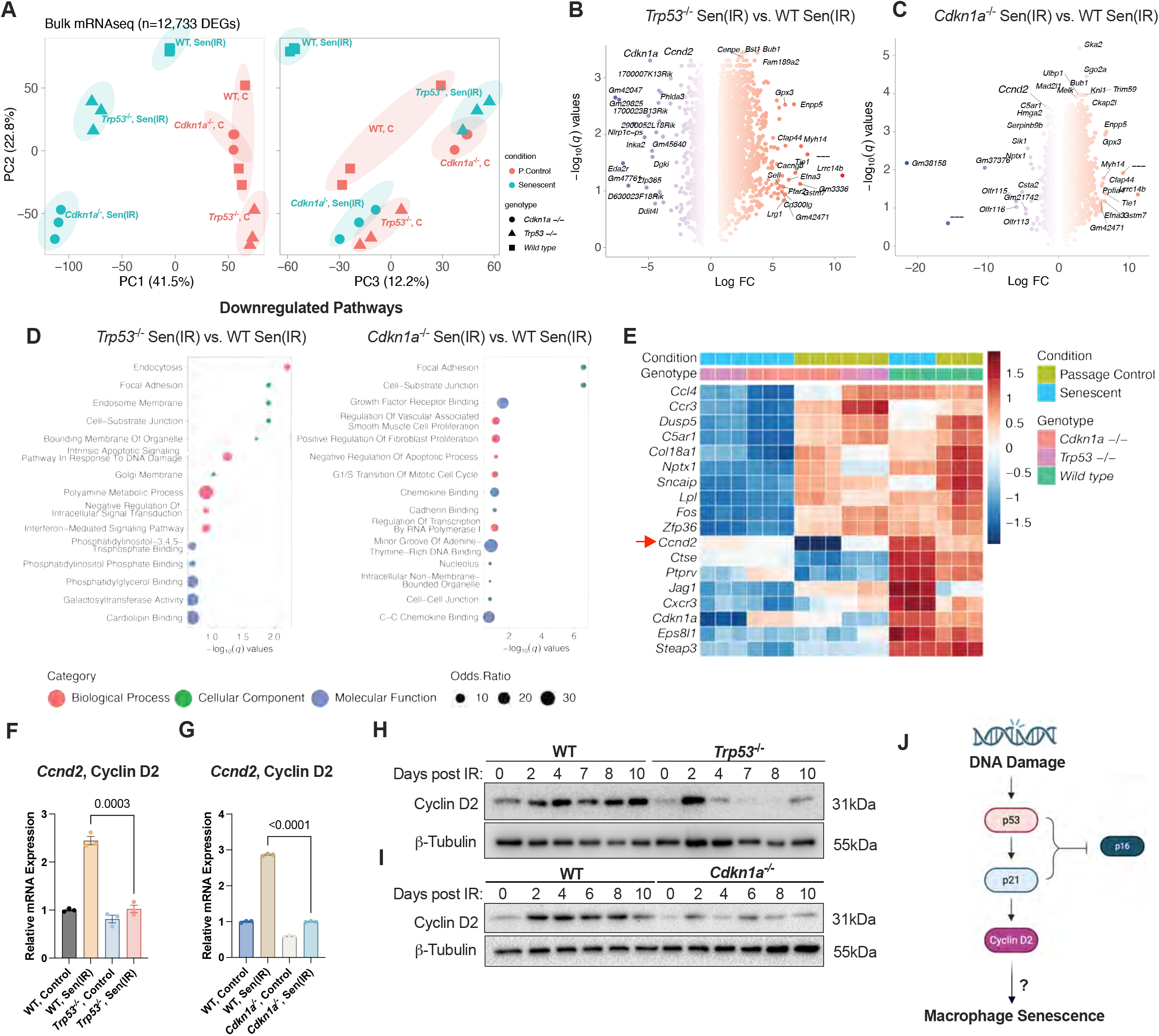
Cyclin D2 is upregulated in senescent macrophages, in a p53–p21-dependent manner. (A) Principal component analysis (PCA) of bulk RNA-seq data from control and Sen(IR) WT, *Trp53*^⁻/⁻^, and *Cdkn1a*^⁻/⁻^ macrophages. PC1, PC2, and PC3 explain 41.5%, 22.8%, and 12.2% of the variance, respectively. (B) Volcano plot showing differentially expressed genes between *Trp53*^⁻/⁻^ and WT Sen(IR) macrophages. (C) Volcano plot showing differentially expressed genes between *Cdkn1a*^⁻/⁻^ and WT Sen(IR) macrophages. (D) Gene ontology analysis of downregulated pathways in *Trp53*^⁻/⁻^ and *Cdkn1a*^⁻/⁻^ Sen(IR) macrophages compared with WT Sen(IR) macrophages. I Heatmap of the top 18 genes commonly downregulated in Sen(IR) macrophages following loss of either *Trp53* or *Cdkn1a*. The red arrow identifies *Ccnd2* (Cyclin D2), the candidate gene of interest. (F) Mean ± s.e.m. *Ccnd2* mRNA transcript levels, relative to control, in WT and *Trp53*^⁻/⁻^ macrophages following senescence induction. *P* value of unpaired two-tailed Student’s *t*-test of *n* = 3 biological replicates. (G) Mean ± s.e.m. *Ccnd2* mRNA transcript levels, relative to control, in WT and *Cdkn1a*^⁻/⁻^ macrophages following senescence induction. *P* value of unpaired two-tailed Student’s *t*-test of *n* = 3 biological replicates. (H) SDS-PAGE gels and immunostaining (western blot) for Cyclin D2 in WT and *Trp53*^⁻/⁻^ macrophages at the indicated days following irradiation. Β-Tubulin was used as a loading control. (I) SDS-PAGE gels and immunostaining (western blot) for Cyclin D2 in WT and *Cdkn1a*^⁻/⁻^ macrophages at the indicated days following irradiation. Β-Tubulin was used as a loading control. (J) Proposed model depicting the p53–p21–Cyclin D2 axis in macrophage senescence. Created with BioRender.com.

Gene ontology analysis further revealed that genes downregulated upon loss of p53 were enriched in biological processes including polyamine metabolism and interferon-mediated signaling (Fig. 3D). These pathways have established roles in cellular growth, proliferation, and survival, as well as inflammatory signaling in macrophages. Notably, our laboratory has previously identified type I interferon signaling as a characteristic component of macrophage senescence and, consistent with findings from other groups, this pathway has been implicated in the establishment and maintenance of the SASP. Together, these findings indicate that loss of p53 or p21 alters transcriptional programs associated with macrophage senescence and prompted us to identify genes whose induction during senescence depends on the p53–p21 axis.

Among the genes altered by loss of p53 or p21, several were associated with inflammatory and chemokine signaling, including *Ccl4*, *Ccr3*, and *Cxcr3*, consistent with the broader inflammatory programs associated with macrophage senescence. Other differentially expressed genes were linked to cellular metabolism, including *Lpl*, a key regulator of lipid metabolism that hydrolyzes triglyceride-rich lipoproteins and promotes fatty acid uptake by tissues^13^ (Fig. 3E). Thus, loss of p53 or p21 altered diverse transcriptional programs associated with inflammation and cellular metabolism. Among these candidates, we focused on *Ccnd2* (Cyclin D2) because its expression was strongly induced in WT Sen(IR) macrophages but markedly attenuated following loss of either p53 or p21 (Fig. 3E). Notably, Cyclin D2 is classically recognized as a G1/S cell-cycle regulator that promotes cell-cycle progression, and its expression is typically associated with proliferating cells^14^. However, Cyclin D2 induction has also been reported in fibroblasts undergoing cellular senescence, suggesting that its expression may extend beyond its canonical role in proliferating cells ^6,15,16^. Despite these observations, the mechanisms underlying Cyclin D2 upregulation and its functional significance in the senescent state remain poorly understood. Its robust induction in senescent macrophages despite their profound cell-cycle arrest therefore suggested that Cyclin D2 may serve a non-canonical function in the senescent state.

Consistent with the RNA-seq findings, *Ccnd2* mRNA was significantly increased in WT Sen(IR) BMDMs compared with WT controls, whereas this induction was significantly attenuated in both p53 KO and p21 KO Sen(IR) BMDMs (Fig. 3F,G). At the protein level, Cyclin D2 was acutely induced in early time points post DNA damage in WT BMDMs and sustained during the transition into senescence (Fig. 3H,I). This sustained expression was markedly reduced in both p53- and p21-deficient BMDMs, further supporting p53–p21-dependent regulation of Cyclin D2 during macrophage senescence. Consistent with these findings, CRISPR-mediated loss of *Cdkn1a* similarly attenuated Cyclin D2 induction following IR, demonstrating that p21-dependent regulation of Cyclin D2 is reproducible across independent genetic approaches (Supp. Fig. 3A). CCND2 has been reported to be interferon response gene (IRG) and type I interferon signaling is upregulated in both the acute DDR and macrophage senescent programming response ^6,17,18^. Therefore, we next asked whether this pathway also contributes to Cyclin D2 induction. In *Ifnar1*^−/−^ BMDMs, Cyclin D2 induction following IR was attenuated compared with WT BMDMs, indicating that type I interferon signaling also contributes to regulation of Cyclin D2 during macrophage senescence (Supp. Fig. 3B). Thus, Cyclin D2 expression, in response to genotoxic stress and cellular senescence is co-regulated by both the p53–p21 and type I interferon pathways.

Finally, the regulatory hierarchy of Cyclin D2 was supported by reciprocal genetic analyses. Loss of *Ccnd2* did not alter *Trp53* or *Cdkn1a* expression in either control or Sen(IR) BMDMs, indicating that Cyclin D2 does not act upstream to regulate the p53–p21 axis (Supp. Fig. 3C,D). Conversely, loss of p16 did not affect *Ccnd2* induction following IR (Supp. Fig. 3E), further demonstrating that Cyclin D2 regulation is specific to the p53–p21 axis. These findings identify Cyclin D2 as a downstream effector of p53–p21 signaling during macrophage senescence (Fig. 3J).

### Cyclin D2 regulates metabolic reprogramming in senescent macrophages

Having established that Cyclin D2 is induced downstream of the p53–p21 axis during macrophage senescence, we next asked what occurs downstream of Cyclin D2 to support the senescent macrophage program. Given its canonical role as a regulator of cell-cycle progression, we first asked whether Cyclin D2 contributes to other established features of macrophage senescence. BMDMs from WT and *Ccnd2*^−/−^ (Cyclin D2 KO) mice were subjected to our *in vitro* model of macrophage senescence (Fig. 4A). Despite robust loss of Cyclin D2, Cyclin D2 KO Sen(IR) macrophages retained SA-β-gal activity and exhibited a comparable morphology to WT Sen(IR) cells (Supp. Fig 4A; Fig. 4B). Similarly, Cyclin D2 KO Sen(IR) macrophages maintained the cell-cycle arrest characteristic of senescent macrophages, with no significant difference in S-phase entry compared with WT Sen(IR) cells (Fig. 4C,D). Thus, despite its canonical role in promoting cell-cycle progression, Cyclin D2 is not required for establishment of the senescence-associated cell-cycle arrest or SA-β-gal-lysosomal phenotype.

**Figure 4.**
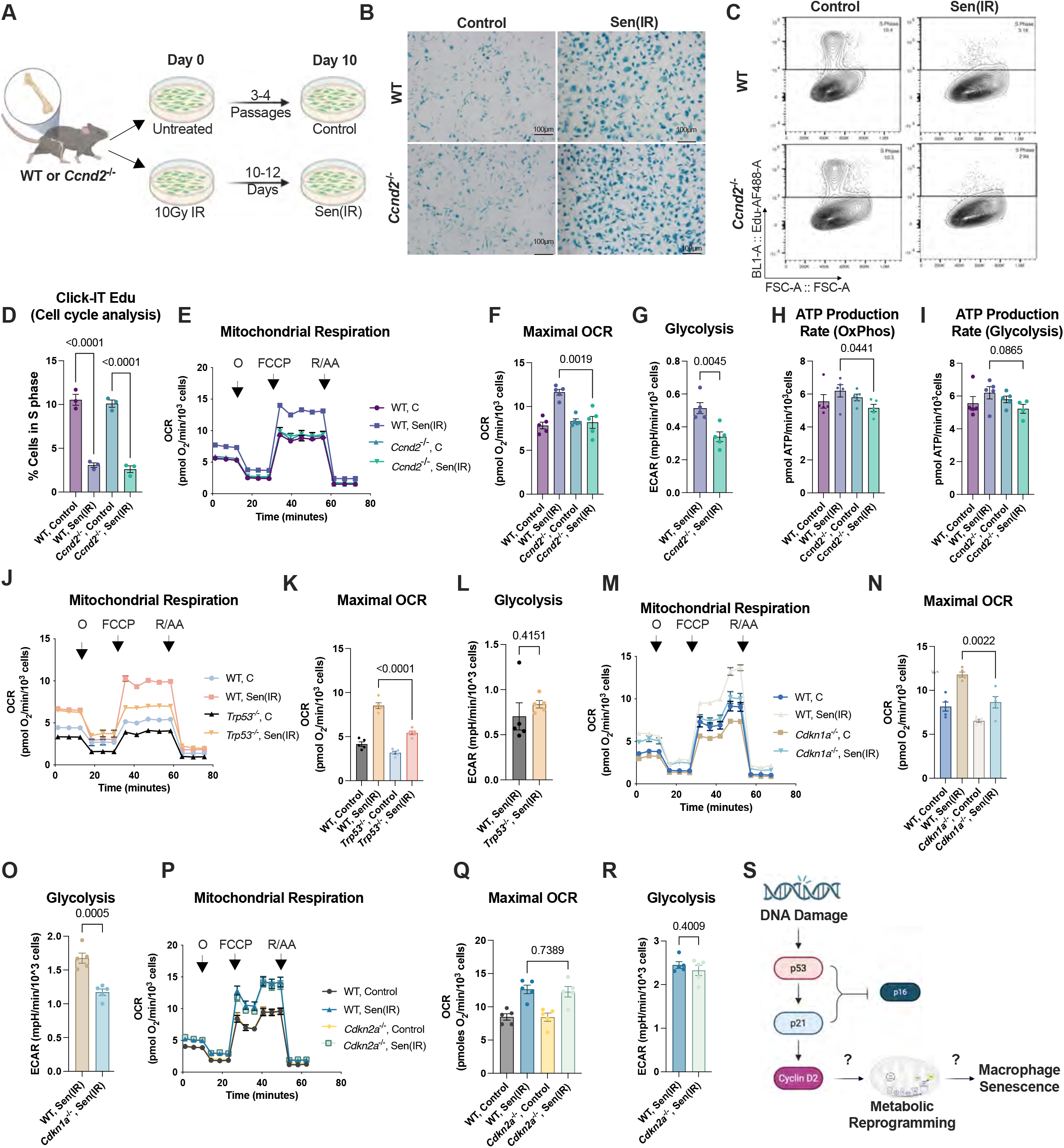
Cyclin D2 mediates p53–p21-dependent metabolic reprogramming during macrophage senescence. (A) Experimental schematic for induction of irradiation-induced senescence in BMDMs from WT or *Ccnd2*^⁻/⁻^ mice. BMDMs were either left untreated or exposed to 10 Gy IR and cultured for 10 days to generate control and Sen(IR) macrophages, respectively. Created with BioRender.com. (B) Representative images of senescence-associated β-galactosidase (SA-β-gal) staining in control and Sen(IR) WT and *Ccnd2*^⁻/⁻^ macrophages. (C) Representative flow cytometry plots of EdU incorporation in control and Sen(IR) WT and *Ccnd2*^⁻/⁻^ macrophages. (D) Mean ± s.e.m. percentage of cells in S phase based on EdU incorporation in control and Sen(IR) WT and *Ccnd2*^⁻/⁻^ macrophages. *P* value of ordinary one-way ANOVA of *n* = 3 biological replicates. I Representative mitochondrial respiration profiles measured by extracellular flux analysis in control and Sen(IR) WT and *Ccnd2*^⁻/⁻^ macrophages. Sequential injections of oligomycin (O), FCCP, and rotenone/antimycin A (R/AA) were used to assess mitochondrial respiratory parameters. (F) Mean ± s.e.m. maximal oxygen consumption rate (OCR) from technical replicates in the representative experiment shown in E. *P* value of unpaired two-tailed Student’s *t*-test. (G) Mean ± s.e.m. glycolytic rate from technical replicates in the representative experiment shown in E. *P* value of unpaired two-tailed Student’s *t*-test. (H) Mean ± s.e.m. ATP production rate through oxidative phosphorylation (OxPhos) from technical replicates in the representative experiment shown in E. *P* value of unpaired two-tailed Student’s *t*-test. (I) Mean ± s.e.m. ATP production rate through glycolysis from technical replicates in the representative experiment shown in E. *P* value of unpaired two-tailed Student’s *t*-test. (J) Representative mitochondrial respiration profiles measured by extracellular flux analysis in control and Sen(IR) WT and *Trp53*^⁻/⁻^ macrophages. Oligomycin, FCCP, and R/AA were sequentially injected as indicated. (K) Mean ± s.e.m. maximal OCR from technical replicates in the representative experiment shown in J. *P* value of unpaired two-tailed Student’s *t*-test. (L) Mean ± s.e.m. glycolytic rate from technical replicates in the representative experiment shown in J. *P* value of unpaired two-tailed Student’s *t*-test. (M) Representative mitochondrial respiration profiles measured by extracellular flux analysis in control and Sen(IR) WT and *Cdkn1a*^⁻/⁻^ macrophages. Oligomycin, FCCP, and R/AA were sequentially injected as indicated. (N) Mean ± s.e.m. maximal OCR from technical replicates in the representative experiment shown in M. *P* value of unpaired two-tailed Student’s *t*-test. (O) Mean ± s.e.m. glycolytic rate from technical replicates in the representative experiment shown in M. *P* value of unpaired two-tailed Student’s *t*-test. (P) Representative mitochondrial respiration profiles measured by extracellular flux analysis in control and Sen(IR) WT and *Cdkn2a*^⁻/⁻^ macrophages. Oligomycin, FCCP, and R/AA were sequentially injected as indicated. (Q) Mean ± s.e.m. maximal OCR from technical replicates in the representative experiment shown in P. *P* value of unpaired two-tailed Student’s *t*-test. I Mean ± s.e.m. glycolytic rate from technical replicates in the representative experiment shown in P. *P* value of unpaired two-tailed Student’s *t*-test. (S) Proposed model depicting the role of Cyclin D2 downstream of the p53–p21 axis in regulating metabolic reprogramming during macrophage senescence. Created with BioRender.com.

Macrophage activation is highly intertwined with metabolic reprogramming which supports stimuli-specific effector functions^19^. Given the sustained SASP phenotype associated with senescent macrophages, we hypothesized macrophage senescence is also associated with metabolic shifts to support this energetically demanding phenotype. In support of this, senescent macrophages exhibited increased mitochondrial respiration and glycolytic capacity relative to non-senescent control cells (Fig. 4E-G). Notably, loss of Cyclin D2 significantly reduced both maximal mitochondrial respiration and glycolytic activity in Sen(IR) macrophages (Fig. 4E-G). Consistent with reduced metabolic capacity, Cyclin D2 KO Sen(IR) macrophages exhibited decreased ATP production rate through both oxidative phosphorylation and glycolysis compared with WT Sen(IR) macrophages (Fig. 4H,I).

To determine whether this metabolic phenotype was dependent on the upstream p53–p21 pathway, we next examined mitochondrial respiration and glycolysis in p53- and p21-deficient Sen(IR) BMDMs. Indeed, loss of p53 markedly attenuated the increase in maximal mitochondrial respiration observed following IR, indicating that p53 contributes to the metabolic reprogramming that accompanies macrophage senescence (Fig. 4J-K). However, loss of p53 did not alter glycolytic capacity, suggesting that the glycolytic component of the senescence-associated metabolic response can be maintained independently of p53. This distinction is consistent with the context-dependent and pathway-specific effects of p53 on cellular metabolism, which can differentially regulate mitochondrial oxidative metabolism and glycolysis^20^. Similarly, p21 deficiency resulted in reduced maximal mitochondrial respiration and glycolytic activity in Sen(IR) macrophages compared with WT Sen(IR) cells (Fig. 4M-O). In contrast, loss of p16 did not significantly alter the senescence-associated increase in mitochondrial respiratory capacity or glycolysis (Fig. 4P-R). Together, these findings place the metabolic reprogramming observed during macrophage senescence under the regulation of the p53–p21 axis, with Cyclin D2 acting as a downstream effector of this metabolic response.

To validate these findings, we employed our CRISPR/Cas9 ribonucleoprotein (RNP)-mediated approach to target *Ccnd2* (Supp. Fig. 4B). Efficient loss of Cyclin D2 was confirmed by western blot analysis (Supp. Fig. 4C). CRISPR-mediated *Ccnd2* deficiency recapitulated the metabolic phenotype observed in the whole-body knockout model, with reduced maximal mitochondrial respiration and glycolytic activity in Sen(IR) macrophages (Supp. Fig. 4D-F). CRISPR-mediated *Ccnd2* deficiency also reduced ATP production rate through glycolysis, whereas the reduction in ATP production rate through oxidative phosphorylation observed in the whole-body knockout did not reach statistical significance in the CRISPR model (Supp. Fig. 4G,H). This difference may reflect distinct consequences of whole-body versus CRISPR-Cas9 genetic approaches. Consistent with the genetic knockout data, CRISPR-mediated loss of *Cdkn1a* similarly reduced mitochondrial respiratory capacity and glycolytic activity in Sen(IR) macrophages (Supp. Fig. 4I-K), further supporting regulation of this metabolic phenotype downstream of the p53–p21–Cyclin D2 axis.

Together, these findings demonstrate that Cyclin D2 serves an unexpected, non-canonical metabolic function in senescent macrophages, regulating the mitochondrial and glycolytic adaptations that support the senescent state independently of cell-cycle arrest (Fig. 4S).

### Cyclin D2 regulates bioenergetic signaling via AKT1-mTORC1 to sustain metabolic reprogramming in senescent macrophages

Having established that Cyclin D2 is required for the metabolic reprogramming that accompanies macrophage senescence, we next sought to define the molecular pathways regulated by Cyclin D2 to support this metabolic state. We therefore integrated transcriptomic, phosphoproteomic, and metabolomic approaches to identify molecular pathways altered by loss of Cyclin D2 in Sen(IR) macrophages. PCA of the transcriptomic dataset revealed that irradiation was the dominant source of transcriptional variation, with PC1 and PC2 accounting for 79.4% and 7.0% of the total variance, respectively (Fig. 5A). Despite the relatively modest contribution of Cyclin D2 loss to overall transcriptional variation, comparison of *Ccnd2*^−/−^ and WT Sen(IR) macrophages identified a discrete set of differentially expressed genes, with 2.95% of detected genes significantly upregulated and 1.92% significantly downregulated following Cyclin D2 loss (Fig. 5B; Supplementary Fig. 5A). Thus, D2 regulates a defined subset of genes in senescent macrophages.

**Figure 5.**
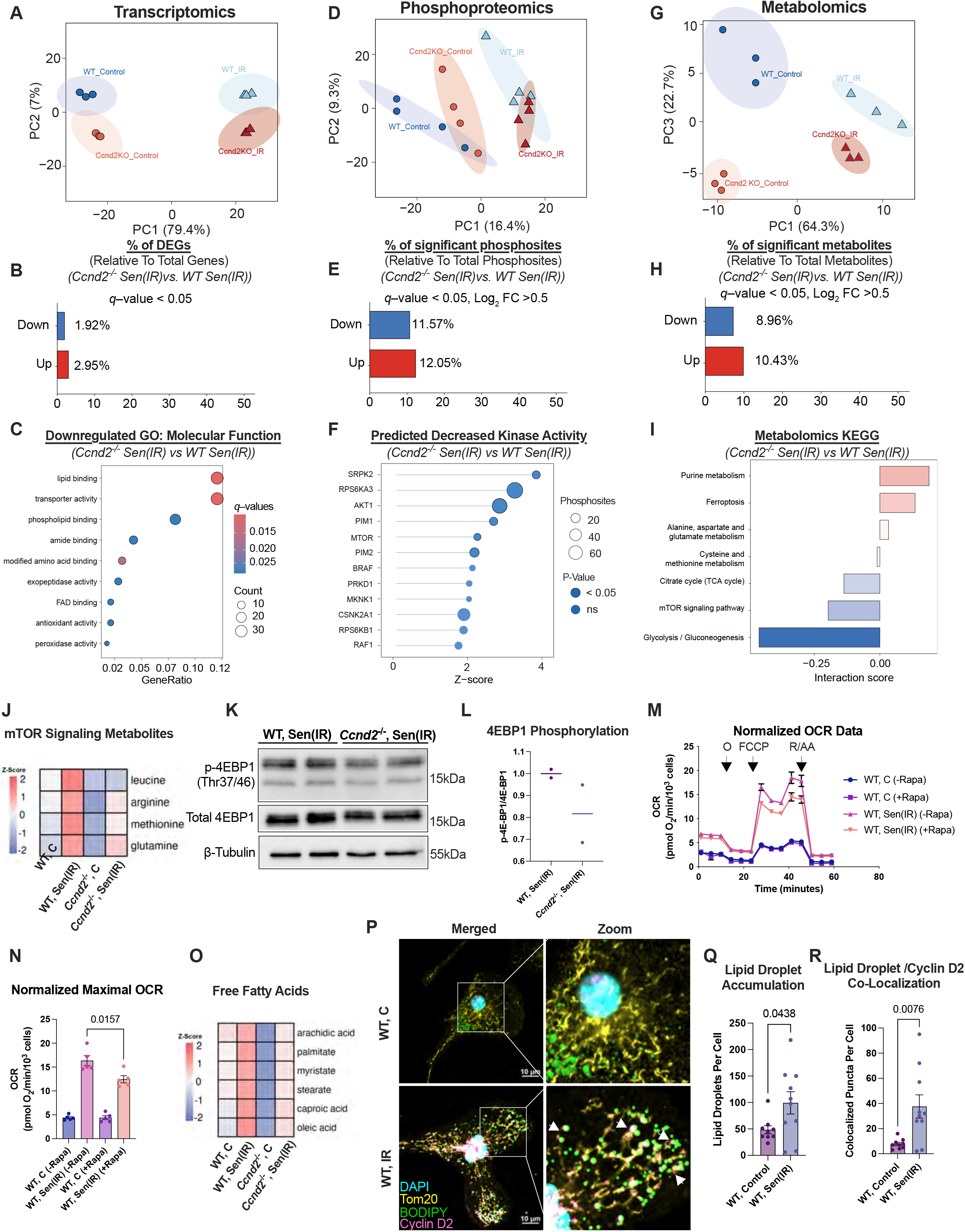
Cyclin D2 regulates bioenergetic signaling via AKT1-mTORC1 signaling to sustain metabolic reprogramming in senescent macrophages. (A) Principal component analysis (PCA) of transcriptomic profiles from control and Sen(IR) WT and *Ccnd2*^⁻/⁻^ macrophages. (B) Percentage of differentially expressed genes (DEGs) identified in *Ccnd2*^⁻/⁻^ Sen(IR) macrophages compared with WT Sen(IR) macrophages. (C) Gene ontology analysis of downregulated molecular functions in *Ccnd2*^⁻/⁻^ Sen(IR) macrophages compared with WT Sen(IR) macrophages. (D) Principal component analysis (PCA) of phosphoproteomic profiles from control and Sen(IR) WT and *Ccnd2*^⁻/⁻^ macrophages. I Percentage of significantly altered phosphosites identified in *Ccnd2*^⁻/⁻^ Sen(IR) macrophages compared with WT Sen(IR) macrophages. (F) Predicted kinase activity analysis highlighting kinases with decreased inferred activity, including AKT1- and mTOR-associated kinases. (G) Principal component analysis (PCA) of metabolomic profiles from control and Sen(IR) WT and *Ccnd2*^⁻/⁻^ macrophages. (H) Percentage of significantly altered metabolites identified in *Ccnd2*^⁻/⁻^ Sen(IR) macrophages compared with WT Sen(IR) macrophages. (I) KEGG pathway analysis of altered metabolites in *Ccnd2*^⁻/⁻^ Sen(IR) macrophages compared with WT Sen(IR) macrophages, identifying glycolysis/gluconeogenesis, mTOR signaling, the citrate cycle, amino acid metabolism, ferroptosis, and purine metabolism among the altered pathways. (J) Heatmap of metabolites associated with mTOR signaling in WT and *Ccnd2*^⁻/⁻^ control and Sen(IR) macrophages. (K) SDS-PAGE gels and immunostaining (western blot) for phosphorylated 4E-BP1 (Thr37/46) and total 4E-BP1 in WT and *Ccnd2*^⁻/⁻^ Sen(IR) macrophages. Β-Tubulin was used as a loading control. (L) Mean phosphorylated 4E-BP1 normalized to total 4E-BP1 expression from technical replicates. (M) Representative mitochondrial respiration profiles measured by extracellular flux analysis in WT control and Sen(IR) macrophages treated with vehicle or 50nM rapamycin for 24h. Oligomycin, FCCP, and R/AA were sequentially injected as indicated. (N) Mean ± s.e.m. maximal oxygen consumption rate (OCR) from technical replicates in the representative experiment shown in M. *P* value of unpaired two-tailed Student’s *t*-test. (O) Heatmap of fatty acids altered in *Ccnd2*^⁻/⁻^ Sen(IR) macrophages compared with WT Sen(IR) macrophages. (P) Representative images of lipid droplets, mitochondria, and Cyclin D2 localization in control and Sen(IR) WT macrophages. Cells were stained with BODIPY, Tom20, and Cyclin D2, with DAPI used to visualize nuclei. (Q) Mean ± s.e.m. lipid droplets per cell in control and Sen(IR) WT macrophages. Each data point represents lipid droplets within an individual cell. *P* value of unpaired two-tailed Student’s *t*-test. I Mean ± s.e.m. lipid droplet/Cyclin D2 colocalized puncta per cell in control and Sen(IR) WT macrophages. Each data point represents Cyclin D2 puncta (pink) co-localized with lipid droplets within an individual cell. *P* value of unpaired two-tailed Student’s *t*-test.

To determine the functional pathways associated with these transcriptional changes, we performed gene ontology analysis of the downregulated genes. This analysis revealed enrichment for molecular functions related to lipid handling, including lipid binding, phospholipid binding, and transporter activity, as well as FAD binding and antioxidant activity (Fig. 5C). These findings suggested that the transcriptional effects of Cyclin D2 loss are particularly associated with pathways involved in lipid metabolism and cellular redox homeostasis, consistent with the metabolic phenotype observed in *Ccnd2*^−/−^ Sen(IR) macrophages.

We next used unsupervised k-means clustering to identify broader transcriptional patterns across the dataset without restricting the analysis to genes meeting differential-expression thresholds. Consistent with the PCA, the major expression patterns were driven primarily by irradiation status, with three major clusters identified across control and Sen(IR) samples (Supplementary Fig. 5B). Notably, gene ontology analysis of the cluster exhibiting reduced expression following Cyclin D2 loss revealed enrichment for processes related to gene expression and protein synthesis, including ribosome biogenesis, mRNA transport, and regulation of translation, as well as molecular functions related to protein kinase activity and kinase binding (Supplementary Fig. 5C). Together, these analyses indicate that while Cyclin D2 loss produces modest changes in overall transcript abundance, it regulates transcriptional programs associated with lipid metabolism as well as protein synthesis and kinase-dependent signaling. The enrichment of kinase-related functions was particularly notable given the established association of Cyclin D2 with cyclin-dependent kinases^14^ and suggested that Cyclin D2 may influence cellular metabolism through changes in kinase-dependent signaling at the level of protein phosphorylation. We therefore examined the phosphoproteome to identify signaling pathways that may underlie the metabolic alterations observed following loss of Cyclin D2.

PCA of the phosphoproteomic dataset revealed distinct phosphoproteomic states between WT and *Ccnd2*^−/−^ macrophages, with PC1 and PC2 accounting for 16.4% and 9.3% of the variance, respectively (Fig. 5D). Loss of Cyclin D2 resulted in widespread changes in phosphorylation, with 12.05% of detected phosphosites significantly increased and 11.57% significantly decreased (Fig. 5E). To infer changes in kinase activity from these phosphorylation patterns, we performed kinase activity prediction based on changes in known kinase substrate phosphosites. This analysis identified multiple kinases predicted to have decreased activity in *Ccnd2*^−/−^ Sen(IR) macrophages, including AKT1, PIM1, PIM2, RPS6KA3, and, notably, mTOR and RPS6KB1 (Fig. 5F). These findings implicated reduced AKT1-mTORC1 signaling as a potential mechanism underlying the metabolic defects observed following Cyclin D2 loss.

To determine whether these signaling alterations were accompanied by changes in the cellular metabolome, we performed metabolomic profiling of WT and *Ccnd2*^−/−^ macrophages. PCA revealed distinct metabolic states between genotypes, with PC1 and PC3 accounting for 64.3% and 22.7% of the variance, respectively (Fig. 5G). Consistent with the transcriptomic and phosphoproteomic analyses, 10.43% of detected metabolites were significantly increased and 8.96% were significantly decreased following Cyclin D2 loss (Fig. 5H). Pathway analysis of differentially abundant metabolites identified glycolysis/gluconeogenesis and mTOR signaling among the pathways most negatively associated with Cyclin D2 deficiency, together with perturbations in the TCA cycle and several amino acid metabolic pathways (Fig. 5I). The identification of glycolysis as a downregulated pathway is consistent with the reduced glycolytic capacity observed following Cyclin D2 loss (Fig. 4), while the convergence on mTOR signaling across the phosphoproteomic and metabolomic datasets further supported a role for mTOR in Cyclin D2-dependent metabolic reprogramming.

Notably, metabolomic analysis revealed decreased abundance of several amino acids associated with mTOR signaling, including leucine, arginine, methionine, and glutamine, in Cyclin D2 KO Sen(IR) macrophages (Fig. 5J). Given the central role of mTORC1 in integrating nutrient availability with cellular growth and metabolism^21^, and macrophage activation^22^, these changes suggested that Cyclin D2 deficiency may impair mTORC1 signaling through both altered kinase activity and reduced availability of mTOR-regulating nutrients. We therefore validated mTORC1 activity by examining phosphorylation of 4E-BP1 at Thr37/46, a canonical mTORC1 target and regulator of cap-dependent translation. Consistent with the multi-omic analyses, Cyclin D2 KO Sen(IR) macrophages exhibited reduced 4E-BP1 phosphorylation relative to WT Sen(IR) macrophages (Fig. 5K,L), confirming reduced mTORC1 signaling following loss of Cyclin D2.

Because mTOR signaling has been implicated in regulation of the SASP^23,24^ and our data indicate that Cyclin D2-dependent metabolic reprogramming increases both mitochondrial respiration and glycolytic capacity during senescence, we next asked whether mTOR activity contributes to this bioenergetic state. WT control and Sen(IR) macrophages were treated with 50 nM rapamycin for 24 h to inhibit mTORC1. Rapamycin treatment reduced mitochondrial respiratory capacity in Sen(IR) macrophages (Fig. 5M,N) and similarly reduced glycolytic capacity (Supp. Fig. 5D), supporting a functional relationship between mTORC1 signaling and the enhanced bioenergetic capacity characteristic of senescent macrophages.

Given the enrichment of lipid-associated molecular functions among genes downregulated following Cyclin D2 loss, together with the established accumulation of lipids in senescent macrophages, we next examined changes in individual free fatty acid species. Metabolomic profiling revealed decreased abundance of multiple free fatty acids in *Ccnd2*^−/−^ Sen(IR) macrophages, including arachidic acid, palmitate, myristate, stearate, capric acid, and oleic acid (Fig. 5O). These findings suggested that Cyclin D2 may contribute to the altered lipid handling that accompanies macrophage senescence^6^. Cyclin D2 has classically been characterized as a nuclear cell-cycle regulator, and previous studies have reported nuclear localization of Cyclin D2 during growth arrest, including cellular senescence^6,15,16^. However, Cyclin D2 can also undergo nuclear-cytoplasmic redistribution, raising the possibility that changes in its subcellular localization could enable non-canonical functions outside the nucleus^25^. Given our prior report of the accumulation of lipids in senescent macrophages^6^ and our observation that Cyclin D2 loss alters lipid-associated metabolic pathways, we therefore examined whether senescence was accompanied by redistribution of Cyclin D2 toward lipid storage and metabolic compartments. We assessed the spatial relationship between Cyclin D2, mitochondria, and neutral lipids by immunofluorescence staining for Cyclin D2, the mitochondrial marker TOM20, together with the neutral lipid dye BODIPY. In WT non-sen(IR) macrophages, Cyclin D2 was predominantly nuclear, whereas following induction of senescence, Cyclin D2 redistributed to the cytoplasm, where it exhibited increased colocalization with BODIPY-positive lipid droplets and TOM20-positive mitochondria (Fig. 5P, Supp. Fig. 5E). Importantly, Cyclin D2 immunofluorescence signal was absent in Cyclin D2 KO control macrophages, validating the specificity of the antibody and supporting that the cytoplasmic signal observed in WT Sen(IR) macrophages represents bona fide Cyclin D2 rather than nonspecific staining (Supp. Fig. 5E). Senescent macrophages also exhibited a marked increase in lipid droplet accumulation, with lipid droplets frequently observed in close association with mitochondria. Quantification confirmed increased lipid droplet accumulation in Sen(IR) macrophages (Fig. 5Q) as well as increased colocalization between Cyclin D2 and lipid droplets (Fig. 5R).

In sum, these findings identify Cyclin D2 as a regulator of metabolic remodeling during macrophage senescence, linking lipid-associated metabolism and mitochondrial bioenergetics with AKT1-mTORC1 signaling to support the metabolic demands of the senescent state. The convergence of transcriptomic, phosphoproteomic, and metabolomic alterations, together with functional validation of mTORC1 activity and the senescence-associated redistribution of Cyclin D2 toward lipid- and mitochondria-associated compartments, supports a model in which Cyclin D2 coordinates metabolic reprogramming through interconnected metabolic, signaling, and spatial mechanisms.

### Cyclin D2 coordinates lipid and mitochondrial remodeling during macrophage senescence

Given the redistribution of Cyclin D2 toward lipid droplets and mitochondria during senescence, we next asked whether Cyclin D2 contributes to lipid droplet accumulation and lipid-mitochondrial organization. BODIPY staining revealed a robust increase in lipid droplet accumulation following induction of senescence in WT macrophages, whereas the senescence-associated increase was attenuated in *Ccnd2*^−/−^ macrophages (Fig. 6A). Quantification of lipid droplet accumulation confirmed this decrease in total number of lipid droplets per cell in response to senescence following Cyclin D2 loss (Fig. 6B), while quantification of lipid droplets associated with mitochondria revealed a decrease in the number of lipid droplets touching mitochondria in *Ccnd2*^−/−^ Sen(IR) macrophages (Fig. 6C). These findings indicate that Cyclin D2 contributes to the remodeling of lipid storage and association of lipid droplets with mitochondria during macrophage senescence.

**Figure 6.**
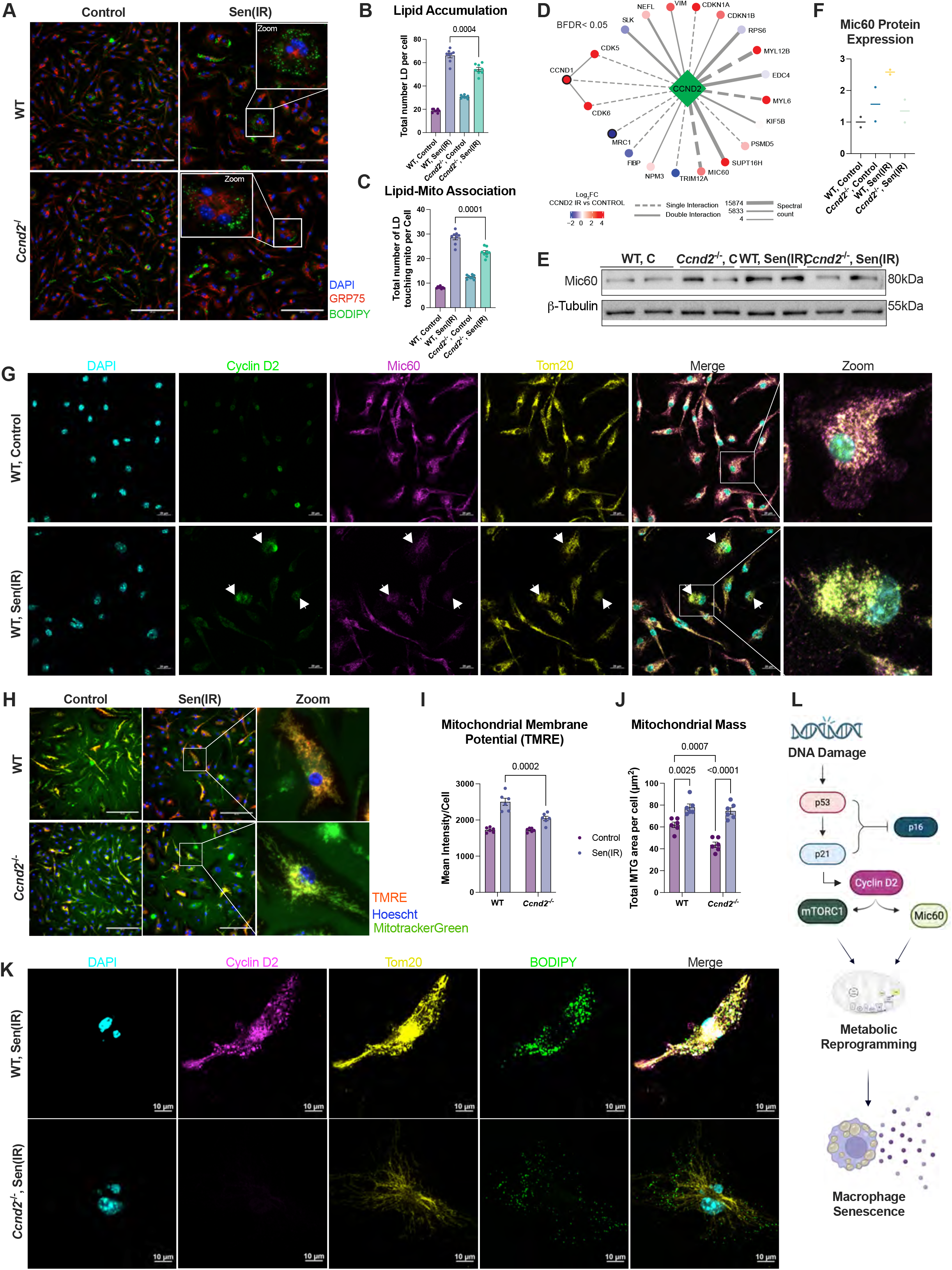
Cyclin D2 regulates mitochondrial organization and function during macrophage senescence. (A) Representative images of control and Sen(IR) WT and *Ccnd2*^⁻/⁻^ macrophages stained with BODIPY to visualize neutral lipids, GRP75-647 to visualize mitochondria, and DAPI to visualize nuclei. (B) Mean ± s.e.m. lipid accumulation in control and Sen(IR) WT and *Ccnd2*^⁻/⁻^ macrophages. *P* value of unpaired two-tailed Student’s *t*-test. (C) Mean ± s.e.m. lipid-associated mitochondrial signal in control and Sen(IR) WT and *Ccnd2*^⁻/⁻^ macrophages. *P* value of unpaired two-tailed Student’s *t*-test. (D) Protein interaction network identified by Cyclin D2 immunoprecipitation followed by mass spectrometry, highlighting candidate Cyclin D2-interacting proteins, including Mic60 and cytoskeletal/structural proteins VIM, MYL6, and MYL12B. I SDS-PAGE gels and immunostaining (western blot) for Mic60 in WT and *Ccnd2*^⁻/⁻^ control and Sen(IR) macrophages. Β-Tubulin was used as a loading control. (F) Mean Mic60 protein expression. (G) Representative immunofluorescence images showing Cyclin D2, Mic60, and the mitochondrial marker Tom20 in WT control and Sen(IR) macrophages. DAPI was used to visualize nuclei. (H) Representative immunofluorescence images showing Cyclin D2, Mic60, Tom20, and BODIPY in WT and *Ccnd2*^⁻/⁻^ Sen(IR) macrophages. (I) Representative images of mitochondrial membrane potential measured by TMRE staining in WT and *Ccnd2*^⁻/⁻^ control and Sen(IR) macrophages. MitoTracker Green was used to visualize total mitochondria, and Hoechst was used to visualize nuclei. (J) Mean ± s.e.m. mitochondrial membrane potential measured by TMRE fluorescence in WT and *Ccnd2*^⁻/⁻^ control and Sen(IR) macrophages. *P* value of two-way ANOVA with Tukey’s multiple-comparisons test. Comparisons were restricted to the Sen(IR) condition between genotypes. (K) Mean ± s.e.m. total mitochondrial content measured by MitoTracker Green fluorescence in WT and *Ccnd2*^⁻/⁻^ control and Sen(IR) macrophages. *P* value of two-way ANOVA with Tukey’s multiple-comparisons test. Comparisons were made between control and Sen(IR) conditions within each genotype and between genotypes under the Sen(IR) condition. (L) Proposed model depicting Cyclin D2-dependent regulation of mitochondrial organization and function downstream of the p53–p21 axis, with Mic60 as a candidate mediator linking Cyclin D2 to mitochondrial organization and metabolic reprogramming during macrophage senescence. Created with <u>BioRender.com</u>.

To identify candidate effectors that could link Cyclin D2 to mitochondrial remodeling, we next characterized the Cyclin D2 interactome by immunoprecipitation followed by mass spectrometry, using Ccnd2^-/-^ macrophages as a control to identify proteins specifically associated with Cyclin D2. This analysis identified Cyclin D2-associated proteins spanning several functional categories, including mitochondrial organization, senescence regulation, cytoskeletal organization, and kinase/signaling pathways (Fig. 6D; Supp. Fig. 6A). Among these candidates, MIC60, also known as Mitofilin/IMMT, is a core component of the mitochondrial contact site and cristae organizing system (MICOS), with established roles in mitochondrial inner-membrane organization and cristae architecture^26,27^, was of particular interest given the mitochondrial metabolic defects observed following Cyclin D2 loss. The interactome also identified other candidates of interest including CDKN1A (p21), consistent with the established p53–p21–Cyclin D2 pathway, as well as cytoskeletal proteins including vimentin (VIM), myosin light chain 6 (MYL6), and myosin light chain 12B (MYL12B) (Fig. 6D; Supp. Fig. 6A). As an orthogonal assessment of the relationship between p21 and Cyclin D2, immunofluorescence revealed increased nuclear co-localization of p21 and Cyclin D2 in WT Sen(IR) macrophages (Supplementary Fig. 6B), consistent with their association within the p53–p21–Cyclin D2 pathway.

We next examined whether candidate cyclin and cyclin-dependent kinase interactors could account for the cell-cycle and metabolic phenotypes associated with Cyclin D2. *Ccnd1* was of particular interest because Cyclin D1 was identified in the Cyclin D2 interactome and was also induced during macrophage senescence (Supplementary Fig. 6A,C), consistent with reports of increased Cyclin D1 expression in other senescent cell types, including hepatocytes^16^. Given the canonical role of Cyclin D1 in promoting cell-cycle progression through CDK4/6^28,29^, we asked whether Cyclin D1 might contribute to cell-cycle arrest during senescence or instead function independently of Cyclin D2’s non-canonical role in metabolic regulation. CRISPR-mediated disruption of *Ccnd1* resulted in loss of Cyclin D1 protein expression (Supplementary Fig. 6D). Notably, *Ccnd1*^−/−^ macrophages exhibited reduced S-phase entry under control conditions, demonstrating a proliferative defect in the absence of Cyclin D1. However, *Ccnd1*^−/−^ macrophages retained the ability to undergo cell-cycle arrest following irradiation (Supplementary Fig. 6E,F). Thus, whereas Cyclin D1 contributes to proliferative cell-cycle progression in control macrophages, its loss does not prevent irradiation-induced cell-cycle arrest.

We then asked whether Cyclin D1-associated CDK4/6 activity contributes to the metabolic reprogramming associated with senescence. Because CDK6 was also identified as a Cyclin D2-interacting protein (Fig. 6D; Supplementary Fig. 6A), we treated WT macrophages with palbociclib, a CDK4/6 inhibitor^30^, and assessed mitochondrial and glycolytic function. Palbociclib treatment did not alter mitochondrial respiration or maximal respiratory capacity in Sen(IR) macrophages (Supplementary Fig. 6G,H), nor did it affect ATP production rate through oxidative phosphorylation (Supplementary Fig. 6I). Similarly, CDK4/6 inhibition did not significantly alter glycolytic capacity or ATP production rate through glycolysis (Supplementary Fig. 6J,K). These findings indicate that canonical Cyclin D1-CDK4/6 activity is dispensable for the enhanced bioenergetic state of senescent macrophages and is therefore unlikely to account for the non-canonical metabolic function of Cyclin D2.

We next focused on MIC60 as a candidate link between Cyclin D2 and mitochondrial function. Western blot analysis revealed that MIC60 protein abundance increased during senescence in WT macrophages, whereas this induction was diminished following loss of Cyclin D2 (Fig. 6E-F), suggesting that Cyclin D2 contributes to the regulation of MIC60 during macrophage senescence. We next examined the spatial relationship between Cyclin D2 and MIC60. In control macrophages, Cyclin D2 was predominantly nuclear, whereas following induction of senescence, Cyclin D2 redistributed to the cytoplasm and exhibited increased spatial overlap with MIC60 and TOM20-positive mitochondria (Fig. 6G). These findings were consistent with the identification of MIC60 as a Cyclin D2-associated protein and further supported a senescence-associated redistribution of Cyclin D2 toward mitochondria.

Given the established role of MIC60 in maintaining mitochondrial architecture and function^31^, we next examined whether the mitochondrial phenotypes associated with Cyclin D2 loss were accompanied by changes in mitochondrial function. Indeed, loss of Cyclin D2 was associated with reduced mitochondrial membrane potential, as assessed by TMRE staining (Fig. 6H-J). These findings were consistent with the reduced mitochondrial respiration and ATP production rates observed following Cyclin D2 loss (Fig. 4) and with reported consequences of impaired MIC60 function^31^.

To determine if Cyclin D2 loss was associated with changes in mitochondrial abundance and organization, we stained mitochondria with MitoTracker Green and quantified average mitochondrial area and length. Induction of senescence increased mitochondrial mass in both WT and *Ccnd2*^−/−^ BMDMs, accompanied by changes in mitochondrial morphology indicative of increased fragmentation of the mitochondrial network (Fig. 6J; Supplementary Fig. 6L,M). This pattern of mitochondrial remodeling has been previously reported in response to cellular stress and senescence^32–34^. Notably, *Ccnd2*^−/−^ control BMDMs exhibited reduced mitochondrial mass relative to WT control cells, indicating that Cyclin D2 loss alters mitochondrial abundance even in the absence of senescence. Consistent with this phenotype, mitochondria in *Ccnd2*^−/−^ control macrophages were smaller and shorter than those in WT control macrophages (Supplementary Fig. 6C,D). Together, these findings suggest that Cyclin D2 deficiency is associated with pre-existing alterations in mitochondrial abundance and architecture that persist during senescence and may contribute to the impaired mitochondrial function observed following Cyclin D2 loss.

Consistent with these quantitative changes, imaging revealed extensive reorganization of the mitochondrial network in WT Sen(IR) macrophages, with mitochondria exhibiting a pronounced perinuclear organization that extended around and appeared to partially envelop the nucleus. In contrast, *Ccnd2*^−/−^ Sen(IR) macrophages displayed markedly reduced mitochondrial-perinuclear contacts (Fig. 6H). Because the Cyclin D2 interactome also identified multiple proteins involved in cytoskeletal organization, including VIM, MYL6, and MYL12B, we considered whether Cyclin D2-associated interactions with the cytoskeletal machinery could contribute to the altered positioning and organization of mitochondria observed following Cyclin D2 loss (Fig. 6K; Supp. Fig. 6A). Although these observations do not establish a causal role for these proteins, they raise the possibility that Cyclin D2-associated cytoskeletal interactions contribute to mitochondrial network organization during senescence.

Collectively, these data support a model in which DNA damage-induced p53–p21 signaling promotes Cyclin D2 expression and subsequent redistribution from the nucleus toward mitochondria and lipid-associated compartments. Through association with MIC60 and potentially other cytoskeletal regulators, Cyclin D2 may coordinate mitochondrial organization and lipid-associated remodeling during senescence. These changes occur in the context of altered mTORC1 signaling and metabolic reprogramming and may contribute to the bioenergetic and biosynthetic state required to sustain the senescent macrophage phenotype and its associated secretory program (Fig. 6L).

### Cyclin D2 and p21 co-expression increases in macrophages during aging and metabolic liver disease in mice

We next sought to determine whether the p53–p21–Cyclin D2 pathway identified in senescent macrophages *in vitro* is associated with macrophage senescence *in vivo*. Because p21^+^ senescent macrophages accumulate in aged tissues, including Kupffer cells^6^, the resident macrophages of the liver, we first examined *Ccnd2* expression in young and aged liver. We observed a significant increase in *Ccnd2* mRNA in old livers (21-25 months) relative to young mice (3-4 months) (Fig. 7A). Consistent with these findings, immunofluorescence analysis demonstrated increased Cyclin D2 and p21 expression and co-localization within CD68^+^ macrophages in aged liver (25 months), compared to liver from a young mouse (4 months), but not in other cell types. This suggests increased p21-Cyclin D2 expression in the aged liver is a macrophage specific phenotype (Fig. 7B-C).

**Figure 7.**
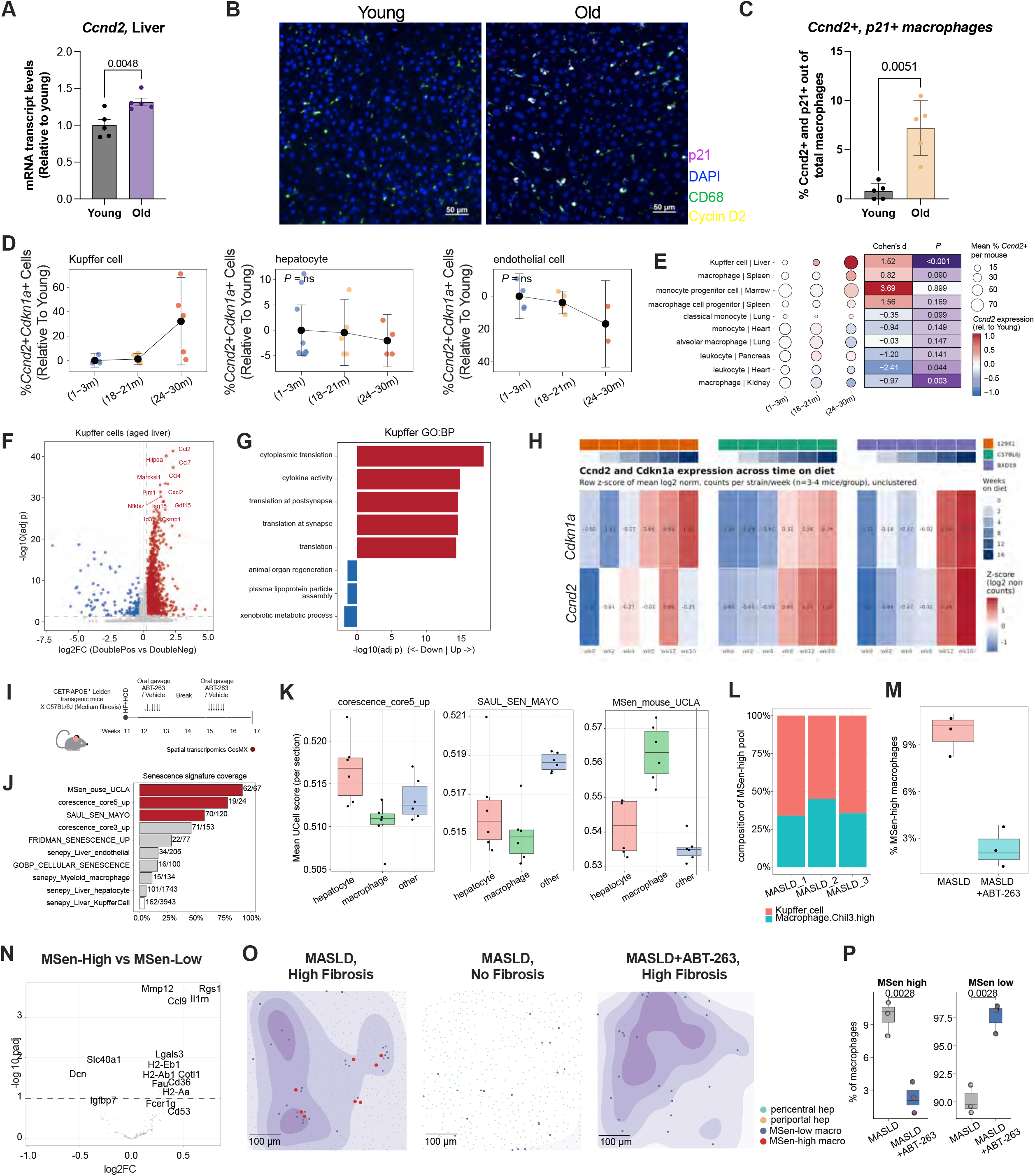
Aging promotes Cyclin D2⁺p21⁺ macrophage accumulation, and spatial transcriptomics identifies a corresponding senescent macrophage population in MASLD targeted by senolytic treatment. (A) Mean ± s.e.m. *Ccnd2* mRNA transcript levels in liver from young and old mice, expressed relative to young mice. *P* value of unpaired two-tailed Student’s *t*-test of *n* = 5 mice per age group. (B) Representative immunofluorescence images of young and old mouse liver stained for p21, CD68, and Cyclin D2, with DAPI used to visualize nuclei. Scale bars, 50 μm. (C) Mean ± s.e.m. Cyclin D2⁺p21⁺ macrophages, expressed as a percentage of total macrophages, in young (4 months) and old (25 months) mouse liver. *P* value of unpaired two-tailed Student’s *t*-test. (D) Quantification of *Ccnd2*⁺*Cdkn1a*⁺ cells in Kupffer cells, hepatocytes, and endothelial cells across young (1-3 months), middle-aged (18-21 months), and old (24-30 months) mice, expressed relative to young mice. (E) Meta-analysis of *Ccnd2* expression across macrophage and immune cell populations from multiple organs in young, middle-aged, and aged mice, showing age-associated changes in *Ccnd2* expression across tissues. Effect sizes are reported as Cohen’s *d*. (F) Volcano plot showing differentially expressed genes between Cyclin D2⁺p21⁺ (double-positive) and Cyclin D2⁻p21⁻ (double-negative) Kupffer cells from aged liver. (G) Gene ontology analysis of differentially expressed genes in Cyclin D2⁺p21⁺ versus Cyclin D2⁻p21⁻ Kupffer cells from aged liver. (H) *Ccnd2* and *Cdkn1a* expression during progression of diet-induced liver disease in mouse strains with differing genetic predispositions to hepatic fibrosis following exposure to a high-fat, high-cholesterol (HFHC) diet. (I) Experimental schematic for spatial transcriptomic analysis of liver tissue from mice fed an HFHC diet with or without senolytic treatment with ABT-263. Created with BioRender.com. (J) Coverage of established senescence gene signatures across cell populations identified by spatial transcriptomics. (K) UCell scores for established senescence signatures across hepatocyte, macrophage, and other cell populations in spatial transcriptomic liver sections. (L) Cellular composition of the MSen-high macrophage population identified by spatial transcriptomics. (M) Quantification of MSen-high macrophages in control and senolytic-treated mice. *n* = 3 mice per treatment group. (N) Volcano plot showing differentially expressed genes between MSen-high and MSen-low macrophages identified by spatial transcriptomics. (O) Spatial distribution of MSen-high and MSen-low macrophages in liver sections from mice fed an HFHC diet, with or without senolytic treatment. Pericentral and periportal hepatocytes are indicated. Scale bars, 100 μm. (P) Quantification of MSen-high and MSen-low macrophages in control and senolytic-treated mice. *n* = 3 mice per treatment group.

To determine and further validate the cellular specificity of this age-associated Cyclin D2/p21 co-expression, we analyzed liver single-cell data from the Tabula Muris Senis Consortium^35^. Age-dependent accumulation of *Ccnd2*^+^*Cdkn1a*^+^ cells was most pronounced in Kupffer cells compared with hepatocytes and endothelial cells, consistent with preferential induction of Cyclin D2 in the resident macrophage population of the aging liver (Fig. 7D). Furthermore, analysis of aged Kupffer cells demonstrated that *Ccnd2*-high expressing cells exhibited increased macrophage senescence scores compared with *Ccnd2*-low expressing cells (Supp. Fig. 7A), supporting an association between Cyclin D2 expression and the senescent macrophage state in aged liver.

We next examined whether age-associated *Ccnd2* expression was restricted to the liver or observed across macrophage populations in other tissues. Analysis across tissues and cell populations in the Tabula Muris Senis dataset revealed tissue- and cell-type-specific changes in *Ccnd2* expression with age, with increased expression observed in several macrophage and immune cell populations, including macrophages in the spleen and progenitor populations in the marrow (Fig. 7E). In aged spleen, *Ccnd2*-expressing macrophages exhibited an inflammatory transcriptional profile characterized by enrichment of TNF production, regulation of TNF production, and inflammatory response programs (Supp. Fig. 7A-C). Together, these findings suggest that the association between Cyclin D2 expression and inflammatory, senescence-associated macrophage states extends beyond the liver in aging mice.

We next examined the transcriptional state of Cyclin D2^+^p21^+^ Kupffer cells in aged liver. Relative to Cyclin D2^−^p21^−^ cells, double-positive Kupffer cells exhibited increased expression of inflammatory genes, including *Ccl2, Ccl7, Ccl4,* and *Cxcl2*, as well as genes associated with stress and inflammatory signaling (Fig. 7F). Gene ontology analysis further revealed enrichment of cytokine activity and translational programs among Cyclin D2^+^p21^+^ Kupffer cells, while genes associated with animal organ regeneration were reduced (Fig. 7G). Together with their p21 expression, these transcriptional features are consistent with a senescence-associated state characterized by inflammatory and secretory activity and reduced tissue-reparative potential.

Given our previous identification of increased p21 and Cyclin D2 expression in senescent macrophages in mouse models of MASLD^6^, we next asked whether induction of the p21-Cyclin D2 axis tracked with disease progression across genetic backgrounds with distinct susceptibility to fibrosis. Across three mouse strains with low, intermediate, or high susceptibility to diet-induced MASLD induced fibrosis, both *Ccnd2* and *Cdkn1a* increased with duration of high-fat, high-cholesterol (HFHC) diet (Fig. 7H). Thus, induction of the p21-Cyclin D2 axis accompanied progression of diet-induced liver disease across distinct genetic backgrounds, indicating that this response was not restricted to a single fibrosis-susceptible strain.

Building on our previous study demonstrating the efficacy of senolytic treatments targeting senescent macrophages in a mouse model of MASLD^6^, we leveraged liver tissue collected from this prior study to perform spatial transcriptomic analysis. Specifically, CETP-APOE Leiden transgenic mice on a C57BL/6J background were subjected to a high-fat, high-cholesterol (HFHC) diet and treated with the senolytic ABT-263 or vehicle using an intermittent “hit-and-run” dosing regimen consisting of 7 days of daily oral gavage followed by a 2-week treatment break and a second 7-day dosing period (Fig. 7I). Livers collected at the end of the treatment paradigm were subsequently analyzed by CosMX spatial transcriptomics to define the cellular and spatial features of senescent macrophages in MASLD (Fig. 7I; Supp. Fig. 7D-I). The resulting spatial transcriptomic datasets met established quality-control criteria based on transcript and gene detection and enabled identification of major hepatic and immune cell populations in both vehicle- and senolytic-treated samples (Supp. Fig. 7D-F). Consistent with the senolytic response observed in our previous study, ABT-263 treatment reduced the proportion of MSen-positive cells in the spatial transcriptomic dataset (Supp. Fig. 7G). We further examined the cellular distribution of *Ccnd2* and *Cdkn1a* within the spatial transcriptomic dataset. Both transcripts were reduced in multiple liver cell types following ABT-263 treatment (Supp. Fig. 7H-I), indicating that the senolytic intervention altered Cyclin D2/p21-associated cell populations beyond macrophages.

We next compared the representation of established senescence-associated gene signatures within the spatial transcriptomic dataset. Our Msen(mouse) signature exhibited robust coverage, with 62 of 67 signature genes detected, along with 19 of 24 genes for the CoreScence signature and 70 of 120 genes for the SenMayo signature (Fig. 7J). We therefore evaluated the ability of these signatures to identify senescence-associated macrophages in vehicle-treated liver. The Msen(mouse) signature showed preferential enrichment in macrophages relative to hepatocytes and other cell populations, supporting its use to identify the Msen macrophage population in the spatial dataset (Fig. 7K).

We next examined the cellular composition of the Msen-high population. The Msen-high pool was predominantly composed of Kupffer cells, with a smaller contribution from a *Chil3*-high macrophage population, indicating that the Msen signature was enriched in resident Kupffer cells rather than being broadly distributed across macrophage populations (Fig. 7L). These findings indicate that the Msen signature preferentially identifies a resident Kupffer cell population rather than broadly labeling macrophages within the diseased liver.

Finally, we assessed whether this Msen-high macrophage population was altered following senolytic treatment. ABT-263 treatment significantly reduced the proportion of Msen-high macrophages, with a corresponding increase in the Msen-low macrophage population (Fig. 7M). Spatial mapping further revealed that Msen-high macrophages were enriched within regions of high fibrosis in vehicle-treated livers but were largely absent from non-fibrotic regions, consistent with a spatial association between Msen-high macrophages and fibrotic pathology (Fig. 7O). Following senolytic treatment, Msen-high macrophages were no longer detected within regions of high fibrosis, which were instead predominantly occupied by Msen-low macrophages (Fig. 7O-P). These findings suggest that Msen-high macrophages are preferentially associated with fibrotic regions and that their depletion following senolytic treatment is accompanied by an increase in the relative representation of Msen-low macrophages within these regions, in improved liver health^6^. Taken together, these findings identify a Cyclin D2-associated senescent macrophage population that accumulates in aging liver, is enriched among resident Kupffer cells, exhibits a distinct transcriptional state in MASLD, and is preferentially localized to fibrotic regions where it can be reduced by intermittent senolytic treatment to reduce MASLD disease and inflammaging^6^.

### Cyclin D2 and p21 are enriched in disease-associated macrophage states in human liver cirrhosis

Having established that Cyclin D2^+^p21^+^ macrophages accumulate during aging and that a corresponding Msen-high Kupffer cell population is enriched within fibrotic regions of mouse MASLD liver, we next asked whether Cyclin D2 expression is similarly associated with senescence-associated macrophage states in human liver disease. We analyzed a single-cell RNA-sequencing dataset spanning healthy liver, cirrhotic liver, and hepatocellular carcinoma (HCC) to examine Cyclin D2 expression across hepatic immune cell populations (Fig. 8A-C). We first assessed the expression of disease-associated macrophage and senescence-associated genes across the myeloid compartment. Feature plots demonstrated expression of TREM2, SPP1, CDKN1A, and CCND2 within distinct macrophage populations predominately from cells corresponding to cirrhotic tissue (Fig. 8C, D). Notably, *TREM2* is a component of our macrophage senescence signature, and its co-expression of *CDKN1A* and *CCND2* identified macrophage populations exhibiting features of the senescence-associated state in human liver disease (Fig. 8D).

**Figure 8.**
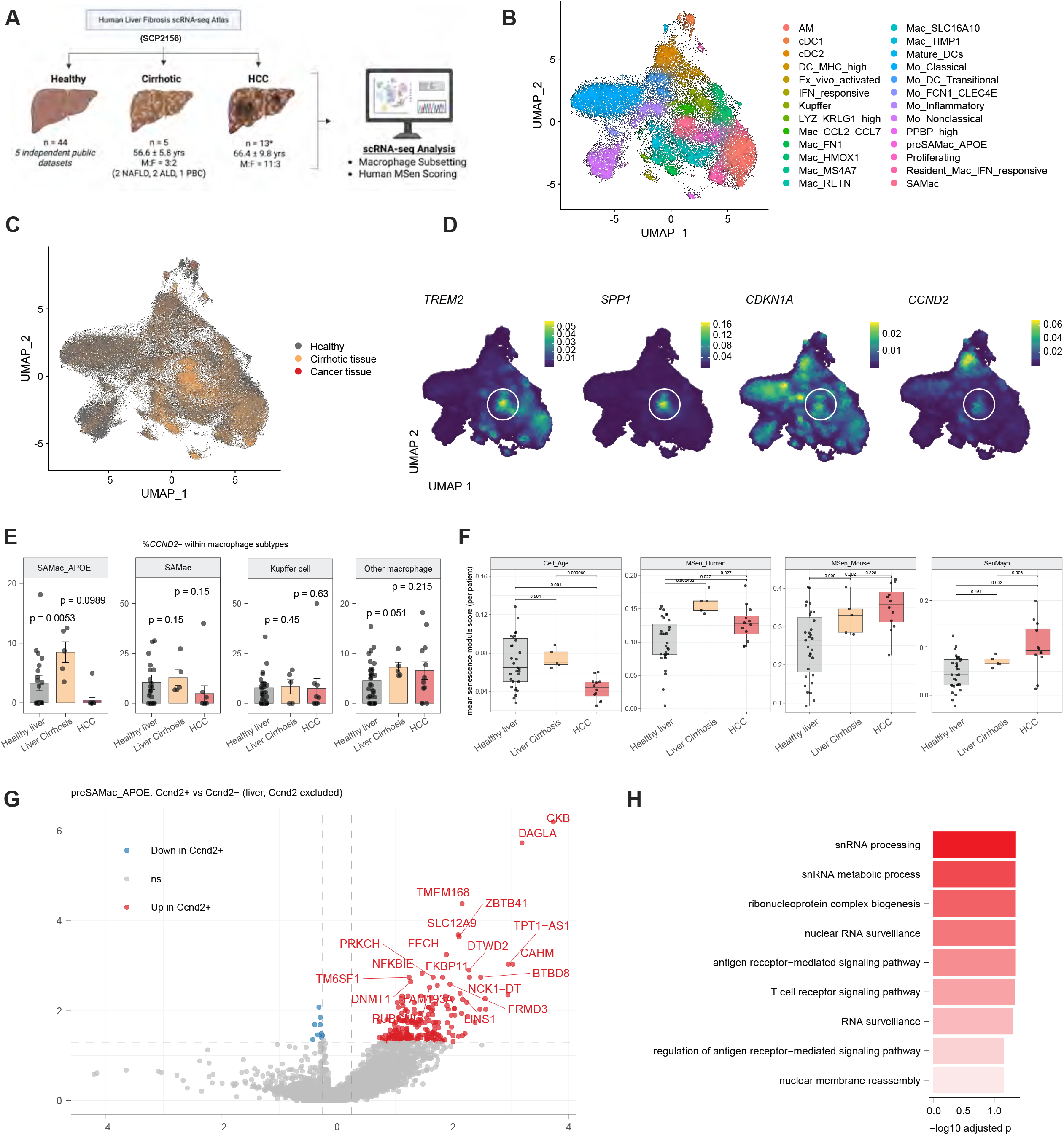
Identification of p21⁺Cyclin D2⁺ scar-associated macrophages in cirrhotic human liver. (A) Schematic of the human liver fibrosis scRNA-seq Atlas used to investigate *CCND2* and *CDKN1A* expression across healthy liver, cirrhotic liver, and hepatocellular carcinoma (HCC). The healthy liver cohort comprises 44 patients integrated from five independent public datasets, while the cirrhotic and HCC cohorts comprise 5 and 13 patients, respectively, from separate datasets within the Atlas. Created with BioRender.com. (B) UMAP visualization of macrophage and immune cell populations identified in the human liver scRNA-seq dataset. (C) UMAP visualization of the human liver scRNA-seq dataset colored by disease state, with healthy cells shown in gray, cirrhotic tissue cells in orange, and cancer tissue cells in red. (D) Feature plots showing expression of *TREM2*, *SPP1*, *CDKN1A*, and *CCND2* across human liver macrophage and immune cell populations. (E) Quantification of the percentage of *CCND2*-positive cells within SAMac_APOE, SAMac, Kupffer cell, and other macrophage populations across healthy liver, cirrhotic liver, and HCC. (F) Senescence module scores across healthy liver, cirrhotic liver, and HCC for four established senescence signatures: Cell Age, MSen(Human), MSen(Mouse), and SenMayo. (G) Volcano plot showing differentially expressed genes between *CCND2*^+^ and *CCND2*^−^ preSAMac_APOE macrophages from human liver, with *CCND2* excluded from the analysis. (H) Gene Ontology analysis of genes differentially expressed between *CCND2*^+^ and *CCND2*^−^ preSAMac_APOE macrophages, highlighting enrichment of RNA processing, ribonucleoprotein biogenesis, antigen receptor signaling, and nuclear RNA surveillance pathways. *P* values are shown in the figure.

We next quantified the fraction of CCND2^+^ cells within individual macrophage populations across healthy liver, cirrhosis, and HCC. CCND2 expression was detected across multiple macrophage states, including Kupffer cells, scar-associated macrophages (SAMacs), and the APOE-expressing pre-SAMac population (SAMac_APOE), but was particularly enriched within the SAMac_APOE population in cirrhotic liver (Fig. 8E). Because SAMac_APOE cells represent a pre-SAMac population, their enrichment for CCND2 in cirrhotic liver raises the possibility that Cyclin D2 expression is associated with an earlier disease-associated macrophage state that precedes or contributes to the emergence of mature scar-associated macrophages. However, the cross-sectional nature of these data does not establish a temporal or lineage relationship between these populations.

To assess whether *CCND2*^+^ macrophages exhibited broader transcriptional features of cellular senescence, we calculated four independent senescence module scores: Cell_Age, Msen_Human, Msen_Mouse, and SenMayo, across macrophage populations from healthy liver, cirrhosis, and HCC (Fig. 8F). These analyses revealed variation in senescence module scores across macrophage populations, with disease-associated macrophage populations in cirrhotic liver exhibiting increased senescence-associated scores across several signatures. Together with the enrichment of *CCND2* within the preSAMac_APOE population, these findings support an association between Cyclin D2 expression and senescence-associated disease macrophage states in human cirrhotic liver.

Focusing on the preSAMac_APOE population, we sought to determine whether *CCND2* expression was associated with distinct transcriptional features within this disease-associated macrophages. To do so, we compared the transcriptional profiles of *CCND2*^+^ and *CCND2*^−^ preSAMac_APOE cells, while excluding *CCND2* itself from the analysis. Differential expression analysis revealed a distinct transcriptional profile in *CCND2*^+^ cells, with gene ontology analysis identifying enrichment of RNA regulatory and nuclear processes, including RNA surveillance, snRNA processing, ribonucleoprotein complex biogenesis, and nuclear RNA surveillance, as well as pathways related to antigen receptor signaling (Fig. 8G). Notably, the enrichment of RNA- and ribonucleoprotein-associated processes parallels the translational and RNA-processing programs identified in our murine senescent macrophage models.

Interestingly, *CCND2* expression was also detected in several dendritic-cell populations, allowing us to examine whether Cyclin D2-associated transcriptional programs were shared across immune cell types (Supplementary Fig. 8A). *CCND2* expression was enriched in cDC2, mature dendritic cells, and transitional monocyte-derived dendritic cells. In cDC2 cells, *CCND2*+ cells were enriched for ribosome biogenesis, ribonucleoprotein complex biogenesis, RNA processing, rRNA metabolism, and DNA metabolic processes (Supplementary Fig. 8B). Several of these pathways overlapped with the RNA- and ribosome-associated programs identified in *CCND2*^+^ preSAMac_APOE macrophages and in our senescent macrophage models. In mature dendritic cells, *CCND2*^+^ cells were instead enriched for positive regulation of cell-cycle transition and G1/S progression, consistent with the canonical role of Cyclin D2 in cell-cycle regulation (Supplementary Fig. 8C). In transitional monocyte-derived dendritic cells, *CCND*^+^ cells were enriched for leukocyte and lymphocyte activation, T-cell activation, and MHC class II complex assembly and antigen presentation, consistent with immune activation and antigen-presenting functions (Supplementary Fig. 8D).

To determine whether the *CCND2*-associated programs observed in dendritic cells were accompanied by broader senescence-associated features, we compared senescence module scores between *CCND2*^+^ and *CCND2*^−^ dendritic cells. Across cDC2, DC_MHC_high, and transitional monocyte-derived dendritic cells, the association between *CCND2* expression and senescence scores varied across the independent signatures, with no consistent enrichment of the Msen_Human signature in *CCND2*^+^ cells. Although *SenMayo* scores were higher in *CCND2*^+^ cells in some populations, these differences were not consistently reproduced across scoring approaches or donor-level comparisons (Supplementary Fig. 8E-G).

These findings identify a Cyclin D2-enriched preSAMac_APOE population in cirrhotic human liver that exhibits distinct transcriptional features and broader senescence-associated programs. At the same time, examination of Cyclin D2 expression across other immune cell types revealed that similar RNA- and ribosome-associated programs can occur in *CCND2*^+^ dendritic cells without a consistent broader senescence signature, while other dendritic-cell populations associate Cyclin D2 with cell-cycle or immune activation programs. In conclusion, these data suggest that Cyclin D2 can engage distinct cellular programs depending on cell type and cellular context, supporting a context-dependent role for its non-canonical functions in senescent macrophages.

## Discussion

In this study, we show that macrophage senescence in response to genotoxic stress is controlled by a p53–p21-dependent program that not only sustains cell-cycle arrest and viability but also actively represses *Cdkn2a* (p16), establishing a p21-high, p16-low senotype rather than the canonical p16-high state observed in many other contexts^8–10^. This finding positions the p53–p21 pathway as a critical fate-determining node, governing whether a macrophage undergoes apoptosis following genotoxic stress or instead adopts a senescence program. This active suppression of p16 suggests that its downregulation is not a passive bystander effect of macrophage senescence, but the target of a regulatory program that may facilitate cell survival, particularly during the acute DNA damage response, perhaps with other factors driving a p16+ or p21+ senotype. Because macrophage senescence may occur as a result of diverse stress responses beyond genotoxic stress, including cholesterol loading^6,36^, chronic infection and mitochondrial dysfunction, an important open question is whether this same p53–p21-dependent repression of p16 generalizes across other senescence-inducing stimuli or is specific to the DNA damage response, macrophage ontogeny, and tissue type. In our prior study^6^, we observed that *in vitro* acetylated LDL cholesterol loading similarly drives a p21-high, p16-low senotype, suggesting this regulatory logic may extend beyond the DNA damage response. In support of this, in our manuscript we observed both p21+ and Ccnd2+ macrophages not only in aging livers, but also in mice fed a high-fat, high-cholesterol diet. However, *in vivo* analysis of murine *Cdkn2a* (p16) transcripts and protein expression in senescent cells via immunofluorescence, single-cell and spatial transcriptomics has proven difficult in our hands as well as others^6,35^.

Beyond cell-cycle arrest, senescent macrophages must also meet the substantial bioenergetic and biosynthetic demands of the SASP and other energetically costly functions. We find that Cyclin D2 is induced downstream of the p53–p21 pathway and, rather than promoting cell-cycle progression, undergoes nuclear-to-cytoplasmic redistribution during senescence, where it promotes AKT1-mTORC1 and other kinase signaling pathways. Cyclin D2 physically and functionally associates with MIC60 to support mitochondrial respiration, glycolysis, lipid droplet-mitochondria association, and perinuclear mitochondrial organization, and we hypothesize that this metabolic remodeling is in turn required to sustain the SASP. The relationship between Cyclin D2-MIC60 signaling and mTORC1 is an important open question raised directly by our data: Ccnd2 KO control macrophages already exhibit smaller, shorter mitochondria prior to senescence induction, raising the possibility that these cells enter senescence with a pre-existing bioenergetic deficit that limits their capacity for the metabolic rewiring senescence normally requires, potentially establishing a feed-forward relationship in which impaired mitochondrial architecture and reduced mTORC1 signaling reinforce one another.

This pathway’s relevance extends beyond our *in vitro* model. As discussed above, Cyclin D2 and p21 co-accumulate in macrophages during normal aging across multiple tissues and track with disease progression in diet-induced MASLD across genetically distinct mouse strains. Transcriptional analyses that distinguish Cyclin D2+ p21+ macrophages are enriched for cytokine activity and MMP expression, consistent with the SASP and tissue remodeling we describe *in vitro*, suggesting this population represents the *in vivo* counterpart of the senescent macrophage state we define mechanistically. Consistent with our prior work, senolytic clearance of this p21+ Cyclin D2+ senescent macrophage population with ABT-263 reduced Msen-high macrophages within fibrotic regions of MASLD liver, and Cyclin D2 and p21 similarly mark a disease-associated scar-macrophage state in human liver cirrhosis. Together, these findings indicate that the p53–p21–Cyclin D2 pathway in senescent macrophage populations represents a tractable therapeutic target across species and disease contexts.

Independent of macrophage biology, D-type cyclins have recently emerged as recurrent, non-canonical effectors of senescence-associated inflammation in other cell types. In human fibroblasts and aged mouse hepatocytes, Cyclin D1 (*Ccnd1*) partners with CDK6 to promote persistent DNA damage and cytoplasmic chromatin fragment (CCF) accumulation, thereby activating cGAS-STING signaling to sustain interferon-stimulated gene and SASP expression^16^. Pharmacological CDK4/6 inhibition attenuates this program and improves physical function in aged mice. A parallel study similarly identified a CDK4/6-NF-κB-RARα transcriptional complex that sustains SASP expression during aging and following chemotherapy^37^. Our findings, together with these studies, establish D-type cyclins as recurrent regulators of senescence-associated inflammation that act through distinct mechanisms such as metabolic reprogramming, genome instability, and innate immune signaling, raising the question of whether other cell cycle regulators, in senescent macrophages or other senescent cell types, are instead repurposed toward distinct non-canonical functions uncoupled from cell cycle progression.

## Methods

### Ethical compliance for mouse experiments

All animal experiments and procedures were approved by the University of California, Los Angeles (UCLA) Institutional Animal Care and Use Committee (IACUC) and conducted in accordance with all applicable institutional guidelines and regulations.

### Mice

All mice were maintained on a C57BL/6 background and housed in the University of California, Los Angeles (UCLA) animal facilities in the Center for Health Sciences under specific pathogen-free conditions with ad libitum access to standard laboratory chow and water. Wild-type (WT) C57BL/6J mice (JAX, stock no. 000664), *Trp53*-deficient mice (B6.129S2-*Trp53*^tm1Tyj^/J; JAX, stock no. 002101)^38^, *Cdkn1a* (p21)-deficient mice (B6.129S6(Cg)-*Cdkn1a*^tm1Led^/J; JAX, stock no. 016565)^39^, *Cdkn2a* floxed mice (*Cdkn2a*^tm^^1.1Gsu^/J; JAX, stock no. 036933)^40^, and *Lyz2*-Cre mice (B6.129P2-*Lyz2*^tm1(cre)Ifo^/J; The Jackson Laboratory, stock no. 004781)^41^ were obtained from The Jackson Laboratory and subsequently bred and maintained at UCLA. Whole-body *Ccnd2* knockout mice were kindly provided by the Jan Nolta Laboratory at the University of California, Davis.

Male mice aged 8-12 weeks were used for bone marrow isolation and subsequent generation of bone marrow-derived macrophages (BMDMs) for experiments comparing WT and *Trp53*, *Cdkn1a* (p21), *Ccnd2*, or *Cdkn2a* (p16)-deficient macrophages. For aging studies, young (3-4 months) and aged (21-25 months) male C57BL/6J mice obtained from The Jackson Laboratory and the National Institute on Aging were used. Bulk liver tissue from young and aged mice was used for gene expression analysis by quantitative PCR, while liver sections from the same cohorts were used for immunofluorescence analysis of CD68, p21, and Cyclin D2 expression.

### Murine Primary Macrophage Cultures

Bone marrow-derived macrophages (BMDMs) were generated from femurs of euthanized 8-12-week-old male C57BL/6J mice as previously described ^42^. Bone marrow cells were isolated using a mortar and pestle crushing method and cultured in macrophage growth medium (MGM) consisting of complete RPMI (RPMI-C; RPMI supplemented with 10% fetal calf serum, penicillin-streptomycin, 1 mM sodium pyruvate, 2 mM L-glutamine, 10 mM HEPES, and 50 μM 2-mercaptoethanol) supplemented with 25% L929 cell-conditioned medium as a source of macrophage colony-stimulating factor (M-CSF). Cells were differentiated for 7 days, with additional MGM supplementation on day 5.

On day 7, fully differentiated BMDMs were detached using cold phosphate-buffered saline (PBS) containing 5 mM EDTA, replated in 60/40 medium consisting of 60% RPMI-C and 40% MGM, and allowed to adhere overnight. On day 8, BMDMs were exposed to 10 Gy ionizing radiation (IR) to induce senescence, while control cells were mock irradiated. Following irradiation, cells were maintained for 10–12 days, with media replaced every 2-3 days. Mock-irradiated control macrophages were cultured in parallel and passaged every 2-3 days. All experiments involving senescent macrophages were performed 10-12 days following IR-induced DNA damage, as previously described^42^.

### Immunoblot analysis and quantification

Whole-cell lysates from murine bone marrow-derived macrophages (BMDMs) were prepared using radioimmunoprecipitation assay (RIPA) buffer supplemented with Halt protease and phosphatase inhibitor cocktail (1:100; Thermo Fisher Scientific). Protein concentrations were determined using a bicinchoninic acid (BCA) protein assay kit (Thermo Fisher Scientific). Equal amounts of total protein (10-20 μg) were denatured in sodium dodecyl sulfate (SDS) sample buffer, separated by SDS-polyacrylamide gel electrophoresis using 4-20% polyacrylamide gels, and transferred onto polyvinylidene difluoride (PVDF) membranes.

Membranes were blocked in 5% non-fat milk for 1h at room temperature and subsequently incubated with primary antibodies followed by appropriate secondary antibodies. Primary antibodies were generally used at a dilution of 1:1,000 unless otherwise indicated. The following primary antibodies were used: anti-β-tubulin, anti-phospho-histone H2AX (γH2AX), anti-p21 (1:200), anti-p16 (1:500), anti-Cyclin D2, anti-MIC60 (1:500), anti-phospho-4E-BP1 (Thr37/46), and anti-total 4E-BP1 (see Supplementary Table 2 for antibody manufacturer and catalog information).

Immunoblot images were quantified using ImageJ (v.1.50i). Band intensities were measured by densitometric analysis following background subtraction and normalized to the corresponding β-tubulin loading control.

### Annexin V/propidium iodide apoptosis assay and quantification

Apoptosis and cell death were assessed by Annexin V and propidium iodide (PI) staining followed by flow cytometry. Both adherent and non-adherent cells were collected to ensure inclusion of apoptotic and dead cells. Briefly, conditioned media containing non-adherent cells were collected and centrifuged for 5 min at 400 × *g*. Adherent cells were subsequently detached using cold PBS containing EDTA and combined with the corresponding non-adherent cell fraction. Cells were washed twice with cold PBS and resuspended in 100 μL of 1× Annexin V binding buffer.

Cells were stained with 5 μL Annexin V (Thermo Fisher Scientific, cat. no. A13201) and 1 μL of a 100 μg/mL PI working solution and incubated for 15 min at room temperature in the dark. Following incubation, 400 μL of 1× Annexin V binding buffer was added to each sample. Samples were filtered to obtain a single-cell suspension and subsequently analyzed by flow cytometry. Unstained and single-stained controls were included for compensation and gating.

Flow cytometry data were analyzed using FlowJo software. Cells were classified as viable (Annexin V^−^/PI^−^), early apoptotic (Annexin V^+^/PI^−^), late apoptotic (Annexin V^+^/PI^+^), or necrotic (Annexin V^−^/PI^+^) based on Annexin V and PI staining. Total dead cells were quantified as the combined percentage of early apoptotic, late apoptotic, and necrotic cells.

### Cell number quantification

To quantify cell number following irradiation-induced senescence, bone marrow-derived macrophages (BMDMs) were collected 10 days following irradiation and enumerated using a Moxi automated cell counter (ORFLO Technologies). Cells were identified and quantified based on size-defined gating to exclude cellular debris and non-cellular particles.

### Synthetic guide RNA design

Synthetic single-guide RNAs (sgRNAs) targeting exon 2 of *Cdkn1a* and *Ccnd2* and exon 1 of *Ccnd1* were designed using the Synthego Knockout Guide Design tool. The two highest-ranked guide sequences predicted by the design algorithm were synthesized by Synthego, and editing efficiency was evaluated using Synthego’s Inference of CRISPR Edits (ICE) analysis platform and immunoblotting. Guides achieving a KO score of ≥50 were selected for subsequent experiments. Protein depletion following CRISPR-mediated gene editing was further validated by immunoblotting using antibodies listed in Supplementary Table 2. A negative control sgRNA targeting the *Rosa26* locus was also obtained from Synthego. The sgRNA sequences used were as follows: *Cdkn1a*, 5′-ACAGGCACCATGTCCAATCC-3′; *Ccnd2*, 5′-TGAAGAAGAGGTCTTTCCTC-3′; and *Ccnd1* guide 1, 5′-GCACAGGAGCUGGUGUUCCA-3′.

### CRISPR-Cas9 ribonucleoprotein editing

CRISPR-Cas9 ribonucleoprotein (RNP) editing of macrophage precursors isolated from murine bone marrow was performed as previously described^43^. Synthetic sgRNA (Synthego, 1.5 nmol) and Alt-R Cas9 Electroporation Enhancer (IDT, 10 nmol) were combined in nuclease-free water in a 1.5-ml tube. In a separate tube, SpCas9-NLS (Macrolab, 40 pmol) was diluted 1:5 in nuclease-free water. Diluted SpCas9-NLS was subsequently combined with the sgRNA/electroporation enhancer mixture at a 1:1 ratio and incubated for 10 min at room temperature to form RNP complexes.

On day 3 of differentiation, bone marrow-derived monocytes were electroporated using a Neon NxT Electroporation System (Invitrogen) with the following parameters: 1,900 V, 20 ms, and 1 pulse. Following electroporation, cells were transferred to 1.5-ml tubes containing pre-warmed macrophage growth medium and allowed to recover for 1 h at 37 °C and 5% CO₂ before subsequent collection and downstream culture.

### RNA Isolation and RT-qPCR

Total RNA was isolated from murine bone marrow-derived macrophages (BMDMs) using RNA STAT-60 (Amsbio) according to the manufacturer’s instructions. For *in vivo* experiments, total RNA was extracted from flash-frozen liver tissue using RNA STAT-60 (Amsbio) and a TissueLyser system (Qiagen). Approximately 2 μg of total RNA was reverse transcribed into complementary DNA (cDNA) using the High-Capacity cDNA Reverse Transcription Kit (Applied Biosystems) according to the manufacturer’s instructions.

Gene expression was quantified by real-time quantitative PCR (RT-qPCR) using a CFX384 Real-Time PCR Detection System (Bio-Rad) and Maxima SYBR Green/ROX qPCR Master Mix (2X; Thermo Scientific). Relative gene expression was calculated using the 2^−ΔΔCt^ method and normalized to hypoxanthine phosphoribosyltransferase (*Hprt*). Gene-specific primer sequences are provided in Supplementary Table 1.

### Senescence-Associated β-galactosidase Assay

Senescence-associated β-galactosidase (SA-β-gal) activity was assessed using the Senescence β-Galactosidase Staining Kit (Cell Signaling Technology, cat. no. 9860S) according to the manufacturers’ instructions. Cultured murine cells were fixed and incubated in SA-β-gal staining solution (pH 6.0) at 37 °C for 10-16 h. Following staining, cells were stored in glycerol for long-term preservation and subsequent imaging and analysis.

### SPiDER-gal Flow Cytometry Assay

Senescence-associated β-galactosidase activity in live cells was assessed using the SPiDER-β-gal fluorescent probe (Dojindo, prod. Code SG02). Cells were washed with cold phosphate-buffered saline (PBS) and collected using cold PBS containing EDTA. Following collection, cells were counted and aliquoted at 1 × 10^6 cells per sample. Cells were pelleted by centrifugation and resuspended in 1 mL of FACS buffer containing 20 μM SPiDER-β-gal. Cells were incubated at 37°C for 30 min, filtered immediately before acquisition, and analyzed by flow cytometry.

### Extracellular Flux Assays

Bone marrow-derived macrophages (BMDMs) were detached, counted, and resuspended in 60/40 medium (60% RPMI-C and 40% macrophage growth medium) at a density of 3.5 × 105 cells/mL. Cells were seeded at 100 μL per well in XF96 cell culture microplates and allowed to adhere overnight.

Oxygen consumption rate (OCR) and extracellular acidification rate (ECAR) were measured using an Xfe96 Extracellular Flux Analyzer (Seahorse Bioscience, North Billerica, MA). Mitochondrial stress tests were performed in Seahorse assay medium supplemented with 10 mM glucose, 2mM glutamine, and 1 mM pyruvate, whereas glycolytic stress tests were performed in Seahorse assay medium without glucose.

For mitochondrial stress tests, OCR measurements were obtained at baseline and following the sequential injection of 1 μM oligomycin, 1.5 μM FCCP, and 1 μM rotenone and 2 μM antimycin A/rotenone (Sigma). For glycolytic stress tests, ECAR measurements were obtained at baseline and following the sequential injection of 10 mM glucose, 1 μM oligomycin, and 50 mM 2-deoxy-D-glucose (2-DG; Sigma). OCR and ECAR values were normalized to cell number quantified by 1 µg/mL Hoechst staining imaged and analyzed on an Operetta High-Content Imaging System. ATP production rates derived from oxidative phosphorylation and glycolysis were calculated from extracellular flux measurements according to Desousa et al. ^44^.

### Palbociclib treatment

BMDMs were seeded in XF96 cell culture microplates and allowed to adhere for 4-6 h. Culture medium was then supplemented with 500 nM palbociclib (Selleckchem, PD-0332991), and cells were incubated for an additional 24 h before downstream extracellular flux analysis. Vehicle-treated control cells DMSO-containing medium per well. The 500 nM working concentration was selected based on preliminary dose-response experiments demonstrating sufficient induction of cell-cycle arrest within 24 h and representing the lowest concentration tested that produced this effect (500 nM, 1 μM, 2.5 μM, 5 μM and 10 μM).

### mTOR signaling analysis by immunoblotting

To assess mTOR signaling, bone marrow-derived macrophages (BMDMs) were depleted of macrophage colony-stimulating factor (M-CSF) by replacing 60/40 medium with complete RPMI (RPMI-C) lacking M-CSF for 6 h before whole-cell lysate collection. Whole-cell lysates were subsequently prepared and analyzed by immunoblotting as described above.

### Rapamycin treatment

For pharmacological inhibition of mTOR signaling, BMDMs were seeded in XF96 cell culture microplates and allowed to adhere for 4-6 h. Culture medium was then supplemented with 50 nM rapamycin, and cells were incubated for an additional 24 h before downstream extracellular flux analysis.

### BODIPY and GRP75 staining

Bone marrow-derived macrophages (BMDMs) were detached, counted, and resuspended in 60/40 medium (60% RPMI-C and 40% macrophage growth medium) at a density of 3.5 × 10^5 cells/mL. Cells were seeded at 100 μL per well in PhenoPlate 96-well microplates (Revvity, Prod. No. 6055302) and allowed to settle for 10 min at room temperature before transfer to a 37°C incubator for overnight adherence.

The following day, cells were fixed with 4% electron microscopy (EM)-grade paraformaldehyde (Electron Microscopy Sciences, Cat. No, 15710) in PBS for 15 min at room temperature and washed four times with PBS. Cells were maintained in PBS at 4°C overnight before subsequent staining.

Cells were permeabilized with PBS containing 0.1% Triton X-100 (Thermofisher, Cat. No., HFH10) and 0.05% sodium deoxycholate (Thermofisher, Cat. No. 89905) for 20 min at room temperature and washed three times with PBS. Cells were subsequently blocked with 5% Donkey serum in PBS for 30 min at room temperature. Cells were then incubated with directly conjugated anti-GRP75 antibody (1:300; see Supplementary Table 2 for antibody information) and BODIPY (1:1,000; Thermo Fisher Scientific, Cat. No., D3922) diluted in blocking buffer for 1h at room temperature protected from light. Following incubation, cells were washed three times with PBS and counterstained with DAPI in PBS before imaging.

### Immunofluorescence staining and image analysis

#### Cultured BMDMs

Bone marrow-derived macrophages (BMDMs) were detached, counted, and replated in 60/40 medium (60% RPMI-C and 40% macrophage growth medium) onto chamber slides at a density of approximately 1.5 × 10^5 cells per well and allowed to adhere overnight. Cells were washed once with phosphate-buffered saline (PBS), fixed with 4% paraformaldehyde for 10 min, and washed three times with PBS for 5 min each. Cells were blocked with 3% bovine serum albumin (BSA) in PBS for 1 h at room temperature and subsequently permeabilized with 0.3% Triton X-100 in PBS for 15 min. Following three washes with PBS, cells were incubated overnight at 4 °C in a humidified chamber with primary antibodies diluted in blocking buffer, including anti-Cyclin D2 (SC-452; 1:200), anti-p21 (1:200), and anti-MIC60 (1:200) (see Supplementary Table 2 for antibody manufacturer and catalog information).

The following day, cells were washed three times with PBS and incubated with appropriate fluorophore-conjugated secondary antibodies (Invitrogen Goat anti-Rabbit IgG Secondary Antibody, Alexa Fluor™ 647, Invitrogen Donkey anti-Rat IgG Secondary Antibody, Alexa Fluor™ Plus 488, and Invitrogen Goat anti-Rabbit IgG Secondary Antibody, Alexa Fluor™ 546) diluted 1:500 as well as with fluorophore-conjugated antibodies for Tom20 (1:200, CoraLite®594-conjugated TOM20 Polyclonal antibody) and BODIPY (Invitrogen BODIPY™ 493/503, 1 uM) in PBS for 1 h at room temperature in the dark. Cells were washed three additional times with PBS and mounted using mounting medium containing DAPI (VECTASHIELD Vibrance® Antifade Mounting Medium with DAPI). Slides were subsequently imaged as described below.

#### Liver tissue

Mouse livers from young (4 months) and aged (25 months) mice were collected and either fixed in formalin for paraffin embedding or embedded in Optimal Cutting Temperature (OCT) compound and stored at -80°C. Tissue samples were sectioned at 5 μm and mounted onto glass slides for immunofluorescence staining. For paraffin-embedded sections, slides were deparaffinized using SafeClear (Fisher Scientific, Cat. No., 23-314629) and rehydrated through a graded ethanol series. Antigen retrieval was performed at 95 °C in a 10 mM sodium citrate buffer containing 0.05% Tween-20. OCT-embedded sections were allowed to equilibrate to room temperature, fixed in cold acetone, and permeabilized in PBS containing 0.1% Triton X-100 and 1% BSA.

All tissue sections were blocked in PBS containing 5% BSA and 0.1% Triton X-100 and incubated overnight at 4°C with primary antibodies diluted 1:200 unless otherwise indicated. Slides were subsequently washed and incubated with appropriate fluorophore-conjugated secondary antibodies diluted 1:500 for 1 h at room temperature. Following washing, sections were mounted using VECTASHIELD antifade mounting medium containing DAPI (Vector Laboratories).

For liver immunofluorescence, primary antibodies included anti-CD68 (1:200), anti-p21 (1:200), and anti-Cyclin D2 (SC-452; 1:200). See Supplementary Table 2 for antibody information.

#### Imaging and image analysis

Cell images were acquired using a Zeiss LSM 880 Confocal Microscope with 40x objective (UCLA Broad Stem Cell Research Center, Microscopy Core). Liver tissue sections were imaged using an Axioplan 2 fluorescence microscope (Zeiss) equipped with 10X and 20X objectives and Axiovision Rel. 3.0 software (Zeiss). Tiled whole-tissue images were analyzed using QuPath (v.0.6.0). CD68^+^ macrophages and Cyclin D2^+^ and p21^+^ cells were quantified using QuPath’s cell detection and object classification tools, with thresholds manually set to quantify cells with fluorescence intensity greater than background staining.

#### Mitochondrial membrane potential imaging

Bone marrow-derived macrophages (BMDMs) were detached, counted, and resuspended in 60/40 medium (60% RPMI-C and 40% macrophage growth medium) at a density of 3.5 × 10^5 cells/mL. Cells were seeded at 100μL per well in 96-well PhenoPlate microplates (Revvity, Prod. No., 6055302) and allowed to adhere overnight. Cells were then incubated in phenol red-free RPMI (GIBCO, Cat. No., 11835030) containing 15nM tetramethylrhodamine ethyl ester (TMRE, Fisher Scientific T669), 100nM MitoTracker Green (MTG, Thermofisher, Cat. No., M7514), and 1 μg/mL Hoechst (Thermofisher, Cat. No., 33342) for 1 h at 37 °C. Cells were subsequently washed twice with phosphate-buffered saline (PBS, GIBCO, Cat. no., 14190-144) and maintained in phenol red-free RPMI supplemented with 25 mM HEPES (GIBCO, Cat. no., 15630-080) immediately before imaging.

FCCP (1.5μM, 15 min) was used as a positive control for mitochondrial depolarization. Cells were imaged immediately using an ImageXpress high-content imaging system, with six fields acquired per well. TMRE fluorescence intensity was quantified on a per-cell basis as a measure of mitochondrial membrane potential, while MitoTracker Green fluorescence was quantified as an indicator of mitochondrial mass.

### Metabolomics

#### Metabolite Extraction

Metabolites were processed according to existing protocols{38070508}. Cells were placed on dry ice then rinsed with ice-cold 150 mM NH4AcO, pH 7.3, twice. Ice-cold 80% methanol extraction buffer with 10 nM trifluoromethanosulfanate was applied, then cells were incubated at -80 °C for 15 min. Cell suspensions were collected and centrifuged for 10 min at 17,000g. Supernatants were collected and evaporated using a Nitrogen evaporator (Organomation). Evaporated extracts were stored at −80 °C until analysis.

#### LC-MS Analysis

Dried metabolite extracts were reconstituted in 100 µL of 1:1 dH₂O:acetonitrile (I), and 10µL of each reconstituted sample was injected for analysis. Chromatographic separation was performed using a Vanquish UHPLC system (Thermo Scientific) equipped with a SeQuant ZIC-pHILIC polymeric column (2.1 × 150 mm, 5 µm; EMD Millipore) maintained at 35 °C. Mobile phases consisted of 20 mM ammonium carbonate, pH 9.7 (A), and 100% I (B), delivered at 150 µL/min. The gradient increased from 20% to 80% A over 20 min, returned to 20% A between 20 and 20.5 min, and was maintained at 20% A through 28 min. The UHPLC was coupled to a Q Exactive mass spectrometer (Thermo Scientific) operated in polarity-switching mode at 3.2 kV with an MS1 resolution of 70,000. Metabolites were assigned on the basis of exact mass, retention time, and MS2 fragmentation patterns acquired at a normalized collision energy of 35. Relative metabolite abundance was determined by integrating MS1 extracted-ion chromatograms using Mzmine 2.

#### Statistical Analysis

PCA was performed using R v.4.5.2. Heatmaps were generated using the pheatmap v.1.0.13 package.

#### IP-MS sample processing

Samples were harvested and collected as previously described for similar pull-down mass spectrometry techniques^45^. Cell pellets were resuspended in an IP lysis buffer (50 mM Tris pH 7.5, 150 mM NaCl, 1 mM EDTA, 0.5% NP-40 with Protease/phosphatase inhibitor mix). Lysates were centrifuged at 13,000 g for 10 min at 4°C and clarified lysates were incubated with IP antibody (Cyclin D2, 3741S (1:100; See reference table 2 for antibody information)) overnight at 4°C with gentle rocking. The next day, the mixture of lysate and IP antibody were incubated with 40uL Protein G magnetic beads (Invitrogen 10003D) overnight at 4°C with gentle rocking. Samples were then placed on a magnetic rack (Invitrogen 12-321-D) to isolate pull-down targets. Samples were washed 3 times with IP lysis buffer at different NP-40 concentrations (0.1%, 0.05%, and 0.01%). A mild on-bead digestion was performed right after using LysC at a 1:50 (wt/wt) enzyme-substrate ratio (Wako) in 8M Urea Buffer (8 M Urea (Promega), 50 mM Tris pH 8, 1mM DTT) for 2 hours at 23C at 1200 rpm. Following this initial digestion, reduction and alkylation were performed consecutively using tris-(2-carboxyethyl) (TCEP) (10 mM final) and 2-chloroacetamide (40 mM final) for 30 minutes each at 23°C with shaking at 1100 rpm. Before overnight protein digestion, the 8 M Urea was diluted to 2M with 50 mM Tris-HCl (pH 8) to permit the activity of the proteolytic enzyme trypsin. Trypsin (Promega) and Lys-C (Wako) was added at a 1:100 (wt/wt) enzyme-substrate ratio and placed in a thermomixer at 23°C overnight with shaking at 1000 rpm. Following digestion, samples were acidified with 10% TFA to pH 2-3 and clarified by centrifugation at 18,000 g for 10 min at 4°C. Peptides were desalted using Oasis HLB cartridges (Waters) on a vacuum manifold. Cartridges were activated with 1 mL 80% acetonitrile (I)/0.1% TFA, equilibrated three times with 1 mL 0.1% TFA, and samples were loaded twice. Columns were washed three times with 1 mL 0.1% TFA and peptides were eluted with 800 µL 50% I/0.25% formic acid. Eluates were dried by vacuum centrifugation in a SpeedVac (Labconco) and later resuspended in 0.1% formic acid. A volume corresponding to approximately 500 ng of peptides was injected into the timsTOF HT mass spectrometer (Bruker Daltonics).

#### Phosphoproteomics sample processing

Samples were lysed in 8 M Urea (Promega) in LoBind tubes (Eppendorf) and stored on ice until sonication. Lysed samples were sonicated using a probe sonicator 5x for 10 seconds On and 10 seconds Off at 10% amplitude, and protein was quantified using a bicinchoninic acid assay (BCA). Approximately 200 µg of protein for each sample was used for further processing, starting with reduction and alkylation using a 1:10 sample volume of tris-(2-carboxyethyl) (TCEP) (10 mM final) and 2-chloroacetamide (40 mM final) for 30 minutes each at 23°C with shaking at 1100 rpm. Before protein digestion, the 8 M Urea was diluted 7-fold with 100 mM Tris-HCl (pH 8) to permit the activity of the proteolytic enzyme trypsin. Trypsin (Promega) and Lys-C (Wako) was added at a 1:100 (wt/wt) enzyme-substrate ratio and placed in a thermomixer at 23°C overnight (16 h) with shaking at 1000 rpm. Following digestion, samples were acidified with 10% TFA to pH 2-3 and clarified by centrifugation (18,000 × g, 10 min, 4°C). Peptides were desalted using Oasis HLB cartridges (Waters) on a vacuum manifold. Cartridges were activated with 1 mL 80% acetonitrile (I)/0.1% TFA, equilibrated three times with 1 mL 0.1% TFA, and samples were loaded twice. Columns were washed three times with 1 mL 0.1% TFA and peptides were eluted with 800 µL 50% I/0.25% formic acid. Peptides were dried by vacuum centrifugation in a SpeedVac (Labconco). Approximately 90% of the desalted peptides were used for phosphopeptide enrichment. First, Ti-IMAC beads (Resyn Biosciences) were aliquoted in a 1:2 (w/w) peptide:beads ratio and equilibrated three times with binding buffer (0.1 M glycolic acid in 80% acetonitrile (I), 5% TFA). Dried peptide samples were resuspended in 200 μL of binding buffer, added to the equilibrated beads, and incubated for 30 min at 23°C, 1200 rpm. The unbound fraction was discarded, and beads were washed sequentially once each with 200 μL of the binding buffer, wash buffer 1 (60% I, 1% TFA, 200 mM NaCl), wash buffer 2 (60% I, 1% TFA), and finally with LC-MS grade water. Enriched phosphopeptides were eluted twice by incubating the beads with 150 μL of 1% (v/v) ammonium hydroxide (Sigma) in LC-MS grade water (Fisher Scientific) for 10 min at 23°C, 1200 rpm. The eluted peptides were transferred to a new protein LoBind tube containing 50 μL of 10% (v/v) formic acid in LC-MS grade water. Both eluates were pooled, dried, and resuspended in 0.1% formic acid. A volume corresponding to approximately 500 ng of peptides was injected into the timsTOF HT mass spectrometer (Bruker Daltonics).

#### Mass spectrometry proteomics acquisition

Dried peptides were resuspended in 0.1% (v/v) FA in MS grade water (Fisher) and analyzed on a timsTOF HT mass spectrometer, paired with a Vanquish Neo UHPLC system. Mobile phase A consisted of 0.1% (v/v) FA in MS grade water (Fisher), and mobile phase B consisted of 0.1% (v/v) FA in 100% MS grade Acetonitrile (Fisher). The LC was operated in trap-and-elute mode, where the peptides were first trapped onto a PepMap Neo Trap column (5 mm, 100 Å pore size, 5 µm particle size) and then reversed-phase separated using gradients mentioned below on an Aurora Elite C18 reverse phase column (15 cm, 120 Å pore size, 1.7 µm particle size for captive spray, IonOptiks), kept at 60°C using a column oven for Bruker Captive Spray source (Sonation Lab Solutions), and ionized in a CaptiveSpray source (Bruker Daltonics) at 1800 V. For phosphoproteome analysis, the %B gradient used was 1% to 3% over 2 min, 3.5% in half a minute, 16% in the next 15 min, 30% in the next 17 min, 45% in the next 5 min, and 60% B for 1 min followed by an increase to 95% B over 4.5 min. Flow rates went from 0.7 µL/min (first 2 min) to 0.3 µL/min (38.5min) to 0.7 µL/min (0.5 min) and finally to 0.8 µL/min (last 4 min). The raw data was acquired in dia-PASEF mode with variable isolation window widths in the m/z vs ion mobility plane. For dia-PASEF acquisition, ions were accumulated and separated in the TIMS device over a mobility range of 0.60-1.60 Vs/cm². MS/MS spectra were acquired across a mobility range of 0.65-1.30 1/K0 and a mass range of 275.8-1188.8 m/z. Collision energies were linearly ramped as a function of ion mobility from 20 eV at 1/K0 = 0.72 to 59 eV at 1/K0 = 1.49. The estimated mean cycle time was 1.38 s. For IPMS analysis, the %B gradient used was: 5% to 35% over 37 min at 0.3 µL/min, then to 45% in the next 4 mins, and 60% in the next 1 min, followed by an increase to 95% B over 3 min. During the last 2 minutes, the flow rate was changed to 0.4 µL/min. Equal-size windows of 25 Da were designed with an overlap of 1 Da to maximize the precursors ion coverage for further MS/MS. Ions were accumulated and separated in the TIMS analyzer over a mobility range of 0.60-1.60 V·s/cm² using 100 ms accumulation and ramp times. DIA acquisition comprised one MS1 scan followed by twelve dia-PASEF scans covering a mass range of 100-1700 m/z and an ion mobility range of 0.60-1.50 1/K0. Collision energies were linearly ramped according to ion mobility from 20 eV at 1/K0 = 0.60 to 59 eV at 1/K0 = 1.60. The resulting cycle time was approximately 1.38 s.

#### Mass spectrometry proteomics data search and quantitative analysis

The raw files were processed with Spectronaut (Biognosys) with the directDIA+ (Deep) search algorithm, its in-silico derived DIA analysis. Carbamidomethylation (cysteine) was set as a fixed modification for database search. Acetylation (protein N-term), oxidation (methionine), and phosphorylation (serine, threonine, tyrosine; only for phosphoproteomics dataset) were set as variable modifications. Reviewed mouse protein sequences (UniProt) were used for spectral matching. The false discovery rates for the PSM, peptide, and protein groups were set to 0.01, and the minimum localization threshold for PTM was set to zero. For MS2-level area-based quantification, the cross-run normalization option was unchecked (normalization was performed later using Msstats), and the probability cutoff was set to zero for the PTM localization. Quantitative analysis was performed in the R statistical programming language (v.4.5.0)^46^. Initial quality control analyses, including inter-run clustering, correlations, principal component analysis (PCA), peptide and protein counts, and intensities were completed in R. Statistical analysis of phosphorylation and protein abundance changes between exposed and control samples were computed using the R package Msstats (4.12.1)^47^.For phosphoproteomics, all peptides containing the same set of phosphorylated sites were summarized together into phosphorylation site groups using Tukey’s Median Polish. For both phosphopeptide and protein abundance Msstats pipelines, Msstats performs normalization by median equalization, imputation was turned Off (set to FALSE), and statistical tests of differences in intensity between conditions were calculated using default settings in Msstats. Specifically, Msstats calculates log2 fold changes as the ratio of averaged (across replicates) protein or phosphopeptide intensities between conditions, uses a Student’s t-test for p-value calculation and the Benjamini-Hochberg method of FDR estimation to adjust p-values. For the IPMS experiment, all peptides mapping to the same proteins were summarized together using Tukey’s Median Polish approach and the described computational pipeline above, but additional steps were taken for the scoring of protein-protein interactions. Protein-protein interaction strength was scored using SAINTexpress (http://apostl.moffitt.org/). Protein interactions with a Bayesian false-discovery rate (BFDR) < 0.05 were selected as high-confidence protein-protein interactions and visualized with Cytoscape (ver 3.10.4).

#### Kinase activity analysis

Kinase activities were estimated using known kinase-substrate relationships as performed previously ^48^. Log_2_ fold changes (Log_2_FC) were calculated per phosphorylation site intensities between conditions, and these values were used to infer kinase activity states using prior knowledge networks of kinase-substrate interactions derived from the Omnipath database ^48^. A Z-test from the comparison of fold changes in phosphosite intensity measurements of the known substrates against the overall distribution of fold changes across the sample was performed to infer kinase activities as a -log10(p-value). This statistical approach has been previously shown to perform well at estimating kinase activities ^48–50^.

#### Transcriptomics

In vitro monocultures of macrophages were lysed using 1mL of RNAstat60 to preserve RNA then promptly stored in RNAase free tubes in -80°C. 200 uL of Chloroform was added onto samples and shaken vigorously then allowed to rest on ice 10 min to establish phase separation. After 10 min samples were spun for 15 min at 12,000 x g 4°C. 400 uL of aqueous phase was transferred into new RNAase free microcentrifuge tubes then subsequently mixed with 500 uL isopropanol, rested at RT then spun max speed to form an RNA pellet. Two 75% ethanol washes were performed and nanodrop quantifications to assess RNA quality control was performed. Any RNA product > 1.8 260/280 was used for RTqPCR gene amplification and validation (see protocol above) then remaining RNA was used for tape station and library preparation. High quality RNA at 5 ng/uL was submitted to UCLAs TCGB core for library preparation and sequencing. In short, Poly adenylated RNA was collected and prepared for sequencing at 2 x 100 PE reads at least 20 M reads per sample using the NovaX plus sequencer to generate .fastq files (accessible with GEO numbers below).

#### Transcriptomic analysis

Raw sequencing data in FASTQ format were quality-filtered and trimmed to remove adapter sequences and low-quality reads using SOAPnuke. High-quality reads were aligned to the mouse mm10 reference genome using STAR. Aligned reads were processed to generate gene-level count matrices using FeatureCounts. Downstream analyses were performed in R (v4.5.0) using DESeq2 (v1.48.2). Gene expression values were normalized and variance stabilized (VST) for downstream analyses. Differentially expressed genes (DEGs) were identified using parametric statistical testing, with *P* values adjusted for multiple comparisons using the false discovery rate (FDR) method. Genes with an adjusted *q* value <0.05 and an absolute log2 fold change >0.32 were considered differentially expressed. Heatmaps were generated using pheatmap (v1.0.12), and statistical analyses and data processing were performed using R. Unsupervised K-means clustering was used to identify transcriptional states and visualize expression patterns across conditions using heatmaps generated with pheatmap (v1.0.12). Gene Ontology and pathway enrichment were performed using Enrichr. Senescence-associated transcriptional programs were quantified using GSVA, generating sample-level enrichment scores across in vitro experimental conditions.

#### Generation of MASLD mouse model

For spatial transcriptomics a “humanized” hyperlipidemic mouse model for progressive MASH was used similar to previous studies ^51^. In short, C57BL/6J male mice carrying transgenes for human apolipoprotein E*3-Leiden and cholesteryl ester transfer protein (*CETP*-*APOE*3* Leiden) and were fed HFHCD (33% kcal from cocoa butter and 1% cholesterol; Research Diets, cat. no. D10042101) for 16 weeks.

#### Administration of ABT-263

Starting at week 12 of the HFHCD, mice were either treated with vehicle or ABT-263 diluted in 10% ethanol, 30% polyethylene glycol 400 (Sigma, cat. no. 807485) and 60% Phosal 50 PG (Medchem Express, cat. no. HY-Y1903). ABT-263 was administered by oral gavage at 50 mg per kg body weight per day for 7 consecutive days with a 2-week break or interval between another 7-day cycle. At the end of the study, mice were fasted for 4 h, beginning at 9:00 am and were killed at 1:00 pm.

#### CosMx molecular imaging sample preparation

Formalin-fixed, paraffin-embedded (FFPE) mouse liver tissue sections were prepared by the Translational Pathology Core Laboratory at the David Geffen School of Medicine at UCLA and processed for spatial molecular imaging using the CosMx Spatial Molecular Imager (SMI) platform according to the manufacturer’s RNA assay protocol and established protocols from the UCLA Comprehensive Liver Research Center [43]. Briefly, 5-µm tissue sections were mounted onto slides and baked overnight at 60°C in an oven positioned at a 45° angle. Sections were deparaffinized using sequential xylene and ethanol washes, followed by heat-induced antigen retrieval. Tissue sections were permeabilized using 3 µg/mL Proteinase K, treated with fiducial markers (0.001%), post-fixed, quenched, and blocked with sulfo-NHS-acetate.

In situ hybridization was performed overnight at 37°C using the CosMx mouse RNA assay. Subsequent cyclical probe hybridization, barcode decoding, imaging, and quenching were performed automatically on the CosMx SMI instrument. The assay included the CosMx Mouse Universal Cell Characterization RNA Panel supplemented with additional genes comprising the Msen_mouse_UCLA gene signature: *Itgb2, Acod1, Ccl12, Ccl7, Ccnd2, Ccr3, Cyp4f18, Fgl2, Grina, H2-Q4, H2aw, H2bc4, H4c9, Id1, Id2, Ier5, Ifit2, Iigp1, Impact, Irf1, Irf8, Ly6a, Mmp13, Plin2, Plk2, Plk3, Pmaip1, Rhob, Rnase6, Rsad2, Slc3a2, Sp100, Sp110, Sp140, Trem2, Trib1,* and *Usp18, Zfas1*. Following hybridization, sections underwent stringent formamide/saline-sodium citrate (SSC) washes. Nuclear and cellular segmentation markers, including DAPI, CD298, B2M, CD68, CD45, and PanCK, were then applied.

Prepared slides were assembled into flow cells and imaged using liver-specific pre-bleaching and cell-segmentation configurations. All fields of view were acquired on the CosMx SMI instrument, and image processing, transcript quantification, and downstream spatial analyses were performed using the AtoMx Spatial Informatics Platform.

#### Spatial transcriptomics data analysis and senescence detection

Cell segmentation was performed by CosMx SIPs (v2) default large + small cells option (option E) and iteratively optimized to eliminate artifacts, including abnormal cell gaps and nuclear-only boundaries, by adjusting segmentation parameters and workflows. Quality control was performed using AtoMx, excluding FOVs with ≤100 mean transcripts per cell and cells failing default thresholds for transcript counts, negative probes, complexity, or area. Cell number and size were also calculated in AtoMx. Seurat LogNormalize ^52^ was used for normalization, with UMAP and spatial visualization generated using DimPlot and ImageDimPlot. Cell types were assigned using InSituType, a likelihood-based cell typing for single cell spatial transcriptomics ^53^. Data analysis related to senescence detection was performed using R (v4.6.1). Senescence signature gene coverage was calculated by the genes identified in the data set over total genes in a gene list. Gene signatures with over 50% coverage were used for detection of senescent cells. For scalable single-cell gene signature scoring the mean Ucell score was calculated for every cell per mouse section ^54^. The top 90^th^ percentile of Msen_mouse_UCLA Ucell scorers were labeled as Msen_hi. Volcano plot analysis was performed using a pseudo bulk approach taking the average expression of genes within Msen_hi vs Msen_low macrophages per mouse. Fibrosis density is highlighted as a purple gradient and defined by gene expression density of ECM markers such as Col1a1*/Col3a1/Col1a2/Acta2/Timp1* in addition to the density of stromal cells.

#### Tabula Muris Senis aging dataset analysis

Publicly available single-cell RNA-sequencing data from the Tabula Muris Senis (TMS) atlas were analyzed to characterize age-associated changes in *Ccnd2* expression across macrophage populations and tissues. The TMS dataset was accessed through the Bioconductor TabulaMurisSenisData (v1.14.0) package as processed FACS (Smart-seq2) and droplet-based (10x Genomics) single-cell expression data. Macrophage-like populations were identified from cell ontology annotations using GrepL tools to identify cells matching these terms: macrophage, Kupffer cell, monocyte, microglial, and related myeloid classifications. *Ccnd2* expression was quantified from processed count and log-count matrices. Mice were categorized into young (1-3 months), middle-aged (18-21 months), and aged (24-30 months) groups. Analyses were performed across 15 tissues using pooled FACS and droplet datasets, with individual mice retained as biological replicates for statistical testing. Cell populations were required to contain at least 100 cells per age group, while trend analyses required at least 100 cells per mouse. Age-associated trends were evaluated using a generalized linear model, with age modeled continuously in months. Effect sizes were quantified as Cohen’s *d* comparing aged and young mice, calculated using the pooled standard deviation.

#### Characterization of murine macrophage states

To further characterize *Ccnd2*-associated macrophage states, aged Kupffer cells and splenic macrophages were stratified and double-positive cells ( *Ccnd2/Cdkn1a*+) were defined as cells with detectable expression of both genes. Differential expression between these populations was performed using Seurat’s (5.5.1) Wilcoxon rank-sum test. Genes with adjusted *P* < 0.05 and Log2 fold-change > 0.32 were considered differentially expressed. Gene Ontology Biological Process and Molecular Function enrichment was performed using *clusterProfiler* and *org.Mm.eg.db*, using all genes tested in the differential-expression analysis as the enrichment background and Benjamini–Hochberg correction for multiple testing.

#### Human myeloid single-cell RNA-sequencing analysis

Liver and lung scRNAseq data were obtained from the Single Cell Portal (SCP2156) as processed expression matrices with cell barcodes, gene features, metadata, and published UMAP coordinates. Data were imported into R (v4.5.0) using Seurat (v5.5.0), matched to metadata by cell barcode, and harmonized according to donor, tissue, disease state, and macrophage identity. Cell populations included preSAMac_APOE, SAMac, Kupffer, alveolar, and monocyte-derived macrophages, with HCC-adjacent liver samples classified according to donor provenance. Expression data were normalized using Seurat, followed by identification of the 2,000 most variable genes, scaling, PCA, and UMAP using the first 30 principal components. Graph-based clustering was evaluated across resolutions of 0.1–0.8. UMAPs were additionally used to visualize gene-expression density using Nebulosa. CCND2-positive cells were defined by detectable raw expression and quantified at the donor level as cell counts or percentages within defined macrophage populations. Senescence-associated signatures were scored using Seurat AddModuleScore with 100 control genes per signature, excluding CCND2 to prevent circularity. Differential expression was additionally assessed using donor-level average expression and visualized as volcano plots, with significant genes defined by adjusted P <0.05 and |fold change| >0.32. Gene Ontology Biological Process enrichment was performed using clusterProfiler and org.Hs.eg.db, using Benjamini-Hochberg correction and dataset genes as the enrichment background.

#### Statistics and reproducibility

Sample sizes and statistical analyses for *in vitro* experiments were determined before data collection. No data were excluded from the *in vitro* experiments. Unless otherwise indicated, *in vitro* experiments included at least three independent biological replicates. All *in vitro* experiments were independently repeated three times with similar results, where applicable. Transcriptomic, phosphoproteomic, metabolomic and Co-IP MS experiments included 3-6 biological replicates, with sample numbers determined by sample availability and sequencing or assay requirements.

For *in vivo* experiments, sample sizes were determined based on animal availability, experimental feasibility and prior studies with expected similar effect sizes; no formal power analyses were performed. Samples were excluded if mice developed tumors or became ill. No animals were excluded from analysis unless otherwise indicated. For Fig. 7A, five aged male C57BL/6J mice (22-24 months) and five young male C57BL/6J mice (3-4 months) were analyzed. For Fig. 7B, liver tissue from one aged male (24 months) and one young male (3 months) was analyzed as representative samples selected based on tissue availability, and these data were used for qualitative assessment.

For spatial transcriptomic analyses (Fig. 7I-P), data were re-analyzed from a previously generated humanized MASLD mouse model^6^. Three mice per condition were analyzed, with one tissue section from each mouse. Sample numbers were determined based on sequencing requirements and throughput limitations of the CosMx Spatial Molecular Imager. Differential gene expression analysis was performed using pseudobulked data, with gene expression aggregated across cells within each mouse; each mouse was therefore treated as an independent biological replicate.

No randomization or blinding was used. Statistical analyses were performed using GraphPad Prism 10.4.0 and R 4.5.2. Statistical tests are indicated in the corresponding figure legends. Unless otherwise stated, in vitro comparisons were performed using unpaired two-tailed *t*-tests or ordinary one-way ANOVA. Statistical significance was defined as *P* < 0.05. Individual data points are shown where appropriate.

## Data Availability

Phosphoproteomics proteomics, IPMS, and large computational datasets used here were uploaded to the Proteomics IDEntifications Database (PRIDE) at ebi.ac.uk/pride under project accession “PXD083659” . For reviewers, the token is “rn9hp0iqz2Bb”. All other data supporting the findings are available from the corresponding author upon request. Source data are provided with this paper.

**Supplementary Table 1.**
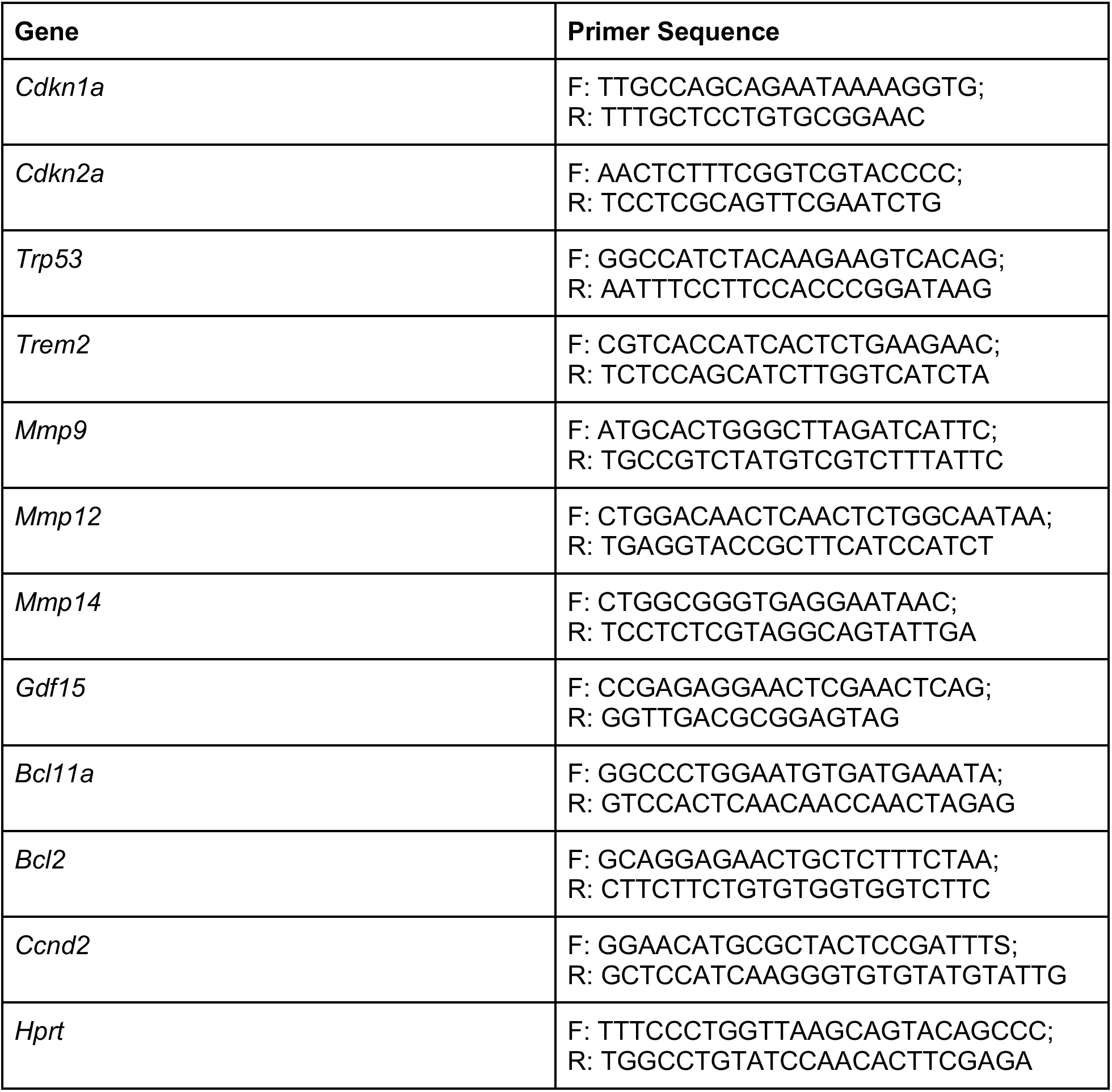
RT-qPCR primers and sequences.

**Supplementary Table 2.** Antibodies.

| Antibody target; Host | Manufacturer information |
| --- | --- |
| P-Histone γH2AX; Rabbit | Cell Signaling; Cat. no. 9718T |
| Cleaved Caspase 3; Rabbit | Cell Signaling; Cat. no. 9664S |
| β-Tubulin; Rabbit | Cell Signaling; Cat. no. 2146S |
| P21; Rabbit | Abcam; Cat. no. AB188224 |
| P16; Rabbit | Abcam; Cat. no. AB211542 |
| Cyclin D2; Rabbit | Cell Signaling; Cat. no. 3741S |
| p-4EBP1 (Thr37/46); Rabbit | Cell Signaling; Cat. no. 2855T |
| 4EBP1 (Total); Rabbit | Cell Signaling; Cat. no. 9644T |
| Mic60; Mouse | Abcam; Cat. no. 110329 |
| Tom20; Rabbit | Proteintech; Cat. No. 11802-1-AP |
| Cyclin D1; Rabbit | Cell Signaling; Cat. no. 2978T |
| Cd68; Chicken | Abcam; Cat. no. 318303 |
| GRP75-647; Mouse | AntibodiesInc; Cat. no. 75-127-FL650 |
| Cyclin D2; Rat | Santa Cruz Biotechnology; Cat. no. SC-452 |
| Anti-rat AF-647 | Invitrogen; Cat. no. A-21247 |
| Anti-rabbit AF-555 | Invitrogen; Cat. no. A-351572 |
| Anti-chicken AF-488 | Invitrogen; Cat. no. A-11039 |

Conceptualization – G.T. and A.J.C. | Methodology – G.T. and A.J.C. | Investigation – G.T. (all wet lab and in vivo experiments), I.A.S.P.(bioinformatic analysis), C.Y.D (tissue immunofluorescence experiments), I.A. (Senolytic testing in vitro and in vivo), G.T. (in vitro senescence experiments), A.C.A (in vitro senescence experiments), A.D (in vitro senescence experiments), Y.D. (Co-IP MS, Phosphoproteomics sample processing and analysis), G.Tong (in vitro senescence experiments and bioinformatic analysis), J.M.E (in vitro senescence experiments and bioinformatic analysis), L.E. (in vitro senescence experiments), W.R.A (Metabolomics analysis), A.S. (in vitro senescence experiments), I.C. (IF quantification), E.E. (bioinformatic analysis), P.K. (Co-IP MS Data Analysis) J.N. (Phosphoproteomic data analysis), K.C. (Spatial transcriptomic sample processing), C.P. (Spatial transcriptomic analysis), L.S. (Cellular Efflux Assay Data Analysis), C.B. (Mitochondrial imaging), S.T.H. (Ex vivo experiments for MASLD model), A.S.K. (Metabolomics), H.R.C. (Metabolomics), A.J.L (provided MASLD mice), T.E.O (Metabolomics), J.M. (bioinformatic analysis), R.S. (Liver Microscopy), V.A.T (bioinformatic analysis), O.S.S. (Mitochondrial studies), M.B. (Co-IP MS, Phosphoproteomics) | Writing – Original Draft, G.T. and A.J.C. | Writing – Review & Editing, all authors. | Supervision – A.J.C | Funding Acquisition – A.J.C.

## Acknowledgements

Research reported in this publication was supported by the National Institute of General Medical Sciences of the National Institutes of Health Maximizing Investigators’ Research Award (MIRA) award number 1R35GM156893. This project was supported by funds provided by the Glenn Foundation for Medical Research and American Federation of Aging Research, Junior Faculty Award (20225528), and the UCLA-UCSD Diabetes Research Center Award funded by the National Institute of Diabetes and Digestive and Kidney Diseases (NIDDK) grant P30DK063491, and also supported by the National Institute of Diabetes and Digestive and Kidney Diseases (NIDDK), the Office of Disease Prevention (ODP), the Office of Nutrition Research (ONR), the Chief Officer for Scientific Workforce Diversity (COSWD), and the Office of Behavioral and Social Sciences Research (OBSSR) of the National Institutes of Health under award number U24DK132746-01, UCLA LIFT-UP (Leveraging Institutional Support for Talented, Underrepresented Physicians and/or Scientists). G.T. is supported by the NIH Ruth L. Kirschstein National Research Service Award AI007323, T32 training grant 2T32AI007323-31. I.S.P. was supported by the National Institute of General Medical Sciences of the National Institutes of Health under Award Number T32GM145388. G.T. was supported bv a UCLA QCBio Collaboratory Fellowship . This work was also supported by UCLA Division of Life Sciences startup funds (V.A.T. and A.J.C). The content is solely the responsibility of the authors and does not necessarily represent the official views of the National Institutes of Health. While preparing this work, the authors used AI tools Grammarly, Claude, and ChatGPT to edit the text for grammar and clarity. After using this tool/service, the authors reviewed and edited the content as needed and are fully responsible for the content of the publication.

