## Supplemental Figures for "The p53–p21–Cyclin D2 regulatory axis drives metabolic reprogramming and a distinct senescent macrophage senotype during aging and MASLD"


### Supplementary materials and figures

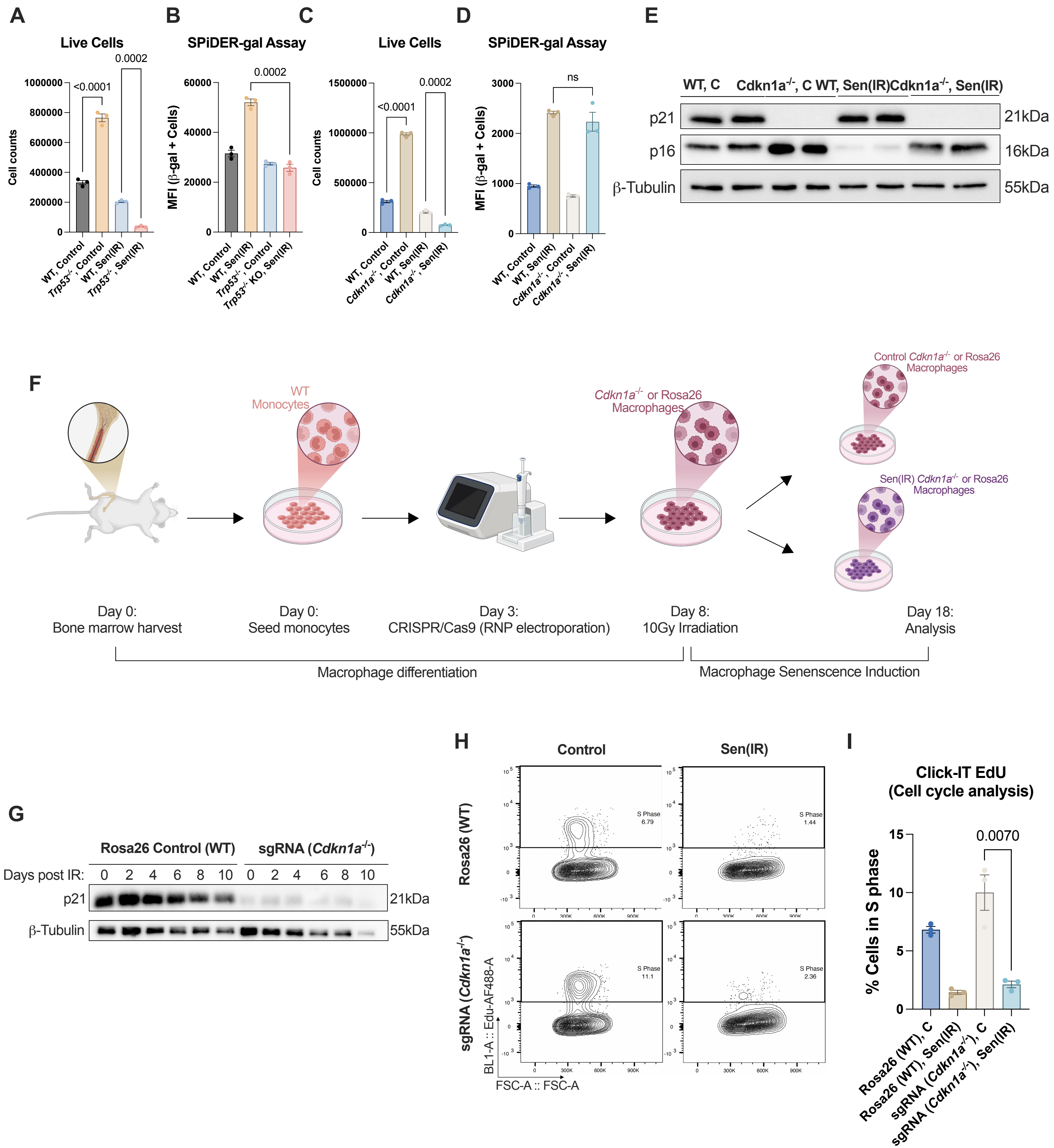


**Supplementary Figure 1. Validation of p53- and p21-dependent phenotypes and CRISPR/Cas9-mediated *Cdkn1a* deficiency in senescent macrophages.**

**Supplementary Figure 1. Validation of p53- and p21-dependent phenotypes and CRISPR/Cas9-mediated *Cdkn1a* deficiency in senescent macrophages.**

(A) Mean ± s.e.m. live cell counts in control and Sen(IR) WT and *Trp53*$^{-/-}$ BMDMs 10 days following irradiation. *P* value of one-way ANOVA with Tukey's multiple-comparisons test of *n* = 3 biological replicates. (B) Mean ± s.e.m. β-galactosidase activity, measured by SPiDER-β-gal staining, in control and Sen(IR) WT and *Trp53*$^{-/-}$ BMDMs. *P* value of unpaired two-tailed Student's *t*-test of *n* = 3 biological replicates. (C) Mean ± s.e.m. live cell counts in control and Sen(IR) WT and *Cdkn1a*$^{-/-}$ BMDMs 10 days following irradiation. *P* value of one-way ANOVA with Tukey's multiple-comparisons test of *n* = 3 biological replicates. (D) Mean ± s.e.m. β-galactosidase activity, measured by SPiDER-β-gal staining, in control and Sen(IR) WT and *Cdkn1a*$^{-/-}$ BMDMs. *P* value of unpaired two-tailed Student's *t*-test of *n* = 3 biological replicates. I SDS-PAGE gels and immunostaining (western blot) for p21 and p16 in control and Sen(IR) WT and *Cdkn1a*$^{-/-}$ BMDMs. Β-Tubulin was used as a loading control. (F) Experimental schematic for generation of *Cdkn1a*-deficient macrophages using CRISPR/Cas9 ribonucleoprotein (RNP) complexes. Bone marrow was harvested from WT mice, monocytes were seeded and differentiated into macrophages, and cells were electroporated with CRISPR/Cas9 RNP complexes containing guide RNAs targeting either *Cdkn1a* or the Rosa26 allele. Following macrophage differentiation, cells were exposed to 10 Gy IR on day 8 and analyzed on day 18. Cells analyzed 10 days following irradiation are designated senescent (Sen(IR)) macrophages. Created with BioRender.com. (G) SDS-PAGE gels and immunostaining (western blot) for p21 in Rosa26(WT) and sgRNA(*Cdkn1a*$^{-/-}$) macrophages at the indicated days following irradiation. Β-Tubulin was used as a loading control. (H) Representative flow cytometry plots of EdU incorporation in control and Sen(IR) Rosa26(WT) and sgRNA(*Cdkn1a*$^{-/-}$) macrophages. (I) Mean ± s.e.m. percentage of cells in S phase based on EdU incorporation in control and Sen(IR) Rosa26(WT) and sgRNA(*Cdkn1a*$^{-/-}$) macrophages. *P* value of unpaired two-tailed Student's *t*-test for the indicated comparison of *n* = 3 biological replicates.

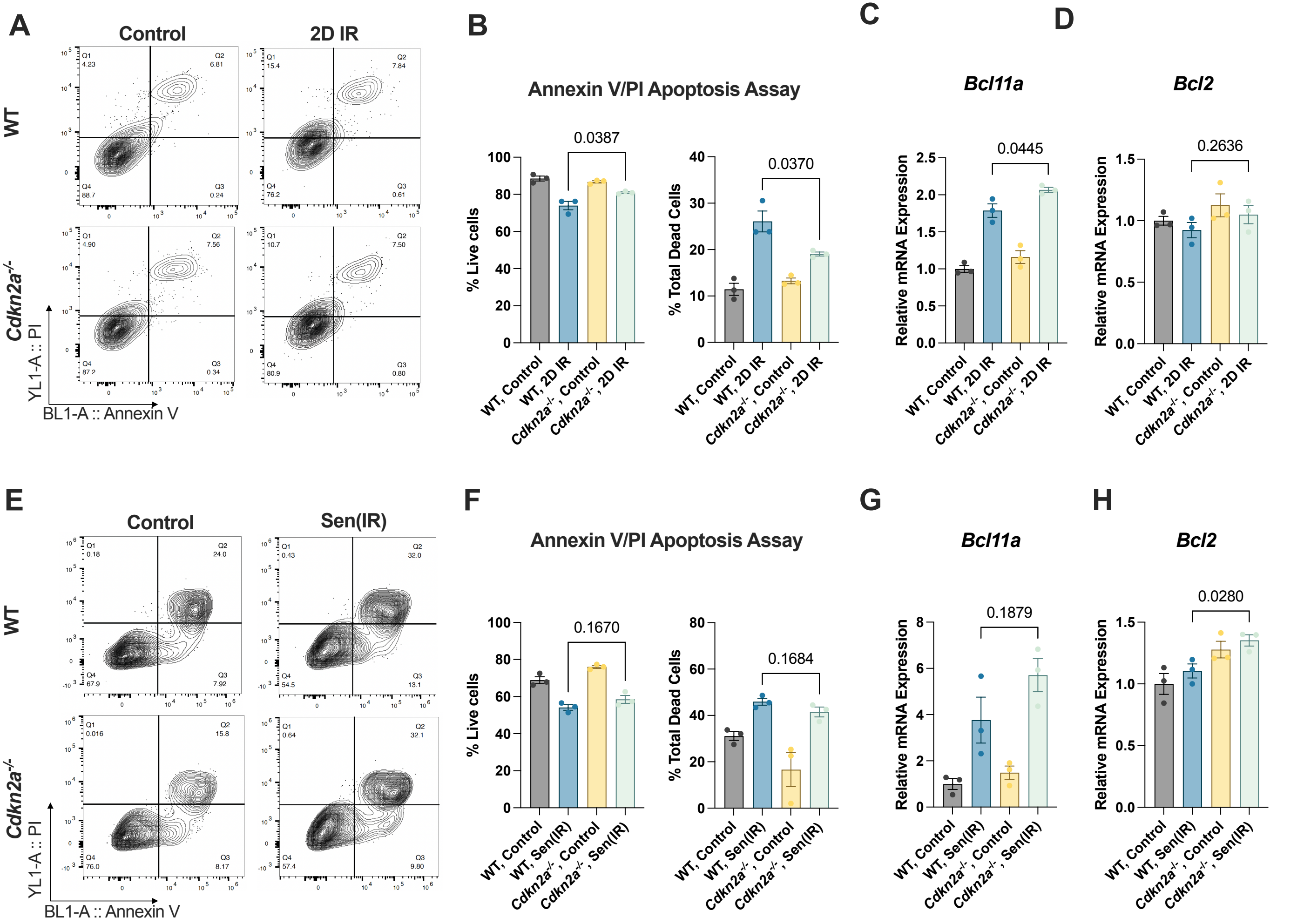


**Supplementary figure 2: Acute p16 downregulation supports cell survival as macrophages become senescent.**

**Supplementary Figure 2. Acute p16 downregulation supports cell survival as macrophages become senescent.**

(A) Representative flow cytometry plots of Annexin V and propidium iodide (PI) staining in control and 2D IR WT and *Cdkn2a*$^{-/-}$ macrophages. (B) Mean ± s.e.m. percentage of live (Q4) and total dead cells (Q1-3) from Annexin V/PI staining in control and 2D IR WT and *Cdkn2a*$^{-/-}$ macrophages. *P* value of unpaired two-tailed Student's *t*-test of *n* = 3 biological replicates. (C) Mean ± s.e.m. *Bcl11a* mRNA transcript levels, relative to control, in control and 2D IR WT and *Cdkn2a*$^{-/-}$ macrophages. *P* value of unpaired two-tailed Student's *t*-test of *n* = 3 biological replicates. (D) Mean ± s.e.m. *Bcl2* mRNA transcript levels, relative to control, in control and 2D IR WT and *Cdkn2a*$^{-/-}$ macrophages. *P* value of unpaired two-tailed Student's *t*-test of *n* = 3 biological replicates. I Representative flow cytometry plots of Annexin V and PI staining in control and Sen(IR) WT and *Cdkn2a*$^{-/-}$ macrophages. (F) Mean ± s.e.m. percentage of live (Q4) and total dead cells (Q1-3) from Annexin V/PI staining in control and Sen(IR) WT and *Cdkn2a*$^{-/-}$ macrophages. *P* value of unpaired two-tailed Student's *t*-test of *n* = 3 biological replicates. (G) Mean ± s.e.m. *Bcl11a* mRNA transcript levels, relative to control, in control and Sen(IR) WT and *Cdkn2a*$^{-/-}$ macrophages. *P* value of unpaired two-tailed Student's *t*-test of *n* = 3 biological replicates. (H) Mean ± s.e.m. *Bcl2* mRNA transcript levels, relative to control, in control and Sen(IR) WT and *Cdkn2a*$^{-/-}$ macrophages. *P* value of unpaired two-tailed Student's *t*-test of *n* = 3 biological replicates.

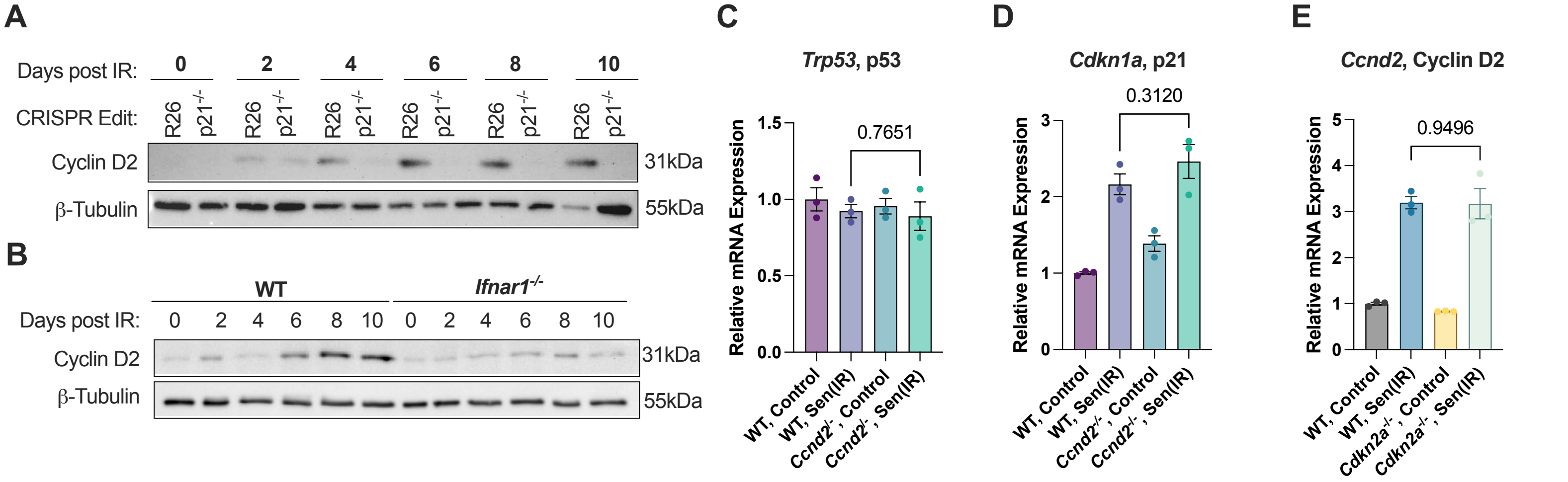


**Supplementary Figure 3. Cyclin D2 is regulated downstream of the p53-p21 axis, independently of p16, and is additionally modulated by Type I interferon signaling.**

**Supplementary Figure 3. Cyclin D2 is regulated downstream of the p53–p21 axis, independently of p16, and is additionally modulated by Type I interferon signaling.**

(A) SDS-PAGE gels and immunostaining (western blot) for Cyclin D2 in Rosa26(WT) and sgRNA(*Cdkn1a*$^{-/-}$) macrophages at the indicated days following irradiation. B-Tubulin was used as a loading control. (B) SDS-PAGE gels and immunostaining (western blot) for Cyclin D2 in WT and *Ifnar1*$^{-/-}$ macrophages at the indicated days following irradiation. B-Tubulin was used as a loading control. Mean ± s.e.m. *Trp53* (p53) mRNA transcript levels, relative to control, in WT and *Ccnd2*$^{-/-}$ macrophages following senescence induction. *P* value of unpaired two-tailed Student's *t*-test of *n* = 3 biological replicates. (D) Mean ± s.e.m. *Cdkn1a* (p21) mRNA transcript levels, relative to control, in WT and *Ccnd2*$^{-/-}$ macrophages following senescence induction. *P* value of unpaired two-tailed Student's *t*-test of *n* = 3 biological replicates. I Mean ± s.e.m. *Ccnd2* (Cyclin D2) mRNA transcript levels, relative to control, in WT and *Cdkn2a*$^{-/-}$ macrophages following senescence induction. *P* value of unpaired two-tailed Student's *t*-test of *n* = 3 biological replicates.

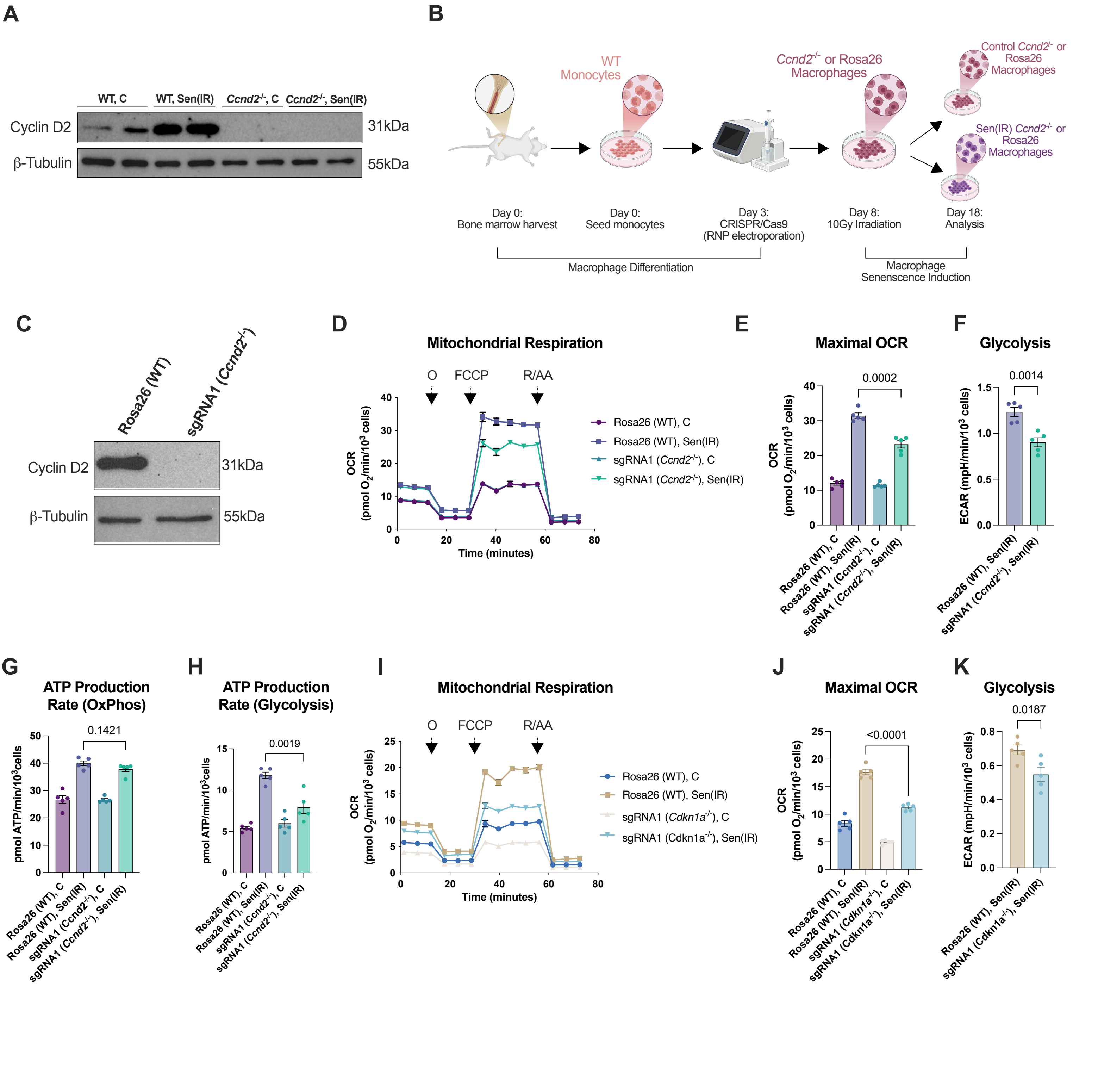


**Supplementary Figure 4. CRISPR/Cas9-mediated deletion validates metabolic phenotypes associated with *Ccnd2* and *Cdkn1a* deficiency.**

**Supplementary Figure 4. CRISPR/Cas9-mediated deletion validates metabolic phenotypes associated with *Ccnd2* and *Cdkn1a* deficiency.**

(A) SDS-PAGE gels and immunostaining (western blot) for Cyclin D2 in control and Sen(IR) BMDMs isolated from WT and *Ccnd2*$^{-/-}$ mice. B-Tubulin was used as a loading control. (B) Experimental schematic for generation of *Ccnd2*-deficient macrophages using CRISPR/Cas9 ribonucleoprotein (RNP) complexes. Bone marrow was harvested from WT mice, monocytes were seeded and differentiated into macrophages, and cells were electroporated with CRISPR/Cas9 RNP complexes containing guide RNAs targeting either *Ccnd2* or the Rosa26 allele. Following macrophage differentiation, cells were exposed to 10 Gy IR on day 8 and analyzed on day 18. Cells analyzed 10 days following irradiation are designated senescent (Sen(IR)) macrophages. Created with BioRender.com. (C) SDS-PAGE gels and immunostaining (western blot) for Cyclin D2 in Rosa26(WT) and sgRNA(*Ccnd2*$^{-/-}$) macrophages. B-Tubulin was used as a loading control. (D) Representative mitochondrial respiration profiles measured by extracellular flux analysis in control and Sen(IR) Rosa26(WT) and sgRNA(*Ccnd2*$^{-/-}$) macrophages. Oligomycin, FCCP, and R/AA were sequentially injected as indicated. I Mean ± s.e.m. maximal oxygen consumption rate (OCR) from technical replicates in the representative experiment shown in D. *P* value of unpaired two-tailed Student's *t*-test. (F) Mean ± s.e.m. glycolytic rate from technical replicates in the representative experiment shown in D. *P* value of unpaired two-tailed Student's *t*-test. (G) Mean ± s.e.m. ATP production rate through oxidative phosphorylation (OxPhos) from technical replicates in the representative experiment shown in D. *P* value of unpaired two-tailed Student's *t*-test. (H) Mean ± s.e.m. ATP production rate through glycolysis from technical replicates in the representative experiment shown in D. *P* value of unpaired two-tailed Student's *t*-test. (I) Representative mitochondrial respiration profiles measured by extracellular flux analysis in control and Sen(IR) Rosa26(WT) and sgRNA(*Cdkn1a*$^{-/-}$) macrophages. Oligomycin, FCCP, and R/AA were sequentially injected as indicated. (J) Mean ± s.e.m. maximal OCR from technical replicates in the representative experiment shown in I. *P* value of unpaired two-tailed Student's *t*-test. (K) Mean ± s.e.m. glycolytic rate from technical replicates in the representative experiment shown in I. *P* value of unpaired two-tailed Student's *t*-test.

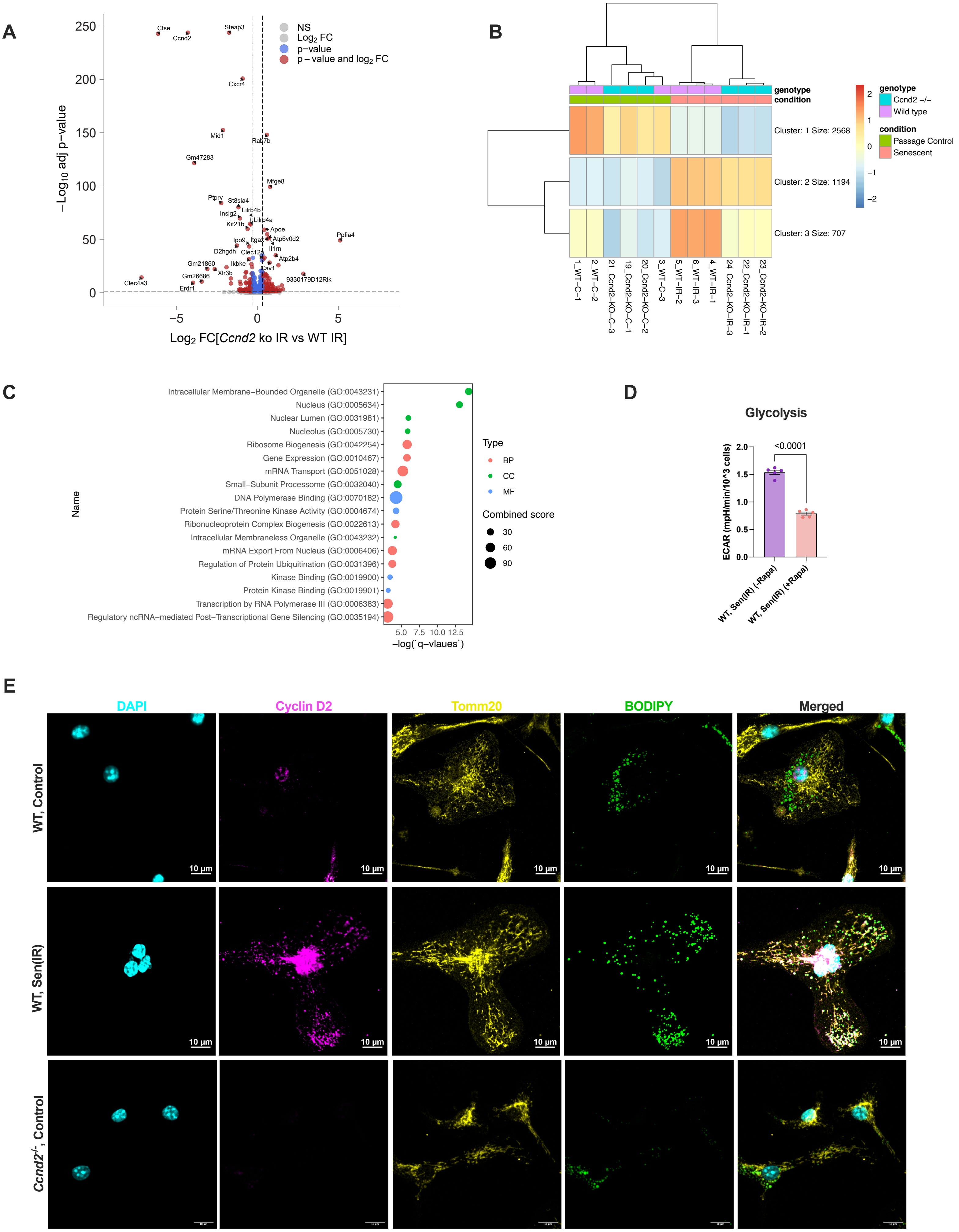


**Supplementary Figure 5. Validation of Cyclin D2 immunofluorescence specificity in *Ccnd2*$^{-/-}$ macrophages.**

**Supplementary Figure 5. Transcriptomic analysis and validation of Cyclin D2-dependent phenotypes.**

(A) Volcano plot showing differentially expressed genes in *Ccnd2*$^{-/-}$ Sen(IR) macrophages compared with WT Sen(IR) macrophages. (B) Heatmap showing k-means clustering of differentially expressed genes into three clusters, with hierarchical clustering of samples based on gene expression profiles. (C) Gene Ontology (GO) enrichment analysis of differentially expressed genes, showing enriched biological process (BP), cellular component (CC), and molecular function (MF) terms. (D) Mean ± s.e.m. glycolytic rate from technical replicates in WT Sen(IR) macrophages treated with vehicle (DMSO) or 50 nM rapamycin for 24h. *P* value from unpaired two-tailed Student's *t*-test. I Representative immunofluorescence images of WT control, WT Sen(IR), and *Ccnd2*$^{-/-}$ control macrophages stained for Cyclin D2, the mitochondrial marker TOMM20, and neutral lipids using BODIPY. DAPI was used to visualize nuclei. The absence of Cyclin D2 signal in *Ccnd2*$^{-/-}$ macrophages validates the specificity of the Cyclin D2 immunofluorescence signal. Scale bars, 10 µm.

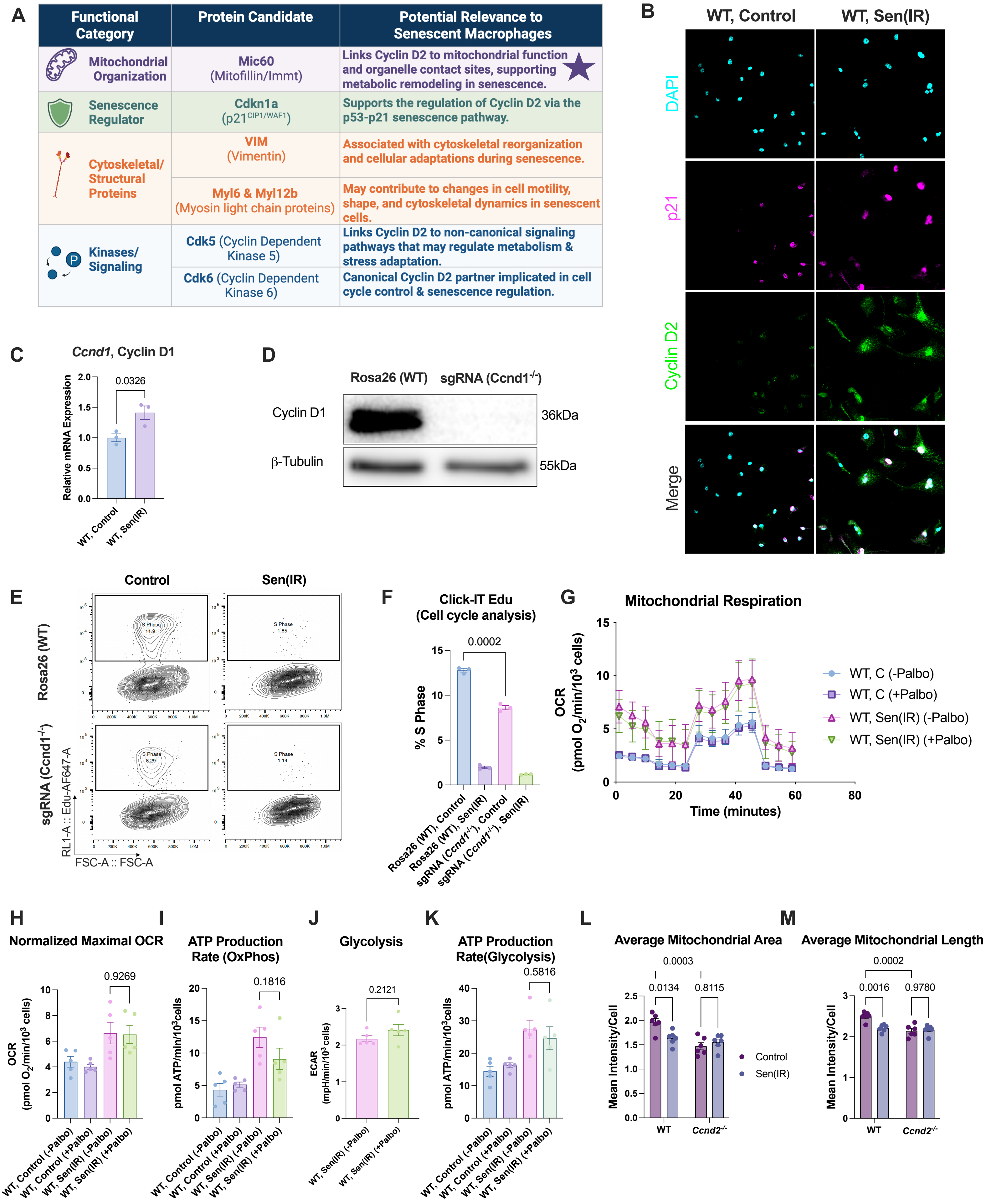


**Supplementary Figure 6. Candidate Cyclin D2-interacting proteins and Cyclin D2-associated mitochondrial phenotypes in senescent macrophages.**

**Supplementary Figure 6. Candidate Cyclin D2-interacting proteins and Cyclin D2-associated mitochondrial phenotypes in senescent macrophages.**

(A) Summary of candidate Cyclin D2-interacting proteins identified by Cyclin D2 immunoprecipitation followed by mass spectrometry, highlighting proteins of interest and their potential relevance to senescent macrophage biology. Candidates include Mic60 (Mitofilin/Immt), the senescence regulator p21 (Cdkn1a), cytoskeletal proteins VIM, MYL6 and MYL12B, and cyclin-dependent kinases CDK5 and CDK6. (B) Representative immunofluorescence images of WT control and Sen(IR) macrophages stained for p21 and Cyclin D2, with DAPI used to visualize nuclei. (C) Relative *Ccnd1* mRNA expression in WT control and Sen(IR) macrophages. (D) Representative immunoblot validating loss of Cyclin D1 protein following CRISPR-mediated *Ccnd1* disruption in Rosa26(WT) and sgRNA(*Ccnd1*$^{-/-}$) macrophages; β-tubulin was used as a loading control. (E) Representative flow cytometry plots of Click-iT EdU incorporation and cell-cycle profiles in Rosa26(WT) and sgRNA(*Ccnd1*$^{-/-}$) macrophages under control and Sen(IR) conditions. (F) Quantification of the percentage of cells in S phase from Click-iT EdU cell-cycle analysis. Data are presented as mean ± s.e.m. *P* value was determined by an unpaired two-tailed *t*-test comparing Rosa26(WT) and sgRNA(*Ccnd1*$^{-/-}$) control macrophages. (G) Mitochondrial respiration measured by Seahorse mitochondrial stress testing in WT control and Sen(IR) macrophages treated with or without 500nM palbociclib (Palbo), 24h. (H) Maximal oxygen consumption rate (OCR) from mitochondrial stress testing. (I) ATP production rate from oxidative phosphorylation (OxPhos). (J) Glycolytic rate measured by extracellular acidification rate (ECAR). (K) ATP production rate from glycolysis. (L) Mean mitochondrial area per cell in control and Sen(IR) WT and *Ccnd2*$^{-/-}$ macrophages. (M) Mean mitochondrial length per cell in control and Sen(IR) WT and *Ccnd2*$^{-/-}$ macrophages. Data are presented as mean ± s.e.m. *P* values for panels C and F-K were determined by the statistical tests indicated in the respective panels. *P* values for panels L and M were determined by two-way ANOVA with Tukey's multiple-comparisons test.

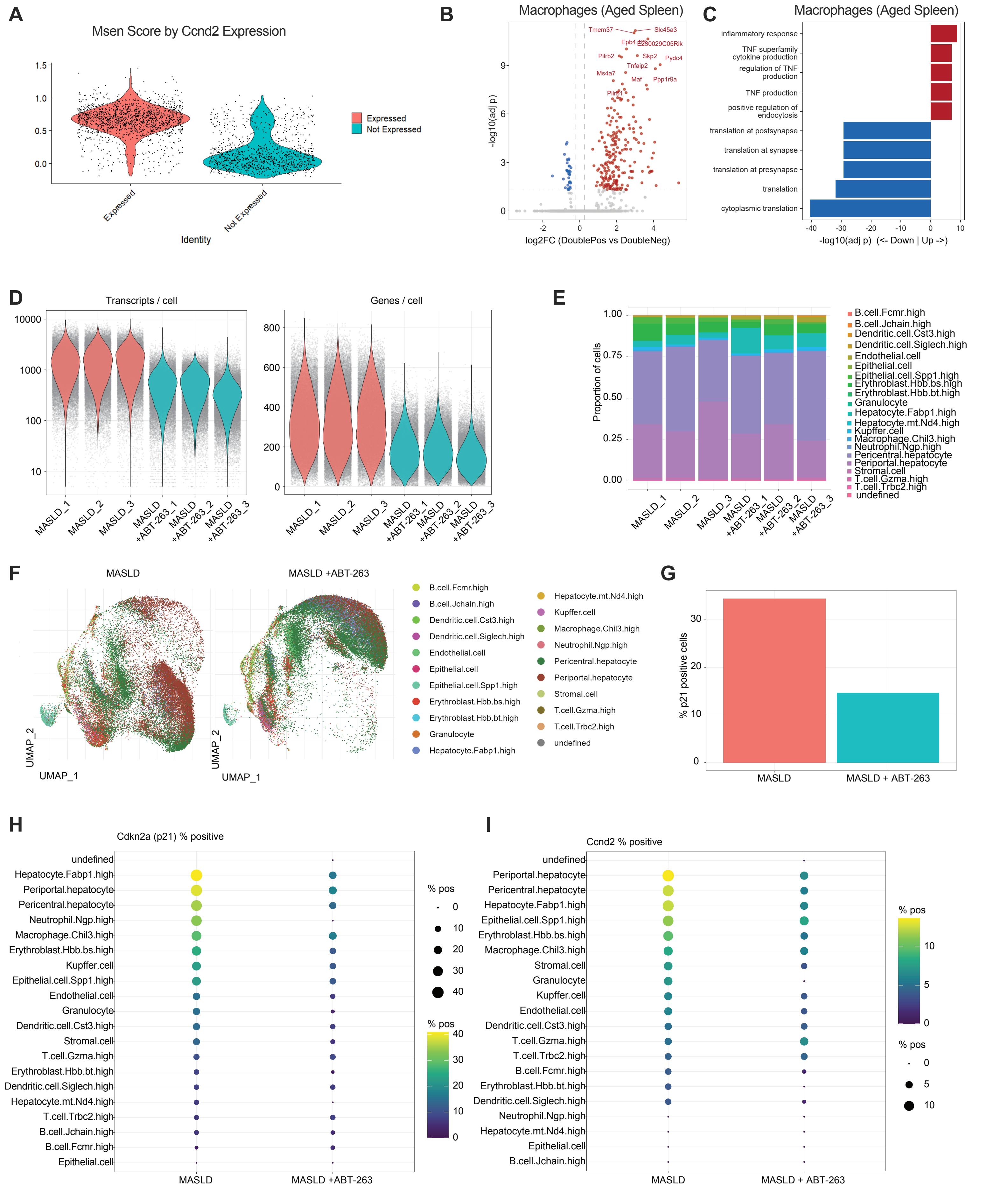


**Supplementary Figure 7. Characterization of Cyclin D2-associated senescence in aging macrophages and spatial transcriptomic profiles of MASLD liver.**

**Supplementary Figure 7. Characterization of Cyclin D2-associated senescence in aging macrophages and spatial transcriptomic profiles of MASLD liver.**

(A) MSen1 senescence scores in *Ccnd2*-expressing and *Ccnd2*-non-expressing Kupffer cells from aged liver, demonstrating enrichment of the senescence phenotype among *Ccnd2*-expressing Kupffer cells. (B) Volcano plot showing differentially expressed genes between Cyclin $D2^{+}p21^{+}$ (double-positive) and Cyclin $D2^{-}p21^{-}$ (double-negative) macrophages from aged spleen, highlighting cytokine- and SASP-associated genes enriched in Cyclin $D2^{+}p21^{+}$ macrophages. (C) Gene ontology analysis of differentially expressed genes between Cyclin $D2^{+}p21^{+}$ and Cyclin $D2^{-}p21^{-}$ macrophages from aged spleen, highlighting enrichment of cytokine production, TNF production, and inflammatory response-associated biological processes. (D) Quality control metrics for spatial transcriptomic datasets from control and senolytic-treated liver sections, showing the number of transcripts and genes detected per cell. (E) Cellular composition of spatial transcriptomic liver sections from control and senolytic-treated mice. (F) UMAP visualization of cell populations identified by spatial transcriptomics in control and senolytic-treated liver sections. (G) Quantification of the percentage of MSen-positive cells in control and senolytic-treated liver sections. (H) Percentage of *Cdkn1a* (p21)-positive cells across identified cell populations in control and senolytic-treated liver sections. (I) Percentage of *Ccnd2*-positive cells across identified cell populations in control and senolytic-treated liver sections.

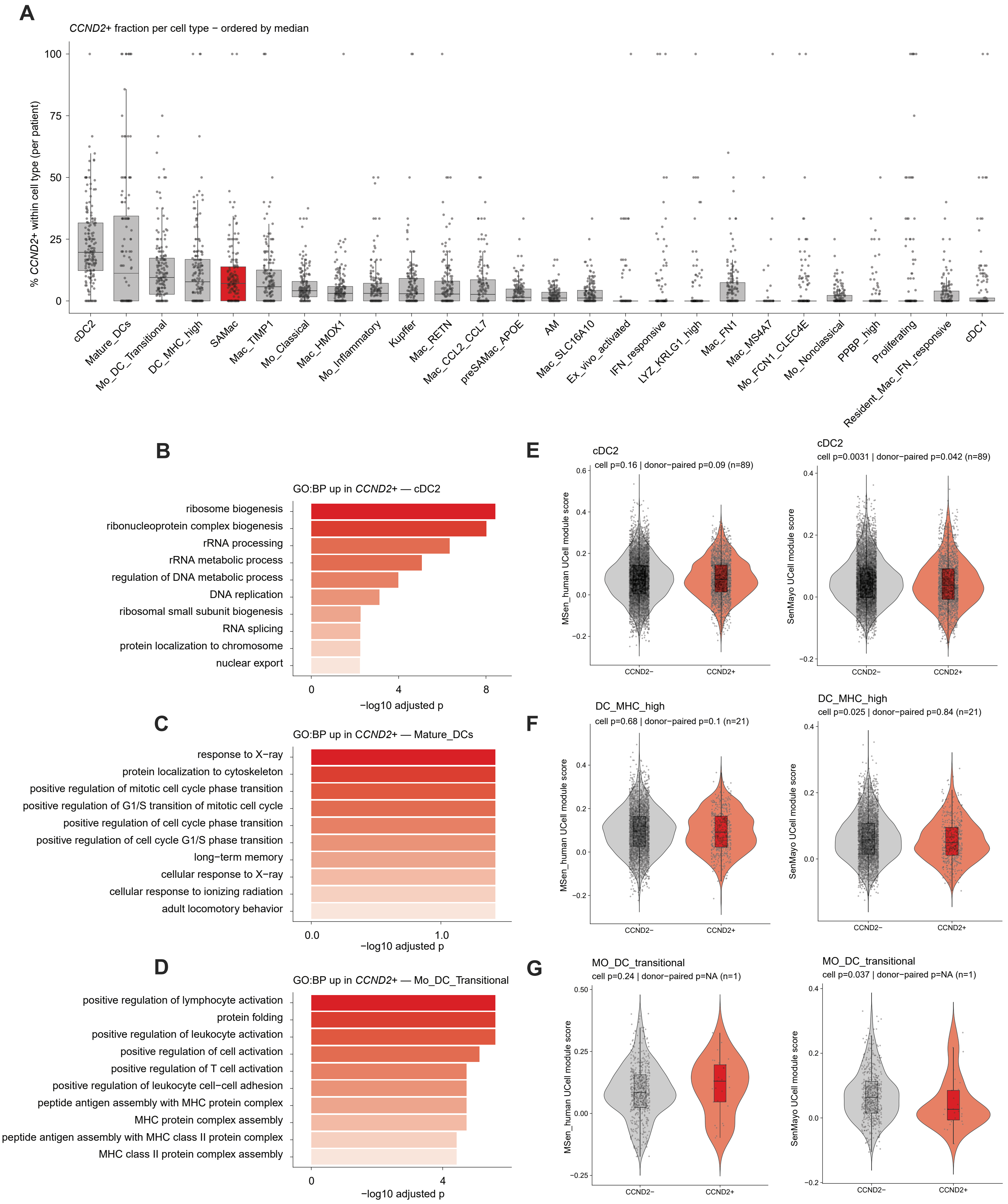


**Supplementary Figure 8. Cyclin D2 is associated with distinct functional programs in a cell type- and context-dependent manner.**

**Supplementary Figure 8. Cyclin D2 is associated with distinct functional programs in a cell type- and context-dependent manner.**

(A) Percentage of *CCND2*-positive cells within identified immune cell populations in human liver, shown for individual patients and ordered by median *CCND2* positivity. (B) Gene Ontology analysis of biological processes enriched among *CCND2*+ versus *CCND2*− cDC2s, highlighting enrichment of RNA processing, ribonucleoprotein complex biogenesis, ribosome biogenesis, DNA replication, and regulation of DNA metabolic processes. (C) Gene Ontology analysis of biological processes enriched among *CCND2*+ versus *CCND2*− mature dendritic cells, highlighting enrichment of cell-cycle-associated processes, including positive regulation of G1/S transition and mitotic cell-cycle progression. (D) Gene Ontology analysis of biological processes enriched among *CCND2*+ versus *CCND2*− transitional monocyte-derived dendritic cells, highlighting enrichment of leukocyte and lymphocyte activation and MHC class II antigen presentation pathways. (E–G) MSen_human and SenMayo UCell module scores in *CCND2*− and *CCND2*+ cDC2s (E), DC_MHC_high cells (F), and transitional monocyte-derived dendritic cells (G). *P* values for cell-level and donor-paired comparisons are shown in the figure.
